# AI-guided design of common light chains to enable manufacturable bispecific antibodies

**DOI:** 10.1101/2025.10.11.681265

**Authors:** Yi Zhou, Brian M. Richardson, Longfei Cong, Tong Li, Roger Shek, Yuxiang Lang, Ziwei Pang, Fernando Garces, Jarrod B. French, Per Greisen

## Abstract

Bispecific antibodies (BsAbs) offer therapeutic advantages but face manufacturing bottlenecks from light chain mispairing, which can generate a substantial fraction of incorrect products and increase manufacturing complexity and cost of goods. Common light chains (cLC) eliminate mispairing, yet existing approaches require screening thousands of variants per target. We present an AI-driven framework that computationally designs cLCs through structure-guided pairing of non-cognate VH-VL interfaces, reducing experimental screening by three orders of magnitude. The platform successfully engineers therapeutic antibodies lacking experimental structures, expanding applicability beyond crystallographic databases. Among 10 therapeutic targets, we successfully generated designs for 7 targets, comprising 55 unique BsAb pairs. Of these, 43.6% (24/55) were successfully as BsAb cLCs. Three bispecific antibodies reached production-ready specifications: >90% purity and 1.6-1.8 g/L titers. This platform democratizes bispecific antibody development, expanding access beyond well-resourced programs.

## Introduction

Manufacturing challenges limit the clinical adoption of bispecific antibodies (BsAbs) despite their proven therapeutic superiority. In IgG-like BsAbs, heavy chain (HC) homodimerization (VH–VH/CH3–CH3) and light chain (LC) mispairing are the two primary assembly errors; knobs-into-holes (KiH) efficiently addresses the former, while LC mispairing remains the dominant bottleneck. Random chain pairing during production can yield a complex mixture of up to ten different molecular species, with only minor amounts of the correctly assembled antibody, creating a significant manufacturing bottleneck^1-4^. With >150 BsAbs in development and a projected $163B market by 2032^2^, efficient chain-pairing solutions are a critical priority.

The fundamental problem that impairs efficient BsAb production stems from promiscuous pairing between HC/LC during co-expression. In IgG-like BsAbs, two HC and two LCs must assemble correctly, but mispairing is frequent because VH–VL interfaces have comparable binding energies^1,11^. While KiH^6^ technology effectively prevents heavy chain homodimerization with >92% efficiency, LC mispairing remains the primary manufacturing bottleneck. Other approaches including chain pairing mutations (CPMs)^7^, CrossMAb^8^, and DuoBody^9^ address LC pairing but typically impose format-specific engineering constraints (e.g., Fc/CH3 mutations, domain swaps, or pairing-orthogonal interfaces) that are not universally portable across antibody backbones^10^. A common light chain (cLC) offers a general solution by eliminating mispairing.

In vivo cLC discovery platforms using genetically engineered mice encode a single LC across the entire B-cell repertoire, severely limiting binder diversity against any given target. Chugai validated the FVIII-mimetic fIXa/fX bispecific concept preclinically with emicizumab^12^ and subsequently reported a multidimensional optimization and manufacturability program that delivered the clinical molecule^13^. Rather than attempting exhaustive combinatorics, we use structure-guided cLC design to pre-prioritize a limited number of VH–VL pairs for experimental testing. We define efficiency operationally as “fewer constructs requiring post-design experimental validation,” acknowledging that cross-program comparisons depend on format, assay funnels, and organizational context. This route relied on a common LC architecture to mitigate LC mispairing^12,13^. The central challenge is to engineer LCs that maintain affinity with multiple non-cognate HCs while preserving stability. This is complicated by the VH-VL interface involving 15– 20 residues, where optimization for one partner can compromise another^13,61^. Current cLC methods achieve limited success^62^ and often require 10-20 experimental validations^15^, highlighting the need for more predictive approaches (Supplementary Note 1).

Recent advances in protein structure prediction and design create opportunities for computational cLC engineering. AlphaFold-based methods achieve near-experimental accuracy^18^, while machine learning enables rational protein design^19^. The Structural Antibody Database (SAbDab) ^20^ contains >10000 antibody structures, providing unprecedented training data for understanding VH-VL pairing preferences. Additionally, molecular dynamics (MD) simulations can predict stability and binding of designed interfaces^21^.

Here we present an AI-driven framework that reduces BsAb screening burden from thousands of variants to only a few per target. Our integrated computational platform successfully engineers cLC for 7/10 of therapeutic targets tested, requiring only limited experimental validations compared to thousands in empirical approaches, with validated designs achieving >80% purity at 1.6–1.8 g/L expression and able to generate 51 BsAbs against 24 diverse sets of targets. The platform supports both κ and λ chains and reveals IGKV1-39 enrichment linked to broad VH–VL compatibility^22^. This establishes a generalizable, de novo route to manufacturable BsAbs, transforming empirical discovery into predictable design.

## Methods

This section provides an overview of key methods; comprehensive protocols, parameters, and computational requirements are detailed in Supplementary Methods with cross-references provided throughout.

### High-Resolution Dataset and Structure Modeling

We compiled 3,639 high-quality antibody-antigen complexes from SAbDab^20^ (resolution <3.0 Å). Structure prediction employed Chai-1 v0.6.1^23^ with custom VH-antigen restraints, generating 5 models per VH-VL combination. Models were validated through 10 independent 6 ns MD simulations using GROMACS v2024.5^52^ with AMBER FF14SB^46^. Model sidechain repacking and interface analysis were performed using Rosetta v3.13^**35**^. Detailed protocols including restraint specifications, simulation parameters, and validation criteria are provided in Supplementary Methods Sections 1– 3.

### Candidate Selection Pipeline

Models passed multi-tier filtering: Chai-1 confidence (pTM >0.7, ipTM >0.7), structural deviation (RMSD <2.0 Å), Rosetta metrics (sc_value >0.6, packstat >0.65), and MD stability (≥70% interaction preservation). MM/GBSA binding energies were calculated for final ranking. Complete filtering parameters, scoring functions, and selection algorithms are detailed in Supplementary Methods Sections 4-5 and Supplementary Fig. S1.

### Germline frequency analysis

Germline V-gene frequencies were computed for (i) therapeutic antibody starting points, (ii) the computational VL sequence pool, and (iii) experimentally validated cLCs using identical parsing rules. Because the starting sets are derived from PDB/Thera-SAbDab and thus reflect known repertoire and crystallization biases, we report germline distributions descriptively and do not ascribe functional superiority to any germline without de-biasing (e.g., template-matched resampling) or prospective validation on unbiased libraries. All antibody sequence numbering was performed using ANARCI^31^ with the Chothia scheme^30^.

### Experimental Validation

Antibodies were expressed in ExpiCHO-S cells using automated transfection (Hamilton STAR). Bispecific constructs used KiH mutations^6^ with differential tags. Binding was measured by biolayer interferometry (Octet RED384) and surface plasmon resonance (BIAcore 8K), purity by SEC-HPLC and SDS-PAGE. Full experimental protocols including expression conditions, purification procedures, and analytical methods are described in Supplementary Methods Sections 6–8 and Supplementary Fig. S3.

### Code and Data Availability

Complete computational pipelines, analysis scripts, and configuration files are available at https://github.com/biomap-research/CommonLightChain. Arpeggia software is publicly available at https://github.com/y1zhou/arpeggia/^24^. All structural models and simulation trajectories will be deposited in appropriate public repositories upon publication, with detailed environment files, example scripts, and parameter configurations provided for full reproducibility.

## Results

We first outline the computational workflow and success metrics, then quantify baseline performance by target, evaluate structure-guided light-chain rescue, and conclude with manufacturability outcomes.

We report a 10-target validation cohort with a target-level success of 7/10, supported by single-arm testing of 204 constructs (85 baseline + 119 post-optimization) across a broader exploratory set; 183 BsAb pairs were assembled from these data (Table 1).

**Table 1.** Hierarchy of success metrics (single-arm testing) with constructs tested and BsAb assembly counts. Metrics are reported at three levels—target, strategy, and construct—to avoid double counting and separate target-level outcomes from construct-level screening/rescue rates. “Success” at the construct level is defined as 1. Purity by SEC Main >80% to avoid aggregate dominated false positive binding, 2. observable binding (K_D_ < 1uM). Notes: This table summarizes outcomes using single-arm testing as the primary assay context. Percentages are paired with n/N. Construct-level “success” requires meeting the purity and affinity criterion observable during the in vitro binding experiment. a) Supplementary Table S2 b) Supplementary Table S4 † 183 denotes the number of unique BsAb pair combinations possible from validated single-arm constructs (See Supplementary Table S3);

| **Metric (label)** | **What it measures (plain-English)** | **Numerator / Denominator** | **Reported value** | **Level of analysis** | **Constructs tested (single-arm)** |
| --- | --- | --- | --- | --- | --- |
| **Target-level success** | Number of unique targets for which $\geq 1$ engineered cLC produced a successful binder (platform validation) | 7 / 10 unique targets | 70% | Target | 85<sup>a)</sup> (baseline screen) + 119<sup>b)</sup> (post-optimization) = 204 tested; 183† bsAb pairs assembled from these data |
| **Strategy-level success – Tier 1** | Targets successful when retaining one native VL as the cLC | 3 / 5 targets | 60% | Target (by strategy) | 49<sup>a)</sup> (single-arm, Tier 1) |
| **Strategy-level success – Tier 2** | Targets successful when identifying new VLs for both arms (true de novo pairing) | 3 / 6 targets | 50% | Target (by strategy) | 36<sup>a)</sup> (single-arm, Tier 2) |
| **Construct-level baseline binding success** | Fraction of constructs that bound before structure-guided rescue/optimization | 33 / 85 | 38.8% | Construct | 85<sup>a)</sup> (baseline single-arm set) |
| **Construct-level post-optimization success** | Fraction of constructs maintaining or restored to binding after<br>structure-guided LC<br>optimization<br>(pre-specified threshold) | 80 / 119 constructs | 67.2% | Construct | 119<sup>b)</sup> (post-optimization single-arm set) |

### Computational platform design and validation

BsAbs face manufacturing challenges from chain mispairing that generate up to ten different molecular species^25^. Our platform reduces >2,300 VL sequences from the PDB to few optimized variants per target. In our validation panel, LC-mispairing was observed in 7/10 products and HC-mispairing in 6/10, consistent with—but not identical to—the theoretical maximum species count. At the target level, success was observed for 7/10 unique targets (1 antigen appears in both Tier 1 and Tier 2; see Supplementary Table S1). Targets were selected to represent diverse therapeutic areas (oncology: ERBB2, VEGFA; immunology: CD19, CD38, CD47; inflammation: TSLP, TNR9 – see Supplementary Table S1), different epitope classes, providing a comprehensive test of platform generalizability. Validation was structured in two strategies at the construct level rather than by target: Tier 1 retained one native VL as the cLC and displayed a success rate of 3 out of 5 targets, while Tier 2 required de novo identification of new VLs for both arms and showed a success rate of 3 out of 6 targets (1 target was the same between Tier1 and Tier2 hence 10 unique targets – see Supplementary Table S1).

The platform integrates five stages: VH–VL interface analysis, restraint-guided structure modeling, MD validation, mutation optimization, and candidate selection (Fig. 1a; comprehensive methods in Supplementary Methods Sections 1-5). Starting from >2,300 VL sequences (574 λ, 1,804 κ) from the PDB, filtering yields ∼1,000 compatible sequences, 100–200 structural models, 30–50 dynamically stable designs, and few final variants per target (Fig. 1b).

**Figure 1.**
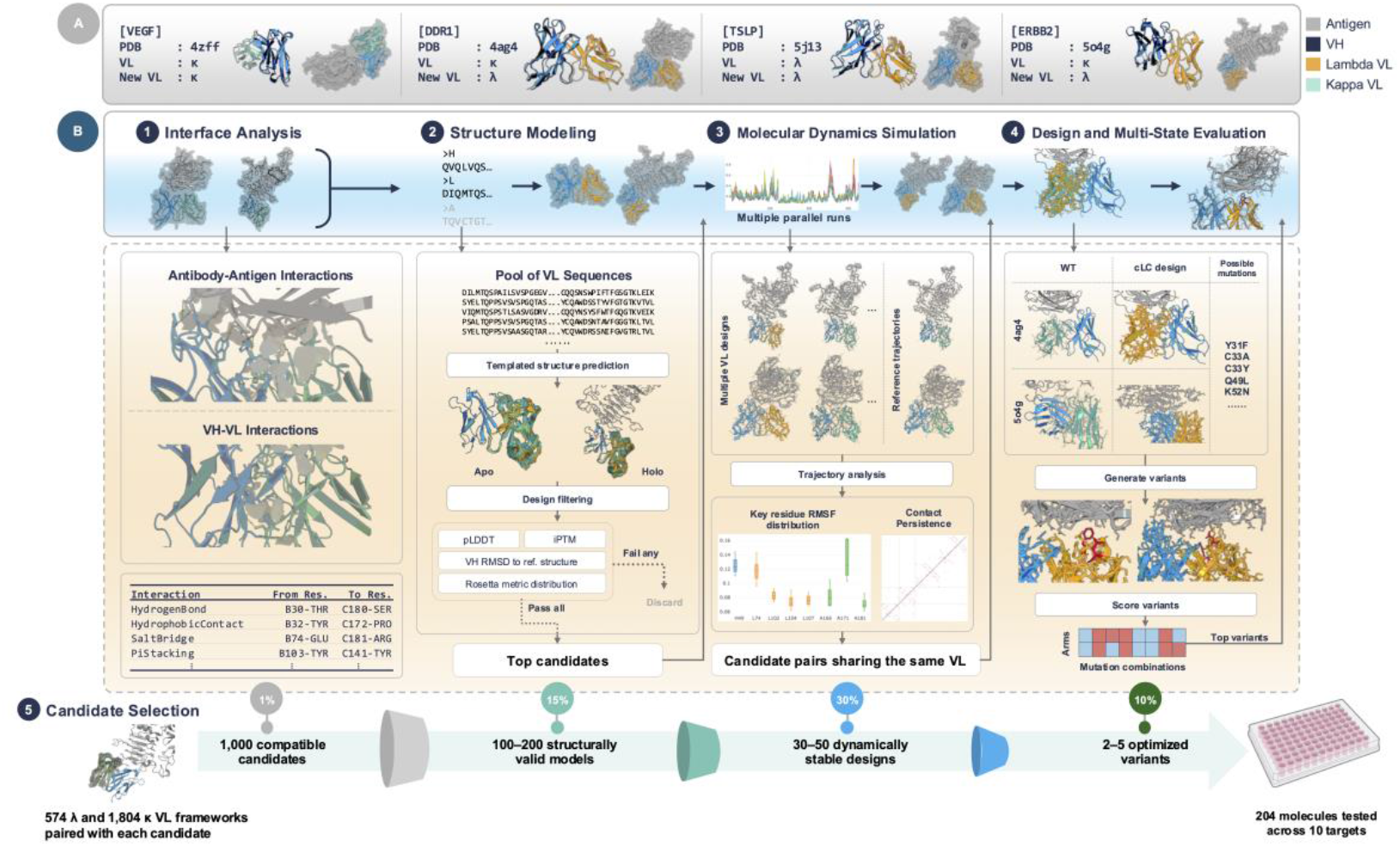
Structure-based computational recipe for cLC design. **(A)** Target diversity analysis showing example antibody-antigen complexes with kappa (green) and lambda (yellow) variable LCs, while HCs remain consistently colored (blue). Both apo (unbound) and holo (antigen-bound) conformations are analyzed with PDB identifiers provided. **(B)** Computational workflow: (B1) Antibody-antigen interface analysis with VL extraction and database searching; (B2) Structure prediction and statistical validation of new LC models in both apo- and holo-states; (B3) MD simulation comparing multiple VL variants, with B-factor coloring and motion arrows illustrating flexibility patterns across different designs; (B4) Rosetta design optimization comparing wild-type versus common LC mutations with multi-state evaluation; (B5) Final candidate selection using integrated computational metrics. Recipe boxes detail the input, process, and output for each step, providing a complete methodology for generating cLC designs. Inset: empirical library sizes reported in prior workflows e.g. NXT007 (second-gen emicizumab) ^16^ versus candidates tested in this work.

Structure modeling employed Chai-1^26^ with VH-antigen restraints followed by ten 6 ns MD simulations per candidate and interaction analysis using Arpeggia^27^. Structural predictions aligned closely with experimental structures, achieving an RMSD of 0.6 Å (Supplementary Fig. S3). Restraints proved critical: unconstrained models often mispositioned the LCs (Supplementary Fig. S2; see Supplementary Methods Section 2 for restraint implementation details). Practically, this meant advancing only pre-prioritized constructs to the bench for each target, concentrating assays on the most plausible designs rather than on large, brute-force libraries.

### Target-specific performance with unmutated cLCs

We tested 85 constructs across 10 targets—substantially fewer than the thousands required by empirical screens (Supplementary Table S3). Performance varied by target: ERBB2 showed 45.5% success (15/33 constructs), while CD38 was refractory (0/19; Table 2). This target-specific variation revealed key determinants of pairing compatibility. (substantially fewer than typical empirical screens; see ref. 12). Targets with strong performance included ERBB2 and VEGFA; for example, ERBB2 appeared in 33 constructs with an overall cLC success of 45.5% (15/33; 95% CI 29.8–62.0), with performance varying by partner (Fig. 2; Supplementary Table S3), consistent with high partner compatibility. By contrast, CD38 was refractory (0/19; 0.0%, 95% CI 0.0– 16.8%). In our structural analysis (Fig. 4 / Supplementary Table S7), CD38 interfaces exhibit increased polar contacts, particularly involving CDR-H3, which likely elevates the requirement for complementary VL polar interactions and increases sensitivity to VL substitutions. Insufficient VL support likely destabilizes CDR-H3, reducing binding (Supplementary Table S7).

**Table 2.** Target-specific performance summary of computational cLC engineering across diverse therapeutic targets. Native LC indicates the original LC type (κ=kappa, λ=lambda), while New LC shows the engineered cLC types tested. Best cLC / Native K_D_ Fold represents the most successful candidate’s affinity change relative to the original antibody without cLC. Key challenges highlight the primary engineering obstacles encountered for each target, ranging from structural complexity to epitope sensitivity. a) NA indicates constructs not advanced to K_D_ quantification under the predefined thresholds.

| Target | Native LC | New LC | Candidates Tested (N=85) | $K_D$ LC (M) | $K_D$ Native LC (M) | Best cLC / Native $K_D$ Fold | Key Challenges |
| --- | --- | --- | --- | --- | --- | --- | --- |
| VEGFA | $\kappa$ | $\kappa(18)$ | 18 | 3.64e-10 | 1.73e-10 | 2.1x | Extensive VH-VL interfaces; mutations improved hydrophobic contacts |
| ERBB2 | $\kappa$ | $\kappa(11)/\lambda 1(1)$ | 12 | 7.013e-10 | 1.325e-09 | 0.53x | Complex conformational rearrangements; epitope sensitivity |
| CD19 | $\kappa$ | $\kappa(9)$ | 9 | 3.28e-08 | 2.63e-10 | 124x | Cell surface marker; mixed $\kappa/\lambda$ approaches tested |
| DDR1 | $\kappa$ | $\kappa(9)/\lambda(2)$ | 11 | 6.45e-10 | 2.058e-09 | 0.31x | Challenging $\kappa \rightarrow \lambda$ conversion; sensitive epitope interfaces |
| TSLP | $\lambda$ | $\kappa(13)/\lambda(2)$ | 15 | 1.12e-09 | 7.29e-12 | 153x | $\lambda \rightarrow \lambda$ substitution; maintained lambda-specific features |
| CD38 | $\kappa$ | $\kappa(8)$ | 8 | NA<sup>a</sup> | NA | NA | NA |
| CD47 | $\kappa$ | $\kappa(4)$ | 4 | NA | NA | NA | NA |
| Z13-IL22-2 | $\kappa$ | $\kappa(3)/\lambda(2)$ | 5 | 6.3e-07 | 1.07e-08 | 58x | NA |
| HAVR2 | $\kappa$ | $\kappa(1)$ | 1 | NA | NA | NA | NA |
| **TNR9** | $\kappa$ | $\kappa(2)$ | 2 | 2.28e-08 | 3.603e-09 | 6.3x | NA |

**Figure 2.**
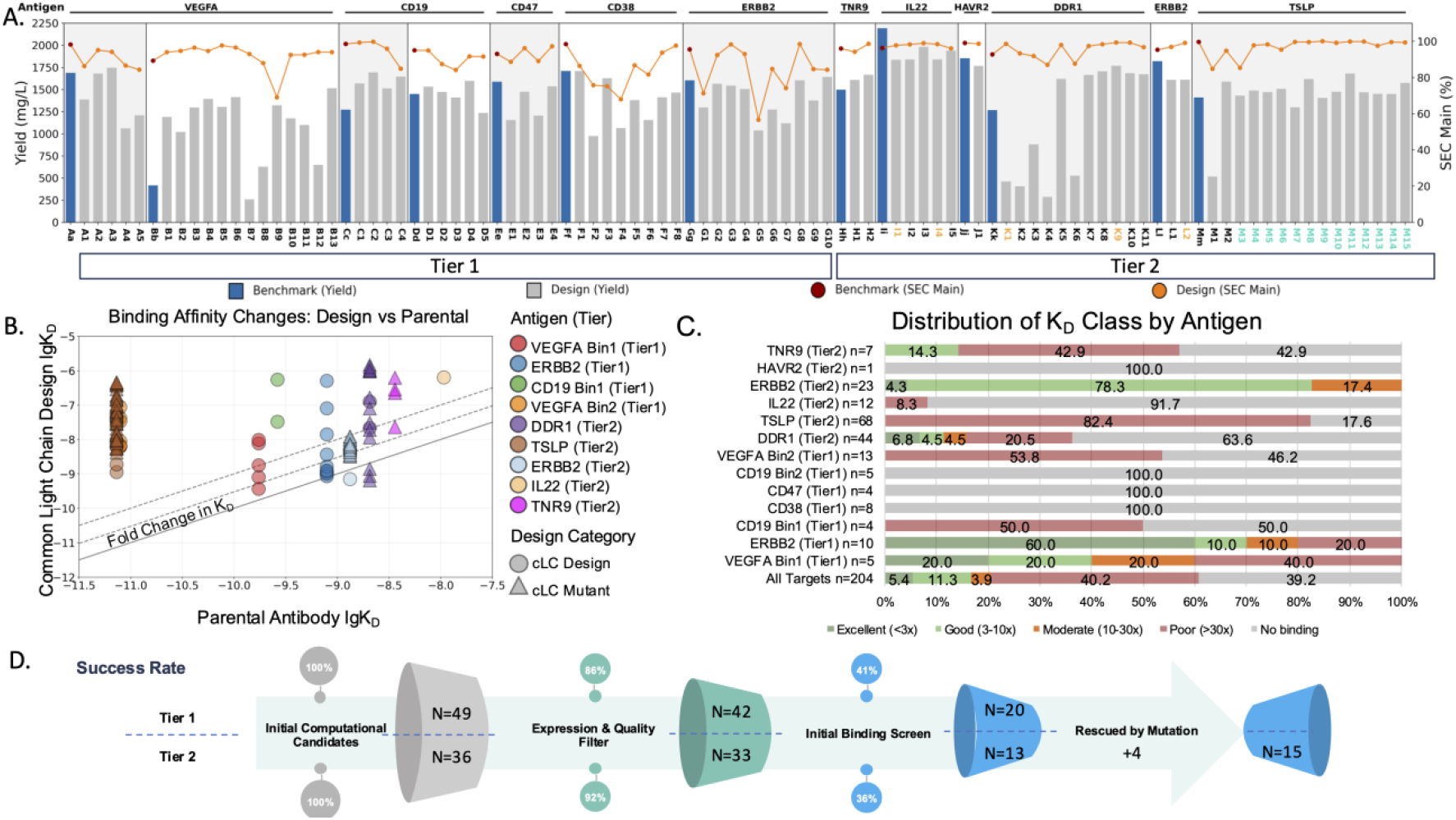
Comprehensive validation of computational cLC engineering. **(A) Target-specific performance metrics**. Expression yields (mg/L, bars) for tested cLC designs (gray) versus parental antibodies (blue). Orange line shows SEC main peak percentage. Red dots indicate SEC main peak percentage for parental antibodies. Left: Tier 1 targets (VEGFA, CD19, CD47, etc.); Right: Tier 2 targets (TNR9, IL2, etc.). **(B) Binding affinity changes: Design vs Parental**. Comparison of K_D_ values between engineered cLCs and parental antibodies on log scale. Points below diagonal indicate improved binding. Solid line represents 3-fold therapeutic threshold. Dashed line represents 5-fold and 10-fold threshold. Symbols: circles (computational designs), triangles (cLC mutant). **(C) Distribution of binding outcomes**. Proportion of designs achieving excellent (<3-fold, dark green), good (3- to 10-fold, green), moderate (10- to 30-fold, orange), poor (>30-fold, dark red), or no binding(gray) relative to parental antibodies across all tested targets. **(D) Screening cascade comparison**. Progressive candidate enrichment through computational and experimental filtering for Tier 1 (top) and Tier 2 (bottom) validation cohorts. Percentages indicate retention rates at each stage.

### Germline convergence reveals structural determinants of promiscuous pairing

To identify sequence features enabling broad VH compatibility, we analyzed germline gene usage across validated cLCs, PDB template structures, and therapeutic repertoires (Thera-SAbDab). Successful designs showed apparentenrichment of IGKV1-39 (11% in templates -> 30% in validated cLC; Fig. 3). Because published repertoire frequencies vary by cohort ^22,59^, sampling depth, and annotation pipeline, we do not assert a specific baseline percentage here (Supplementary Fig. S3). Structural analysis revealed that its CDR-L3 loop contains smaller amino acid residues that create more physical space to accommodate a wide variety of CDR-H3 conformations, a key feature that reduces steric hindrance and enables its broad pairing compatibility^74^. These structural features enable accommodation of diverse VH CDR-H3 conformations without compromising VH-VL interface stability. J-gene usage remained comparatively stable across datasets (IGKJ1: 40–60%), confirming that V-gene—not J-gene— selection is the primary determinant of compatibility (Fig. 3; Supplementary Table S8).

**Figure 3.**
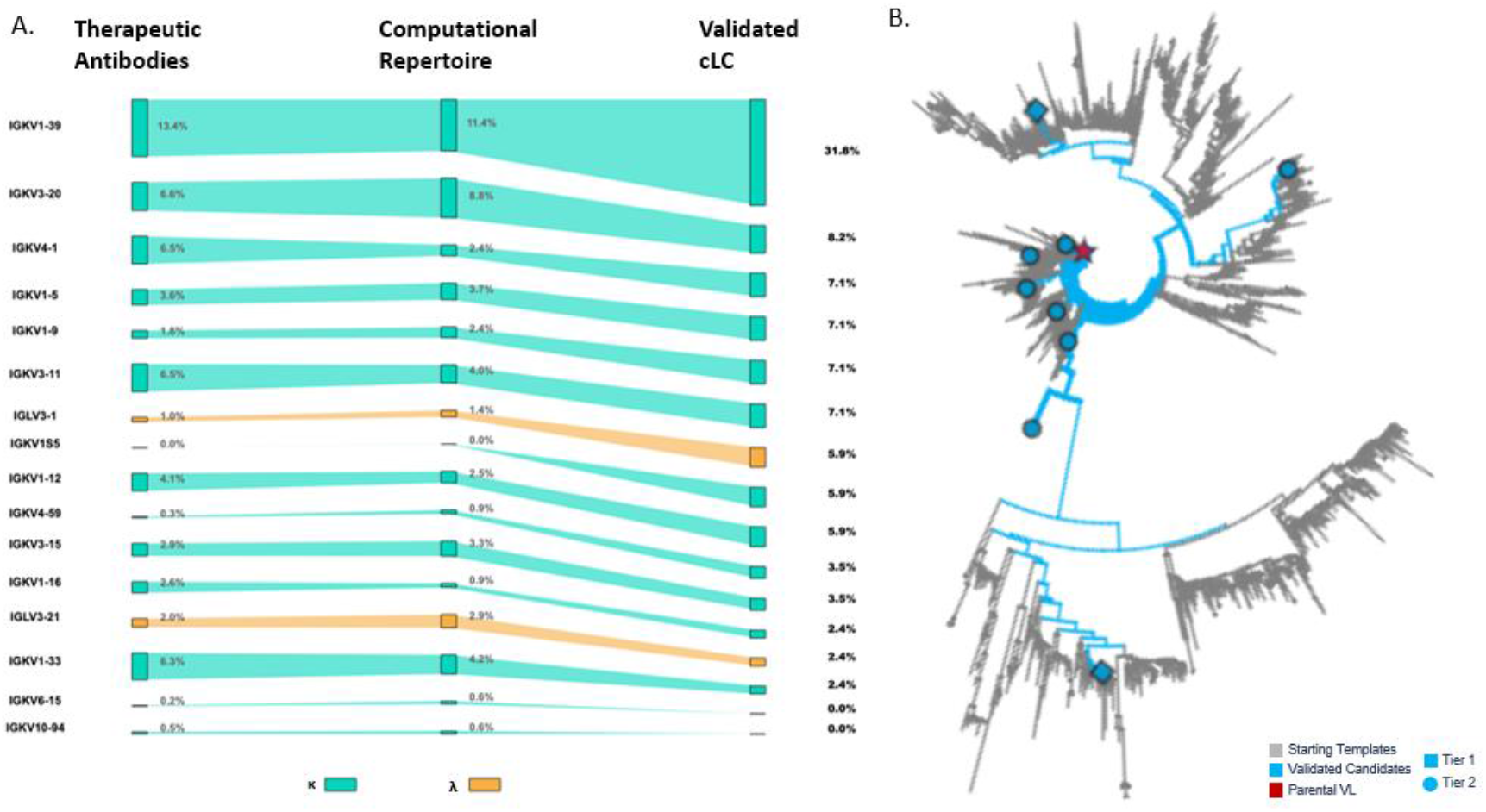
Germline gene usage analysis of successful cLC designs. **(A)** Germline V gene representation at three key stages: therapeutic antibody starting points (left bars), computational VL sequence pool available for in silico screening (middle bars) and experimentally validated cLCs for target pairs (right bars). Each row represents a different V gene family. Bar widths indicate the number of sequences at each stage. Cyan: kappa LCs; orange: lambda LCs. The visualization reveals how computational screening samples broadly across V gene diversity in the pool (middle) but converges on specific germline families—particularly IGKV1-39—in successful validated designs (right), despite relatively even representation in both therapeutic antibodies and the starting computational pool. Enrichment is reported descriptively; no functional advantage is inferred without de-biasing analyses (see Methods). (**B**)Phylogenetic tree (bottom right) shows the parental VL of an ERBB2 antibody (red) and successfully paired designed VLs (blue) distributed across sequence clades.

Successful cLCs nevertheless spanned multiple κ and λ germline families (Fig. 3A-B), demonstrating platform generalizability beyond IGKV1-39 scaffolds. The platform enabled mouse-to-human germline transitions in 17 tested cases (Supplementary Fig. S9), validating applicability to humanize light chains during the design^37^. While PDB and Thera-SAbDab over-represent crystallizable antibodies (IGKV1-39 ∼ 13% in therapeutic repertoires), we treat the observed ∼30% incidence in validated cLCs as a working hypothesis rather than functional evidence, given known database biases. We do not infer intrinsic superiority of IGKV1-39 from these data and defer causal interpretation to future tests on unbiased libraries and prospective, PDB-independent designs.

### Structure-guided LC optimization

cLC rescue is a dual-constraint problem: mutations must sustain productive contacts in both VH–VL–antigen contexts. We extended the design pipeline by adding in silico mutagenesis that scores VH–VL compatibility and VL–antigen contacts for both arms prior to synthesis.

An initial cLC design yielded 85 candidates with binding measurements: 33 with measurable binding and 52 without (Supplementary Table S2; definition in Methods). To improve the set of binder and non-binders, we performed structure-guided optimization across CDR-L1, CDR-L2, and CDR-L3, prioritizing positions from interface-complementarity analyses that incorporated prior information from the cognate VL’s chemical complementarity to each VH–antigen complex. Using Rosetta, we designed 56 sequence-distinct LC substitution patterns spanning single-through quadruple-site edits. These included substitutions and deletions to also address developability liabilities (e.g., Cys→Ser/Ala for free cysteines). Applied to the initial candidates across targets, this yielded 119 expressed VH–VL constructs (with individual LC designs tested across multiple VH contexts) for binding assays (Supplementary Table S4).

Binding was observed for 67.2% of constructs (80/119). Measurable binding was defined as an SPR/BLI response above baseline sufficient to estimate K_D_, with a resolution of ∼1 µM (Methods). Among binding-competent constructs, outcome classes were: rescued binders—transition from no binding to binding 42.9% (24/56); improved binders - K_D_ improved by ≥3× relative to the initial seed 11.1% (7/63); maintained binders -K_D_ change <10× and not improved by >3× 65.1% (41/63); and weakened binders -seed binding decreased by >10× but still detectable 12.7% (8/63) and 11.1% (7/63) losing binding. Subgroup denominators reflect the relevant seed cohort for each outcome class. Notably, 76.2% (48/63) of constructs with initial binding either maintained or improved affinity after rescue, indicating that structure-guided modifications preserved functional compatibility in the majority of cases. Rescue rates varied by target (e.g., TSLP and ERBB2 were permissive, whereas DDR1 and Z13-IL22-2 were refractory; Fig. 4A and Supplementary Fig. S4). Combinatorial designs outperformed single-site edits (74.2% vs 59.6%; Supplementary Table S4), consistent with the need to satisfy constraints across both interfaces.

**Figure 4.**
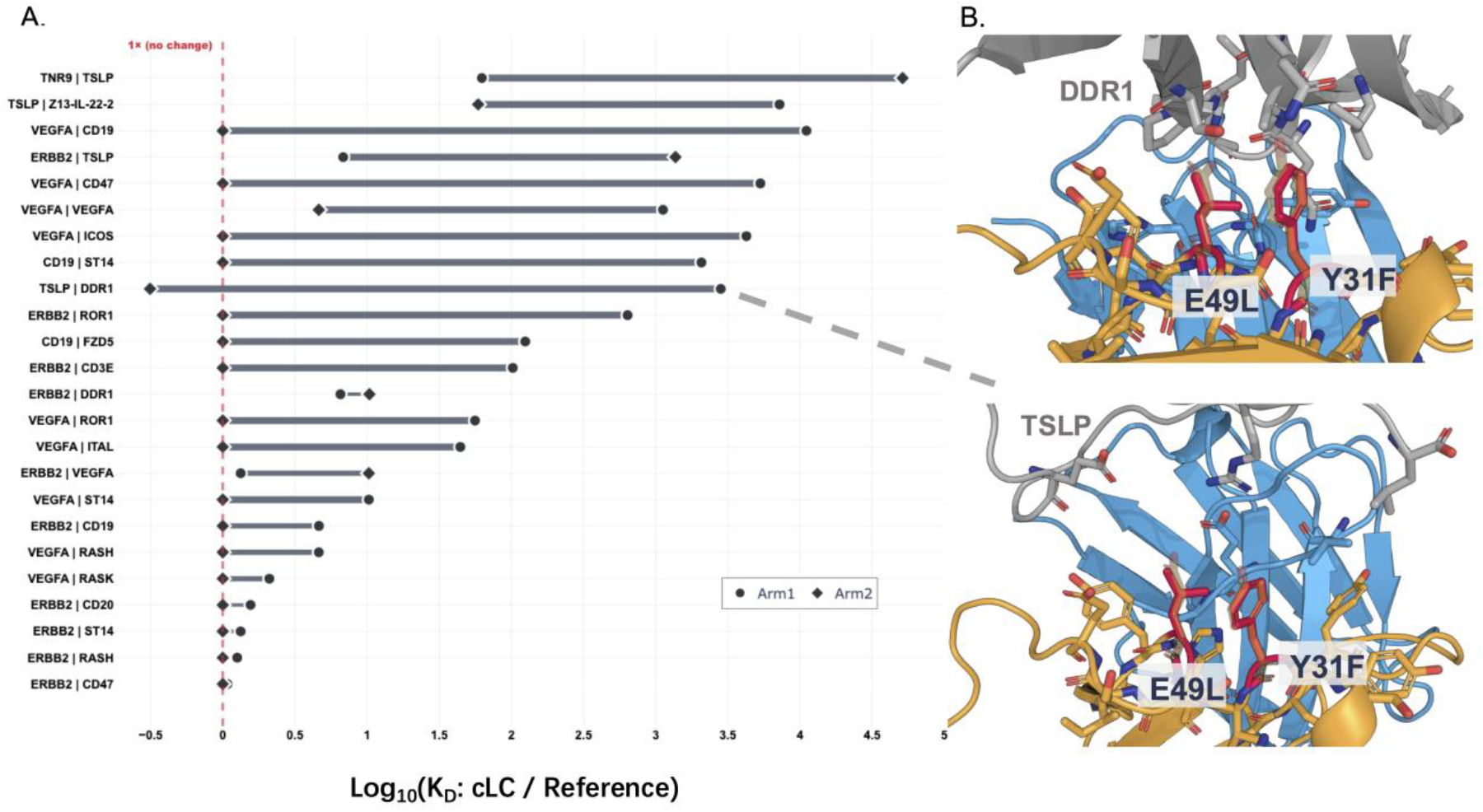
Bispecific rescue outcomes and representative sites. **(A)** Fold-change in affinity relative to the reference LC for each bsAb pair (Arm1, Arm2). Vertical line at **1×** indicates parity; values to the left indicate improved affinity (lower Kd). Rows are ordered by the larger fold-change across arms. For the TSLP/DDR1 pair, for instance, the cLC improved affinity for the TSLP arm (leftward point) while maintaining near-parental affinity for the DDR1 arm (point near 0). **(B)**Representative cLC mutations in **DDR1** and **TSLP** constructs. The cLC (orange) interfaces with VH (blue) and antigen (gray); mutated residues (red sticks) are labeled (e.g., **Y31F, E49L**). Positions are located in **L1** and **L2** loops. See Supplementary Table S3 for all BsAbs tested and Supplementary Table S5 for all mutations generated.

These results underscore both the promise and limits of computational cLC engineering: explicit dual-arm scoring improves recovery of non-cognate VH–VL pairs, yet discrepancies likely reflect effects on expression, stability, and fine-scale interface energetics not fully captured by current scoring functions^61^. We therefore assembled the optimized LCs into bsAbs and asked whether functional output is limited by the weaker arm’s affinity (Table 1).

### Rescuing weak or non-binding arms and restoring BsAb function

We focused on the limiting arm for each target pair and tested whether arm-specific rescue mutations achieved the dual requirement of (i) restoring or improving affinity in the mono assay and (ii) maintaining compatibility in the bispecific format. Fig. 4A shows pair-level outcomes by plotting, for each BsAb, the fold change in affinity of Arm1 and Arm2 relative to their respective reference LCs (1× = parity). Ordering pairs by the largest arm-wise improvement reveals three consistent patterns that align with the preceding “Multi-interface optimization” subsection and Table 2.

First, asymmetric rescues are common. Many pairs show a clear improvement in one arm while the partner arm remains near parity (Supplementary Table S5). This mirrors the earlier finding that single-arm rescue was the most tractable strategy overall, outperforming attempts to rescue two nonfunctional arms simultaneously and tracking with the general advantage of combinatorial over single-site edits (Table 3). Second, outcomes are target-dependent at the pair level (Fig. 4A; Table 3). Pairs containing TSLP or ERBB2 occupy the upper portion of the plot (larger leftward shifts from 1×), consistent with their higher rescue rates in the global analysis, whereas DDR1-containing pairs cluster closer to parity or show smaller gains (Table 3). Representative examples include TSLP–ERBB2, where edits such as Y30S yielded multiple successful variants, and DDR1-containing combinations that required more extensive exploration with lower overall success. Pairs with Z13-IL22-2 show no measurable rescue, consistent with the refractory behavior noted previously (Table 2, Table 3).

**Table 3.**
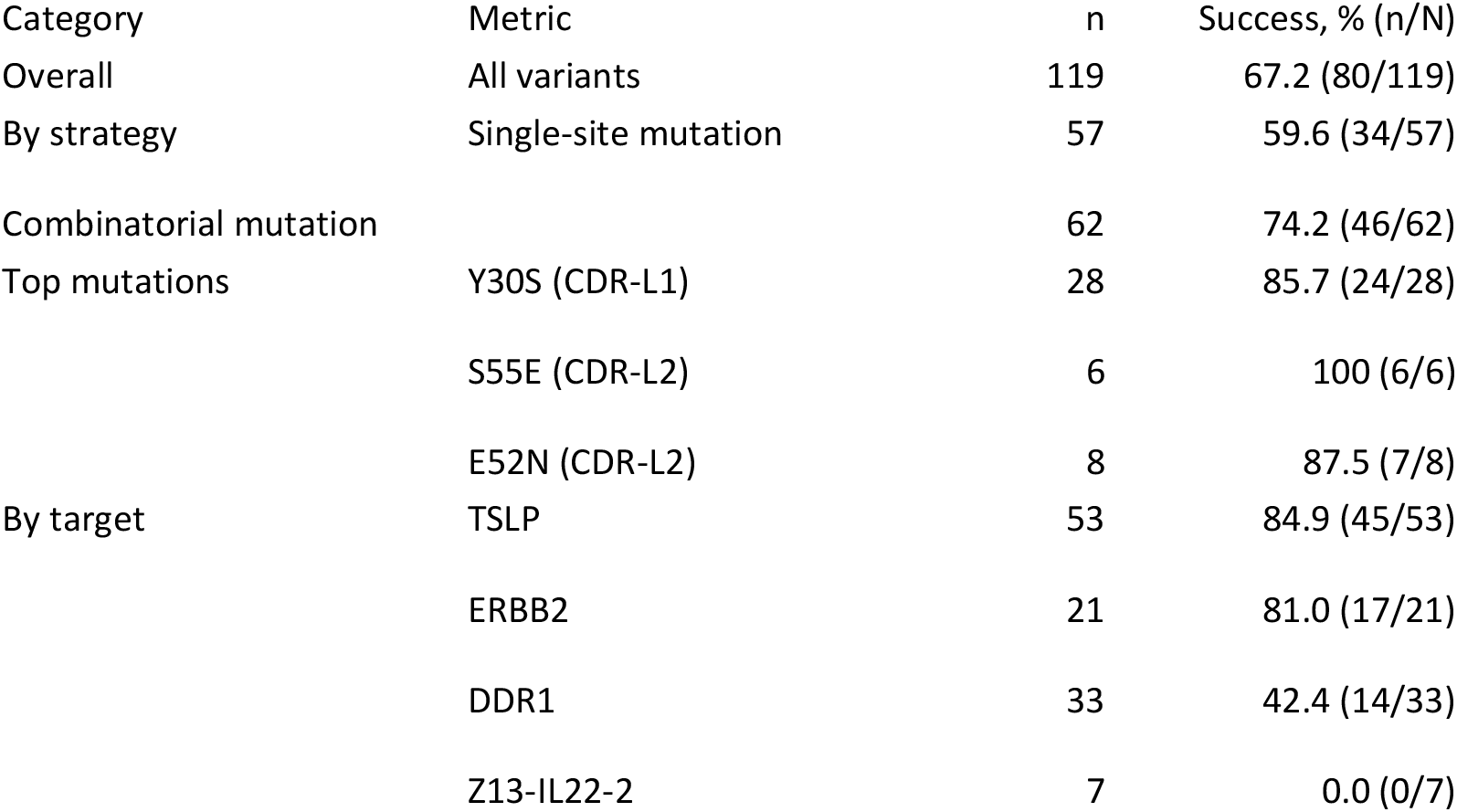
Structure-guided optimization. Success defined as restoration or maintenance of binding while preserving compatibility with both HC partners in the bispecific format. Top mutations shown represent individual substitutions tested across multiple constructs. Positions use Chothia numbering; success rates reflect context-dependent recurrence across frameworks. Complete per-variant data including binding kinetics, developability metrics (yield and purity), and positional/chemical substitution analyses in Supplementary Tables S4–S5.

| Category | Metric | n | Success, % (n/N) |
| --- | --- | --- | --- |
| Overall | All variants | 119 | 67.2 (80/119) |
| By strategy | Single-site mutation | 57 | 59.6 (34/57) |
| Combinatorial mutation |  | 62 | 74.2 (46/62) |
| Top mutations | Y30S (CDR-L1) | 28 | 85.7 (24/28) |
|  | S55E (CDR-L2) | 6 | 100 (6/6) |
|  | E52N (CDR-L2) | 8 | 87.5 (7/8) |
| By target | TSLP | 53 | 84.9 (45/53) |
|  | ERBB2 | 21 | 81.0 (17/21) |
|  | DDR1 | 33 | 42.4 (14/33) |
|  | Z13-IL22-2 | 7 | 0.0 (0/7) |

Third, designs that ultimately succeeded most often included substitutions in the light-chain CDR-L1 and/or CDR-L2 loops, with the specific sites being VL- and antigen-dependent. Pairs exhibiting the largest improvements with substitutions in CDR-L1/L2, concordant with the region-level trends (CDR-L2 > CDR-L1 > CDR-L3) from the prior subsection. Fig. 4B provides structural context with representative sites (e.g., Y31F in L1 and E49L/E52N/S55E/S50D in L2) positioned at the VH–VL/antigen interface. These exemplars illustrate how coordinated, multi-residue designs at L1/L2 can relieve a dominant constraint on one arm while preserving compatibility on the other.

Together, the pair-wise view in Fig. 4A reinforces the central theme of multi-interface optimization: most practical wins arise from targeted, combinatorial edits that rescue the limiting arm without degrading the partner. This interpretation is consistent with the aggregate success rates, the L1/L2 hot-spot pattern, and the target-dependent permissiveness quantified in Table 3. Having established that arm-specific rescue recovers BsAb binding across 7/10 targets, we evaluated whether cLC designs translate into lot-level benefits in expression, purity, and product homogeneity.

### Manufacturability gains from cLC design

cLC designs reduced mispairing and improved downstream metrics, yielding higher titers and SEC purity in the BsAb context relative to native LC baselines (see Table 1 for success definitions). Three BsAbs—VEGFA×MTSP1(002), ERBB2×HRAS (006), and VEGFA×ERBB2 (051)—were produced with KiH mutations^6^ combined with our cLC design, and corresponding parental LC as baseline (Fig 5A, Supplementary Table S3). The KiH technology effectively prevents HC-HC homodimerization (>92% heterodimerization efficiency)^6^, allowing us to focus exclusively on resolving LC mispairing. This experimental design isolates the impact of our cLC platform by controlling heavy chain assembly. Expression yields reached 1.6–1.8 g/L and are consistent with the upper end of reported ranges. For context, reported bispecific titers vary widely by format and stage; early stable pools and some formats often report hundreds of mg/L^16,62^ and comparable to advanced cLC platforms. In 2/3 designs, purity improved from ∼34–68% to ∼94–95% target species; the third is limited by arm expression balance (tunable by HC1:HC2 transfection ratio). LC-MS confirmed cLC-006 and cLC-051 designs were correctly assembled, with target percentage improved from baseline 33.9%-67.8% to cLC 93.7%-95.2% (Fig 5C). cLC-002 showed a low target percentage of 34.0% and 63.1% Half Ab (Supplementary Table S12, Supplementary Fig. 8), which was mostly due to the unbalanced expression level between two arms. Excess amount of arm1 forms HC1*2+LC1*2 (2.9%, HC-mispairing cLC), and HC1+LC1(58.3%, including free Arm1 and non-covalent binding HC1*2+LC1*2). Such an imbalance could be fixed by adjusting the transfection ratio of HC1:HC2, which is beyond discussion of this study.

**Figure 5.**
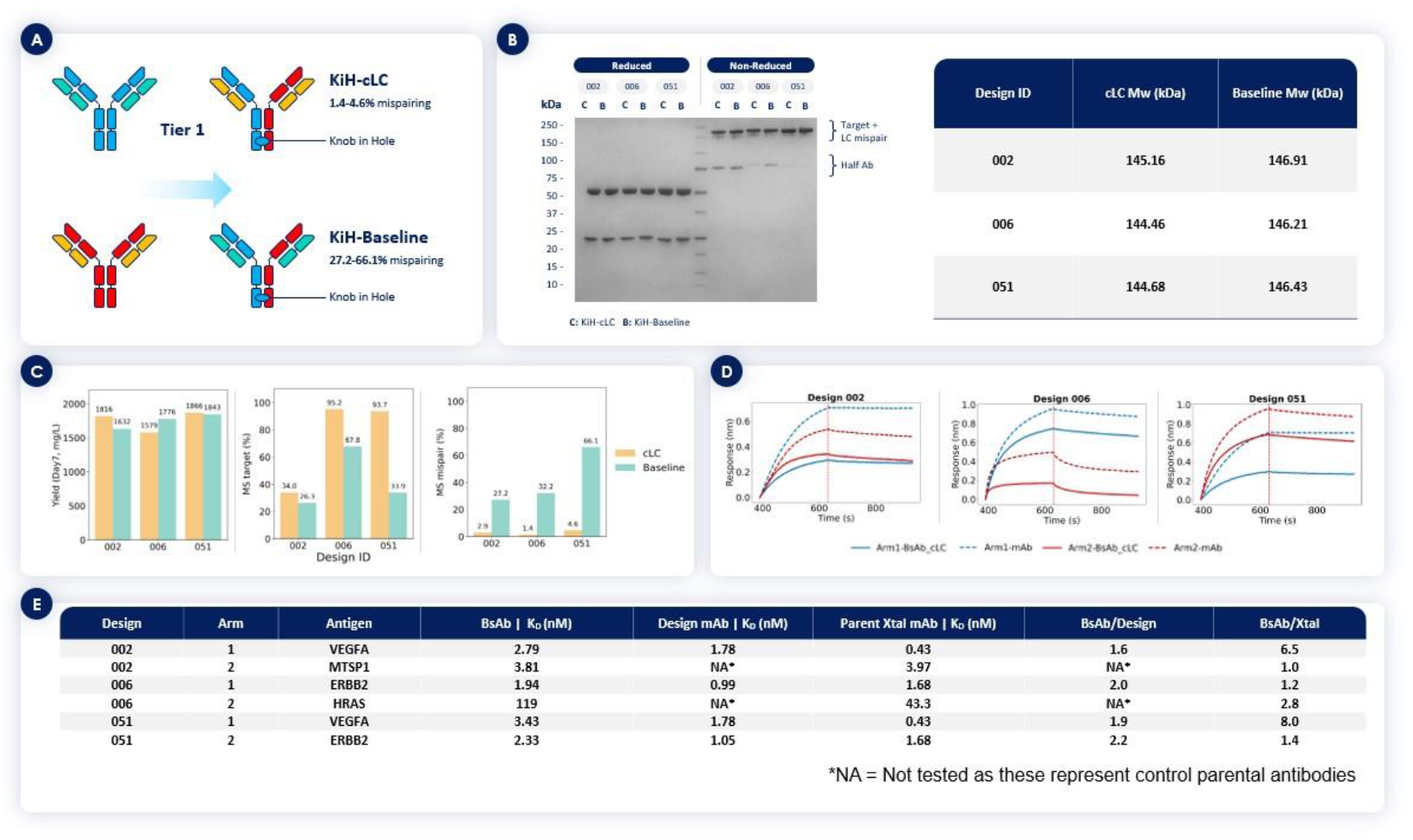
Manufacturing improvements and binding validation of BsAbs with cLCs. (A) Schematic showing Tier 1 comparison between KiH-cLC and KiH-Baseline (parental LCs) formats with expected mispairing reduction. (B) Reduced and Non-reduced SDS-PAGE analysis of three BsAbs designs showing reduced half-antibodies (∼75 kDa) in cLC constructs versus baseline. Molecular weight markers indicated. (C) Expression yields (Day 7, mg/L), percentage of target species and mispaired species by mass spectrometry for baseline versus cLC constructs across three designs. (D) Representative biolayer interferometry sensorgrams showing binding kinetics for Design IDs:002, 006, and 051 at indicated concentrations. (E) Summary table of binding affinities (K_D_) and fold-changes for all three bispecific designs comparing baseline and cLC formats against both target antigens.

### Binding affinity validation

To confirm therapeutic viability in the cLC format, we characterized binding kinetics for the three BsAbs designs against their respective targets (Fig. 5D, E). Biolayer interferometry of 002 (VEGFA×MTSP1) showed VEGFA binding at 2.79 nM, representing a 6.5-fold change from baseline. The K_D_ for MTSP1 binding was 3.81 nM, representing no change from baseline (1.0-fold). 006 (ERBB2×HRAS) maintained ERBB2 affinity at 1.94 nM, a 1.2-fold change from baseline, while HRAS showed 119 nM, representing a 2.8-fold change. 051 (VEGFA×ERBB2) demonstrated VEGFA binding at 3.43 nM with 8.0-fold change and ERBB2 at 2.33 nM with 1.4-fold change. These affinity changes are consistent with expected 2- to 10-fold variations when transitioning from bivalent parental IgG to monovalent bispecific binding, reflecting both loss of avidity and altered VH-VL interface geometry^1,4,75^.

## Discussion

### Computational design resolves LC pairing constraints in bispecific antibodies

We present a computational platform that transforms bispecific antibody development by addressing the chain-pairing bottleneck through predictive, structure-led pre-selection and targeted rescue. At the target level, engineered cLCs succeeded for 7/10 targets. At the construct level, structure-guided optimization achieved 67.2% success (80/119 variants) compared to 38.8% baseline (33/85), with screening depths of ∼1 to >30 constructs per target. For bispecific assembly, we generated 24 functional BsAbs from 55 combinations (43.6% success rate)—concentrating validation on compact, model-prioritized sets rather than broad empirical libraries. Exemplars met manufacturability benchmarks (SEC purity (target species) >90%, titers 1.6–1.8 g/L).

Our optimization strategy addresses a focused problem: given pre-existing VH–antigen binders, we computationally identify cLCs pairing productively with one or two non-cognate VHs. Although we assume pre-existing VHs here, the same pre-selection logic is compatible with upstream VH discovery, suggesting a path toward end-to-end computational BsAb design as prediction tools mature. Tier 2 validation demonstrates the most stringent case—selecting VLs with no prior association to either VH—using limited experimental candidates per target (Fig. 1 inset).

### Key Computational Innovations Enable Predictive Design

The critical innovation lies in embedding VH–antigen restraints into AI-based structure prediction to enforce correct LC positioning; without restraints, models frequently misplace the LC (Supplementary Fig. S10). Our restraint-guided approach with MD validation proved essential for multi-chain complexes. Notably, the platform succeeded with therapeutic VLs lacking experimental structures and enabled mouse-to-human germline transitions (17 cases, Supplementary Fig. S9), extending applicability beyond PDB-represented space.

Where empirical strategies often require large variant sets, our computational pre-selection concentrates validation to <∼20 candidates per target. We treat this reduction as practical rather than absolute, as it depends on target complexity and prior knowledge. Three BsAbs reached production specifications (>90% SEC purity, 1.6–1.8 g/L titers), supporting design accuracy and therapeutic viability. Our rescue strategy salvages initially failed designs via structure-guided edits rather than discarding candidates.

### Structural Determinants of Promiscuous Light Chain Pairing

Successful designs showed enrichment of IGKV1-39 (11% in templates → ∼30% in validated cLCs), whose CDR-L3 loop tends to include smaller residues that accommodate diverse CDR-H3 conformations and may reduce steric clashes (Fig. 3). J-gene usage remained relatively stable (IGKJ1: ∼40–60%), suggesting the V-gene as the primary compatibility determinant. However, given biases in PDB/Thera-SAbDab toward crystallizable therapeutics, we treat IGKV1-39 as a practical heuristic rather than definitive evidence; prospective tests on unbiased repertoires are needed. Validated cLCs also spanned multiple κ and λ families, indicating generalizability beyond IGKV1-39.

### Multi-Interface Optimization Reveals Fundamental Design Principles

Structure-guided mutagenesis of cLCs recovered binding across multiple targets, with clear target-dependence: systems with VH-dominated binding and minimal VL–antigen contacts tended to be permissive, whereas targets with extensive VL engagement were refractory (e.g., TSLP/ERBB2 versus DDR1/Z13-IL22-2; Fig. 4A and Supplementary Fig. S4). These patterns align with broader observations on BsAb interface constraints and format-dependent liabilities^1^. Unlike single-interface optimization, cLC engineering is inherently multi-objective: edits must satisfy two VH–VL interfaces while maintaining two antigen-binding modes^4^. In practice, asymmetric trade-offs can be productive— accepting modest affinity losses in one arm to secure larger gains in the other— consistent with coupled but non-identical constraints across the arms^1,17^. Position-level trends highlight CDR-L1/L2 as frequent levers alongside CDR-L3. Recurrent productive substitutions (e.g., Y30S, Y31F, S50D, E52N, S55E) enabled compatibility across multiple VH backbones, suggesting that L1/L2 repositioning and electrostatic re-balancing are common solutions rather than exceptions (Supplementary Table S4). Consistent with prior experience in structure-guided antibody design, multi-site edits typically outperformed single-site changes, supporting a view of cooperative interface tuning^61^. Beyond affinity, several substitutions addressed developability liabilities (e.g., removing unpaired cysteines via Cys→Ser/Ala), allowing binding rescue and manufacturability to be optimized in the same loop^76^. As precedents, the FIXa/FX bsAb programs Hemlibra and Mim8 exemplify multi-dimensional optimization coupled to clinical translation; our approach systematizes that design playbook computationally ^13,77^

### Limitations and Future Directions

Per-site mutational exploration is uneven, which can bias apparent success rates. Variable outcomes across targets indicate that current scoring may not fully capture expression/stability effects, reflecting both the difficulty of modifying optimized therapeutics and limitations in evaluating non-cognate VH–VL pairs. While affinity is a validated surrogate for many modalities, functional assays (e.g., neutralization, ADCC) remain necessary to establish translational potential.

Future work should reduce dependence on PDB templates via AI-predicted structures, leverage repertoire-scale priors ^63^, incorporate immunogenicity prediction ^60^, and include in vivo validation. Coupling cLC pre-selection with upstream VH discovery could enable more fully computational design flows; extension to trispecifics and ADCs would broaden utility. Broader perspectives on AI-driven approaches to multispecific engineering are discussed elsewhere^57^.

## Conclusion

We demonstrate that computational design addresses the chain-pairing bottleneck in bispecific antibodies, shifting effort from large empirical libraries to compact, model-prioritized sets while achieving 43.6% success for generated BsAbs across tested constructs. Three BsAbs met manufacturability benchmarks (SEC purity (Target species) >90%, 1.6–1.8 g/L), supporting therapeutic viability. By lowering the number of constructs advanced to the bench, this approach reduces experimental burden and can broaden feasibility for teams with more limited resources, while maintaining a cautious, target-dependent outlook.

