## Supplemental Material for "AI-guided design of common light chains to enable manufacturable bispecific antibodies"

### Supplementary Information

#### Supplementary Notes 1: Current Computational Approaches to Common Light Chain Design

Computational approaches to cLC design have emerged but remain limited in scope and efficiency. Sequence-based libraries demand extensive screening[1], structure-based interface engineering still requires dozens of variants[14,15], and generative AI narrows the search to $10^3$–$10^5$ candidates but with substantial validation[16]. Even recent rational selection approaches achieve only 1.4% success rates (2/144 variants)[67], highlighting the continued challenge of predictive cLC design. While traditional phage display screens $10^9$–$10^{11}$ variants[64], recent approaches reduce this to thousands of candidates. However, even state-of-the-art pipelines generate $10^5$–$10^6$ designs, filter hundreds, and still require 10–20 experimental validations[17]. To date, no method has achieved de novo functional cLC design with minimal empirical testing.

#### Supplementary Methods

##### High-Resolution Antibody-Protein Antigen Complex Dataset

The February 2025 snapshot of the Structural Antibody Database (SAbDab)[20] was downloaded from https://opig.stats.ox.ac.uk/webapps/sabdab-sabpred/sabdab/search/?all=true. Structures where the antigen type contained the "protein" keyword and excluded those containing "peptide", "carbohydrate", or "nucleic-acid" descriptors were retained to focus specifically on therapeutically relevant antibody-protein interactions. For structures containing multiple conformations, only the first model in each file was retained. This curation resulted in 8,183 antibody-protein antigen complexes from 4,020 unique PDB identifiers.

##### Structural and Metadata Repository

The first biological assembly for all 4,020 PDB entries was downloaded from the Research Collaboratory for Structural Bioinformatics (RCSB) Protein Data Bank[27,28]. When PDB files were unavailable, mmCIF files were downloaded instead. Forty-nine obsolete entries were identified and removed from the dataset. Structure resolution, complete FASTA sequences, UniProt identifiers, protein-coding genes, and source organisms were retrieved using the RCSB GraphQL data API endpoint.

Structures with resolution coarser than 3.0 Å were excluded to ensure high-quality structural data for computational modeling[29]. All antibody chains were numbered according to the Chothia numbering scheme[30] using ANARCI[31], and variable fragments (Fv) with corresponding antigen chains in contact were extracted. Structures containing missing coordinates within the Fv region were removed, resulting in 3,639 high-quality structures selected as modeling candidates.

#### Single-Chain Template Preparation

Antibody heavy chain variable domains (VH) and antigen chains served as structural templates for prediction models. All protein chains were standardized using PDBFixer[72] and Gemmi[33] to remove alternate atom locations and replace non-canonical amino acids with standard residues.

For antigen chains with missing coordinates, Protein Generator (PG)[34], a diffusion model for co-designing protein sequences and structures, was employed to predict missing regions. PG was conditioned on resolved structural segments and full-length sequences to guide model reconstruction toward reference conformations with known sequences. For each antigen, 50 diffusion time steps were sampled with a noise factor of 1.0. Ten designs were generated per antigen, and the structure with the highest predicted local-distance difference test (pLDDT) score[18] and lowest backbone root-mean-square deviation (RMSD) relative to the resolved structure was selected as the template. The selected structure models for each antigen all had > 0.9 pLDDT and < 0.2 Å backbone RMSD to the corresponding resolved structures.

All VH and antigen templates were energy-minimized using Rosetta[35] with heavy atom coordinate constraints to optimize rotamer conformations and reduce risk of structure prediction model overfitting to input sequences[40].

#### Modeling of Novel VH-VL Combinations

For each reference VH-variable light (VL) pair, alternative VL sequences were collected where all three complementarity-determining regions (CDRs) exhibited length variations within ±2 amino acids of the original VL CDRs. This constraint maintains structural similarity while allowing sufficient sequence diversity, based on natural antibody CDR length distributions observed in therapeutic antibody databases[41]. The alternative VL sequences were collected from the high-quality PDB dataset and the Thera-SAbDab[71] database. Each VL sequence, together with corresponding VH and antigen sequences, was queried against UniRef30[38] using MMSeqs2[39] to generate multiple sequence alignments (MSAs) for evolutionary context.

Structure prediction was performed using Chai-1[23], an open-source implementation of AlphaFold 3[40]. The template module of Chai-1 was modified to accept custom local structure files as templates. Chai-1 parameters were configured with 5 trunk recycling iterations, 200 diffusion steps, and 5 structure models per prediction run[45]. The highest-scoring model based on Chai-1's aggregate confidence score was selected for subsequent analysis.

#### Enhanced Modeling with Structural Restraints

To improve prediction accuracy for antibody-antigen complexes, we implemented a restraint-guided modeling approach. All VH-antigen residue pairs forming cation-π, π-stacking, salt bridge, or hydrogen bond interactions were identified from reference structures using geometric criteria[42,46]. Sidechain centroid distances for these interactions were provided to Chai-1 via restraint files using the "max_distance_angstrom" parameter. These inter-chain distance restraints, combined with single-chain structure templates, guided Chai-1 in accurately folding antibody-antigen complexes while preserving critical binding interactions.

The optimal Chai-1 structure model was superimposed onto the corresponding reference structure using VH chain alignment. A hybrid structure was constructed by combining the reference VH, predicted VL from Chai-1, and reference antigen chains.

#### In Silico Candidate Selection Pipeline

The computational screening pipeline consists of sequential filtering steps designed to identify high-quality cLC candidates (see Figure 1 and Supplementary Fig. 1 for a complete workflow), with the complete experimental workflow illustrated in Supplementary Figure 7. Starting from reference antibody-antigen structures, the pipeline generates alternative VL sequences, predicts new complex structures, and applies multiple validation criteria to ensure both structural integrity and functional preservation. This hybrid model was subsequently energy-minimized using Rosetta with heavy atom constraints to generate the final structure model.

#### Structure Model Quality Assessment

Candidate structures underwent multi-tier quality filtering. Initial screening used Chai-1 confidence metrics, retaining models with overall predicted template modeling scores (pTM) > 0.7 and VH-VL interface predicted template modeling scores (ipTM) > 0.7[43]. The pTM score reflects global structure confidence, while ipTM specifically evaluates interface prediction quality[18]. To preserve critical epitope interactions, candidates with VH backbone root-mean-square deviation (RMSD) or heavy chain CDR3 (HCDR3) backbone RMSD > 2.0 Å relative to template structures were excluded.

Secondary filtering employed Rosetta energy functions and statistical metrics. Percentile thresholds (10th and 90th percentiles) were calculated from SAbDab human antibody distributions using identical scoring procedures. Candidates were required to satisfy shape complementarity (sc_value) > 0.5428, packing statistics (packstat) > 0.6536, unsatisfied hydrogen bonds < 18, and per-residue interaction energy < -1.0. Rosetta InterfaceAnalyzer[44] was applied to VH-VL interfaces, requiring sc_value > 0.6 and packstat > 0.65 to eliminate candidates with unfavorable interface geometries.

#### Molecular Dynamics Simulation Protocol

Topology and coordinate files were prepared using tLeap from AmberTools[45], employing the AMBER FF14SB force field[46]. Protein complexes were immersed in dodecahedral TIP3P water boxes[73], maintaining 9 Å distance between solute and box edges. Periodic boundary conditions were applied with long-range electrostatics calculated using particle-mesh Ewald (PME) method[47].

The equilibration protocol consisted of steepest descent minimization (forces < 1000 kJ/mol·nm), followed by conjugate gradient minimization (forces < 500 kJ/mol·nm). Systems underwent heating to 310 K during 1 ns NVT simulations with 2 fs time steps using the Bussi thermostat[48], with bond lengths constrained using LINCS algorithm[49]. Subsequently, 1 ns NPT simulations were conducted with pressure maintained at 1 bar using Nosé-Hoover thermostat[50] and Parrinello-Rahman barostat[51]. All simulations were performed using GROMACS[52].

Production trajectories consisted of ten independent 6 ns simulations per complex, each initialized with random velocities. Simulations used 1.4 nm cutoffs for van der Waals and real-space PME interactions. The first 1 ns of each simulation was discarded as equilibration, with analysis performed on the final 5 ns (200 frames) to ensure equilibrated sampling.

#### Protein-Protein Interaction Preservation Analysis

Protein-protein interactions (PPIs) were analyzed using Arpeggia[24], a geometry-based tool for quantifying interatomic interactions in protein structures. The tool identifies cation-π, π-stacking, salt bridge, and hydrogen bond interactions across simulation trajectories. For VL-antigen interfaces, electrostatic repulsions and van der Waals contacts were additionally monitored.

A PPI was classified as stable when: (a) more than half of trajectory replicates exhibited the interaction in more than half of their frames, and (b) the interaction was observed in ≥70% of frames on average across all replicates. The 70% threshold was established through systematic analysis of interaction stability in validated antibody-antigen complexes and represents a balance between stringency and practical filtering efficiency.

**VH-Antigen Interface Preservation**: Candidates were retained if ≤2 original VH-antigen interactions became unstable, or if ≥70% of original interactions remained stable, ensuring preservation of critical binding determinants.

**VL-Antigen Interface Assessment**: For VL-antigen interactions observed in reference structures, corresponding antigen positions were monitored in candidate trajectories. Candidates passed this filter with ≤2 unstable PPIs or ≥70% stable interactions. Candidates exhibiting stable electrostatic repulsions or van der Waals contacts absent in reference structures were eliminated to prevent non-native binding modes.

**VH-VL Interface Evaluation**: VH-VL interactions were assessed at VH positions where reference structures exhibited VL contacts. Candidates were required to maintain ≤4 unstable PPIs or ≥70% stable interactions, with the higher tolerance reflecting the engineered nature of these interfaces.

#### AbAngle Conformational Analysis

The six AbAngle parameters[53], which quantify relative VH-VL domain orientations through angular and distance measurements, were calculated for all simulation frames. Values were summarized using 25th and 75th percentile distributions (Q1 and Q3). Reference trajectory statistics included minimum, maximum, and standard deviation for each parameter. Candidates passed AbAngle filtering when Q1 > min(reference) - std(reference) and Q3 < max(reference) + std(reference), ensuring conformational compatibility with native antibody geometries.

#### MM/GBSA Binding Energy Calculations

Molecular mechanics/generalized Born surface area (MM/GBSA) calculations[54] were performed using GROMACS to estimate binding free energies. Calculations were conducted separately for VH-antigen and VH-VL interactions across all trajectory frames using the single-trajectory approach[55]. Delta binding energies were aggregated to 25th and 75th percentile values, with reference trajectory statistics providing filtering thresholds. Candidates satisfied MM/GBSA criteria when Q1 > min(reference)

- std(reference) and Q3 < max(reference) + std(reference), indicating energetically favorable binding relative to native complexes.

#### Mutation of VL Candidates

Structural incompatibilities were systematically evaluated using the Rosetta protein design suite. The parental antibody was superimposed to the candidate using the shared VH chain, and potential VL mutations were selected based on proximal residues and rational design of the interface. For each set of mutations, the selected residues were mutated to new residues using Rosetta FastDesign[35] with the energy function ref2015[36], and residues within 8 Å of the mutated residues were allowed to repack. In parallel, the same selection of residues was repacked, and the delta Rosetta score between the mutated and non-mutated systems were collected in 30 independent runs. Mutations with an average delta score <2.0 were selected for experimental characterization.

#### DNA Construction

Variable heavy (VH) and light (VL) chain genes were synthesized by Genewiz (Azenta Life Sciences) and cloned into pcDNA3.4 vectors (Thermo Fisher Scientific) using Gibson assembly[56]. To minimize protein heterogeneity, all heavy chains were designed with human IgG1 scaffolds (UniProt: P01857), and all light chains included human kappa constant sequences (UniProt: P01834). Following sequence verification by Sanger sequencing, transfection-grade DNA was prepared using QIAprep Spin Miniprep Kits (Qiagen, 27106) and mixed at 1:1 ratio of cognate/non-cognate heavy:light chains for IgG expression.

Selected designs were constructed into common light chain bispecific antibodies using human IgG1 scaffolds with heavy chain knobs-into-holes mutations[6] (T249W, T249S, L251A, Y290V). To facilitate bispecific antibody purification, FLAG tags (GGDYKDDDDK) or His tags (GHHHHHH) were fused to the C-terminus of each heavy chain.

#### Cell Culture and Protein Expression

Proteins were transiently expressed using ExpiCHO-S cells (ThermoFisher, A29127) following optimized protocols. Cells were cultivated in custom-developed medium supplemented with 1% HT supplement (Gibco, 11067030) and 4 mM GlutaMAX (Gibco, 35050061). Before transfection, cells were concentrated to $8 \times 10^6$ viable cells per mL. An automated, high-throughput approach was employed using the Hamilton STAR liquid handling system in 96-well plate format. For transfection, 0.9 μg DNA was complexed with 1.35 μg FectoPRO reagent (Polyplus, 101000014) in 100 μL Opti-MEM (Gibco, 31985062) for 10–15 minutes, then added to 900 μL resuspended cells. Nutrient-rich feed was supplemented on days 1 and 4 post-transfection, with conditioned medium harvested on day 7.

#### Purification

Protein purification was conducted using the Hamilton STAR system with columns packed with MabSelect PrismA resin (Cytiva, 17549801). Clarified supernatant was loaded through 10–20 cycles of repeated aspirations and dispensing. Columns were washed with phosphate-buffered saline and proteins eluted using pH 3.0 acetic acid, immediately neutralized with Tris buffer to final concentrations of 0.1 M acetic acid, 130 mM Tris, pH 5.5. This yielded 230 μL of each IgG variant. Protein concentrations were quantified by UV absorbance at 280 nm using protein-specific extinction coefficients.

His-FLAG-tagged bispecific antibodies underwent tandem purification using Ni-NTA resin (Cytiva, 17526802), anti-FLAG resin (Merck, A2220), and Superdex 200 Increase chromatography (Cytiva, 28990944). Final fractions were collected in PBS buffer for downstream assays.

#### Protein Quality Analysis

Protein purity was analyzed using micro capillary electrophoresis (MCE) and analytical size-exclusion chromatography (SEC). For MCE, 6 μL protein samples were mixed with 21 μL sample buffer (8.4 mM Tris-HCl pH 7.0, 7.98% glycerol, 2.38 mM EDTA, 2.8% SDS) under non-reducing (2.4 mM iodoacetamide) and reducing (0.1 M DTT) conditions. Samples were heated at 85°C for 10 minutes and analyzed on a Caliper LabChip GXII Touch instrument (PerkinElmer). Analytical SEC was performed on an Agilent 1260 Infinity II LC System using TSKgel G3000SW XL columns (Tosoh, 08541) with

50 mM sodium phosphate, 300 mM NaCl, pH 6.8 running buffer at 1 mL/min flow rate.

#### Protein Mass Spectrometry

Approximately 200 pmol of each purified sample was digested with PNGase F or IdeS enzyme to remove N-glycans or obtain smaller subunits for enhanced MS sensitivity. Samples were analyzed both non-reduced and DTT-reduced using an Agilent 1290 Infinity II UPLC connected to an Agilent 6230 ESI-TOF mass spectrometer. Raw data was deconvoluted using Agilent MassHunter BioConfirm software (Version 10.0), with deconvoluted masses matched to theoretical molecular weights requiring absolute mass errors <100 ppm.

#### Crystallography Methods for Fab Fragment B3BC-T13-R2-M027

For crystallography, Fab fragment B3BC-T13-R2-M027 was expressed in HEK293T cells for secretion using the pOPIN system[65], with the light chain modified to improve crystallization[66]. After four days, media was collected and filtered for purification via HisTrap HP column (Cytiva). After concentration and buffer exchange to 0.5 mg/ml in 20 mM Tris pH 7.5 and 200 mM NaCl, the purified protein was screened for crystallization conditions using the TOP96 kit (Molecular Dimensions). Three-dimensional crystals grown in 0.1 M MES pH 6.5 and 12% PEG-20000 were harvested diffracting to 2.91Å at the Advanced Photon Source, beamline 24-ID-E, managed by the North Eastern Collaborative Access Team. The data was collected at 100 K on an Eiger detector. Using the 75% sequence identical structure 3QEG as a molecular replacement model, a model for the structure of the Fab fragment was iteratively refined using PHENIX[67-69] and manually corrected using Coot[70].

**Supplementary Table S12. Data collection and refinement statistics**

| Parameter | Fab27 |
|---|---|
| **Data Collection** | |
| Wavelength | 0.9792 |
| Resolution range | 62.36 - 2.91 (3.014 - 2.91) |
| Space group | P 21 21 21 |
| Unit cell | 52.704 70.405 134.321 90 90 90 |
| Total reflections | 48665 (8160) |

| | |
|---|---|
| Unique reflections | 11245 (1129) |
| Multiplicity | 4.3 (4.5) |
| Completeness (%) | 97.64 (99.47) |
| Mean I/sigma(I) | 5.4 (3.0) |
| Wilson B-factor | 42.71 |
| R-merge | 0.226 (0.453) |
| R-meas | 0.291 (0.584) |
| R-pim | 0.065 (0.364) |
| CC1/2 | 0.980 (0.364) |
| **Refinement** | |
| Reflections used in refinement | 11245 (1129) |
| Reflections used for R-free | 554 (46) |
| R-work | 0.2833 (0.3339) |
| R-free | 0.3343 (0.3514) |
| **Model** | |
| Number of non-hydrogen atoms | 3038 |
| - macromolecules | 3022 |
| - ligands | 38 |
| - solvent | 0 |
| Protein residues | 410 |
| **Geometry** | |
| RMS(bonds) | 0.003 |
| RMS(angles) | 0.64 |
| Ramachandran favored (%) | 92.39 |
| Ramachandran allowed (%) | 7.36 |
| Ramachandran outliers (%) | 0.25 |
| Rotamer outliers (%) | 3.06 |
| Clashscore | 8.50 |
| **B-factors** | |
| Average B-factor | 42.34 |
| - macromolecules | 42.40 |
| - ligands | 31.84 |
| Number of TLS groups | 6 |

*Statistics for the highest-resolution shell are shown in parentheses.*

#### Binding Affinity Characterization

Binding affinities ($K_D$ values) were assessed using biolayer interferometry on a ForteBio Octet RED384 instrument. Antibodies were immobilized on anti-human Fc capture sensors (Sartorius, 18-5142), followed by serial dilution of respective antigens. Assays were designed to measure monovalent 1:1 binding interaction at room temperature with 1000 RPM stirring. Data was processed using Octet BLI Systems Software with 1:1 binding model globally applied to obtain association ($k_{on}$) and dissociation ($k_{off}$) rate constants. Equilibrium dissociation constants ($K_D$) were calculated as $k_{off}/k_{on}$ ratios.

#### Thermal Stability Assessment

Thermal stability was assessed using intrinsic fluorescence on an Uncle instrument (Unchained Labs). Nine microliters of protein samples were dispensed into UNi cuvettes (Unchained Labs, 201–1010) and subjected to controlled thermal ramping from 20°C to 95°C at 0.5°C/min. Melting temperatures ($T_M$) were determined by analyzing first derivatives of barycentric mean fluorescence intensity. Aggregation temperatures (Tagg) were calculated from scattered light intensity at 266 nm or 473 nm wavelengths.

#### Statistical Analysis and Reproducibility

All computational predictions were performed in triplicate with different random seeds to ensure reproducibility. Statistical significance was assessed using two-tailed t-tests with Bonferroni correction for multiple comparisons where appropriate. Confidence intervals (95%) were calculated for all binding energy measurements. The computational pipeline success rate was defined as the percentage of candidates passing all filtering criteria that demonstrated experimental validation.

Random seeds were recorded for all stochastic processes: Chai-1 predictions (seed 42), Rosetta sampling (seeds 1111111–1111141), and MD simulation initial velocities (generated using GROMACS default random number generator with documented seeds). All structure predictions were performed on NVIDIA A100 GPUs with 40 GB memory. MD simulations utilized CPU clusters with Intel Xeon processors.

Sequences of all 204 constructs are available in Supplementary Table S10.

### Supplementary Figures

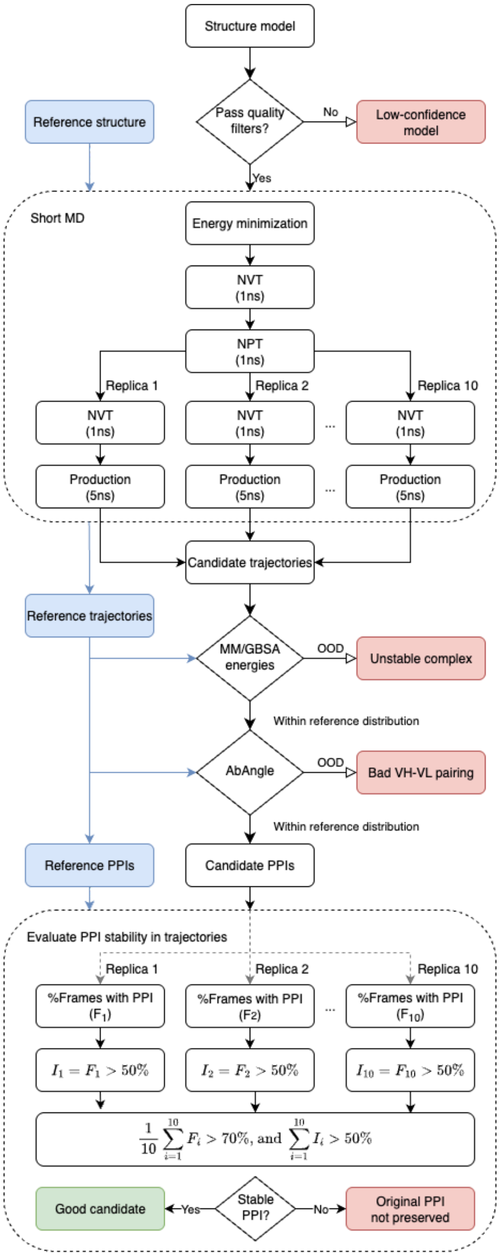

Supplementary Figure S1 **Detailed computational pipeline for common light chain engineering with filtering parameters and decision criteria.**

**(A) Complete workflow schematic** showing all computational steps from initial dataset curation through experimental validation. Starting dataset: 8,183 antibody-antigen complexes from SAbDab, filtered to 3,639 high-resolution structures (≤3.0Å). Structure modeling employs Chai-1 with distance restraints derived from reference VH-antigen interactions (cation-π, π-stacking, salt bridges, hydrogen bonds). Numbers at each filtering step indicate candidates remaining, with branching pathways showing alternative routes for candidates failing specific criteria.

**(B) Quality filtering cascade** with specific parameters: Chai-1 confidence metrics (pTM>0.7 for overall confidence, ipTM>0.7 for interface confidence), structural preservation (VH backbone RMSD<2.0Å, HCDR3 backbone RMSD<2.0Å), Rosetta energy validation (shape complementarity >0.5428, packing statistics >0.6536, unsatisfied hydrogen bonds <18, per-residue interaction energy <-1.0), and interface analysis (VH-VL shape complementarity >0.6, packing statistics >0.65). Statistical thresholds were derived from 10th and 90th percentiles of SAbDab human antibody distributions (n=3,639).

**(C) Molecular dynamics validation protocol** showing 10 independent 6 ns simulations per candidate with detailed equilibration procedures. Protein-protein interaction (PPI) preservation analysis using geometric criteria: stable interactions defined as present in >50% of trajectory replicates with >70% frame occupancy. Interface-specific tolerance thresholds: VH-antigen (≤2 unstable interactions or ≥70% stable), VL-antigen (≤2 unstable interactions or ≥70% stable), VH-VL (≤4 unstable interactions or ≥70% stable). The 70% stability threshold was empirically determined through systematic analysis of interaction persistence in validated antibody-antigen complexes.

**(D) Final validation filters** including AbAngle conformational analysis (6 geometric parameters: HC1, HC2, LC1, LC2, DC, and opening angle, all within reference mean ± 1 standard deviation range) and MM/GBSA binding energy calculations for both VH-antigen and VH-VL interfaces. Energy distributions were compared to reference trajectories using 25th and 75th percentile bounds to ensure energetically favorable binding.

**(E) Branch points and recovery pathways** illustrating decision trees for handling edge cases, alternative parameter sets for borderline candidates, and mutation strategies for failed designs. Quality control checkpoints include manual inspection of top candidates and cross-validation with experimental results. Success rates: initial structure prediction (85%), quality filtering (45%), MD validation (25%), final experimental validation (>90% of computationally validated candidates).

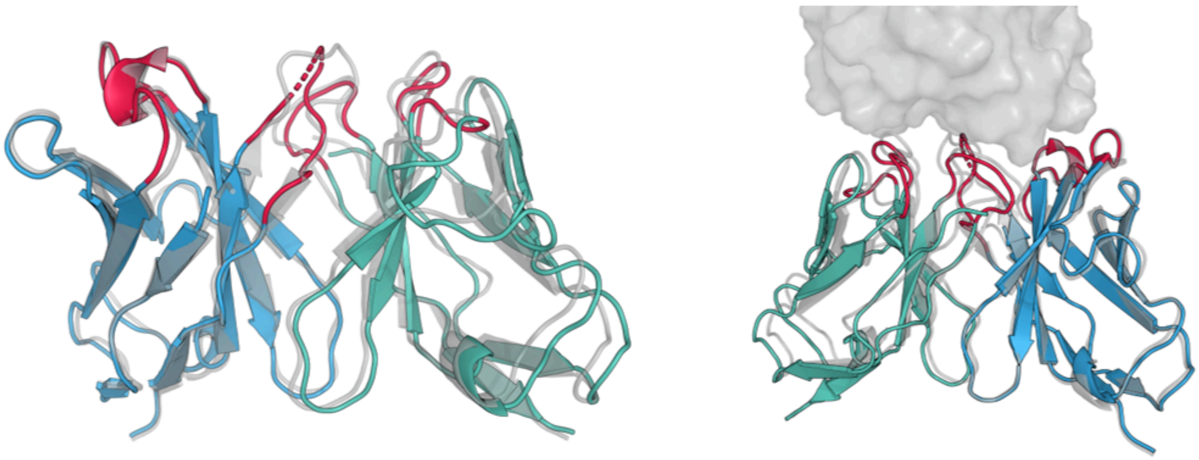

**Supplementary Figure S2. Impact of distance restraints on Chai-1 structure prediction.** Comparison of Chai-1 predictions with and without VH-antigen distance restraints. The restraint-guided approach (blue) shows improved interface positioning compared to unconstrained prediction (red), with backbone RMSD of 0.6 Å validating the methodology.

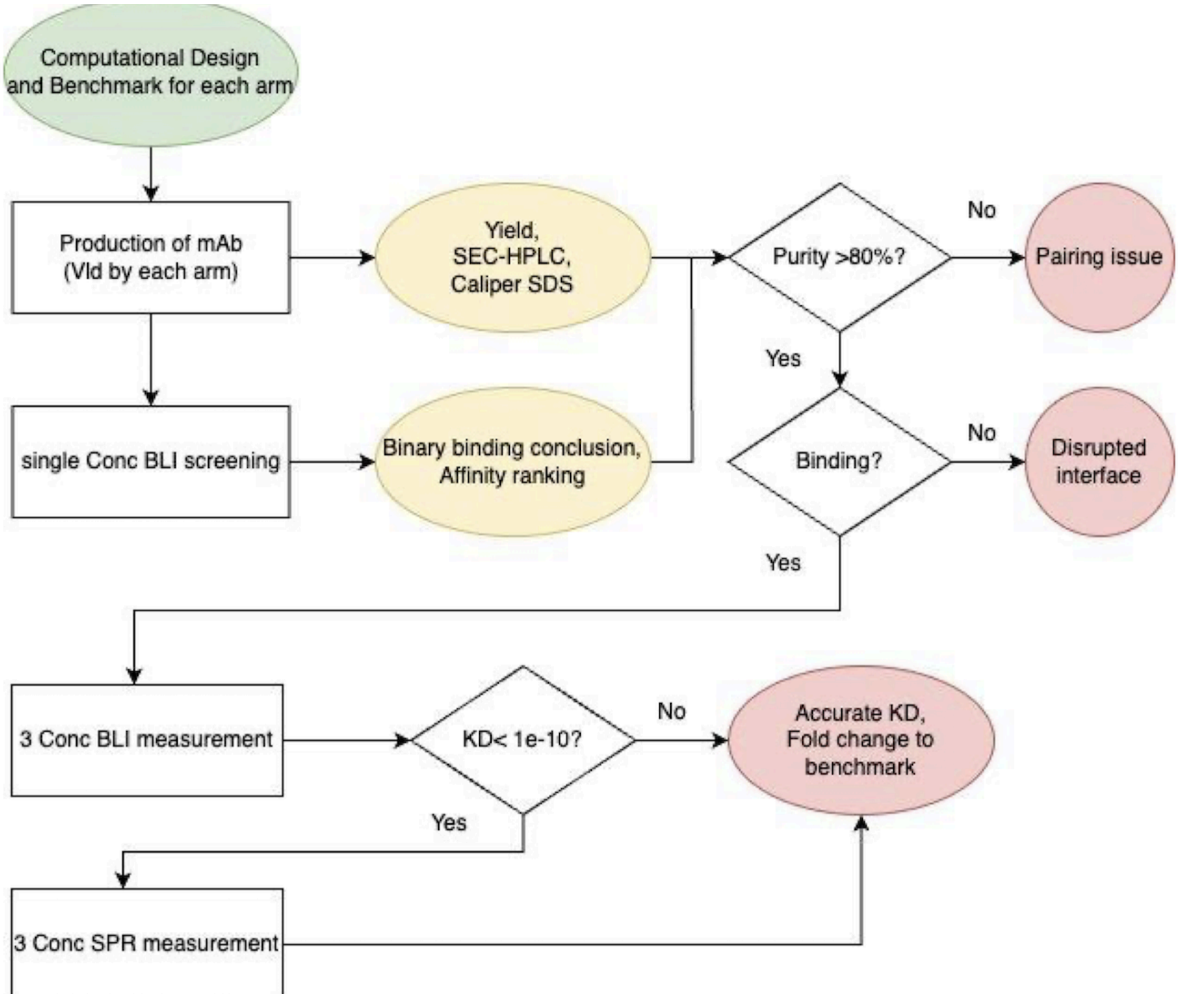

**Supplementary Figure S3. Experimental validation workflow.** Decision tree flowchart from computational design through expression, purification, binding screening, and affinity characterization. Pink boxes indicate failure points; yellow ovals show analytical steps; green ovals indicate success criteria.

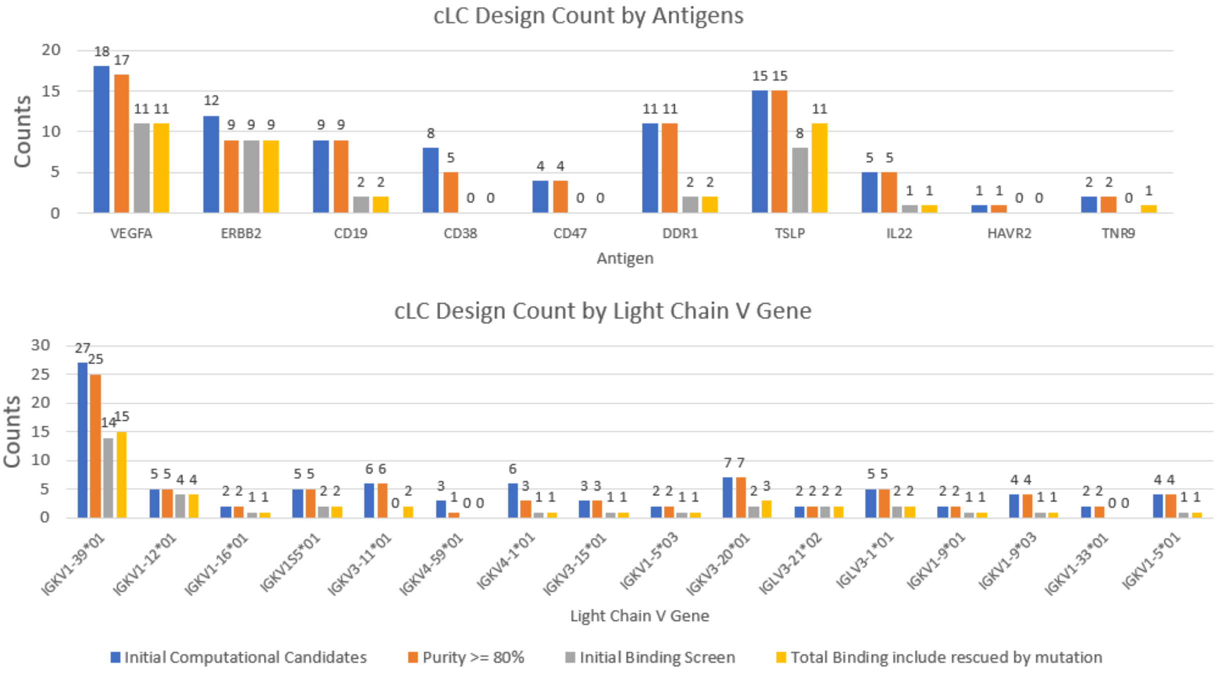

**Supplementary Figure S4. Distribution of designs across antigens and germline families.** (A) Design count by antigens showing distribution across 10 targets with binding success rates. (B) Light chain V gene usage patterns revealing IGKV1-39 enrichment in successful designs.

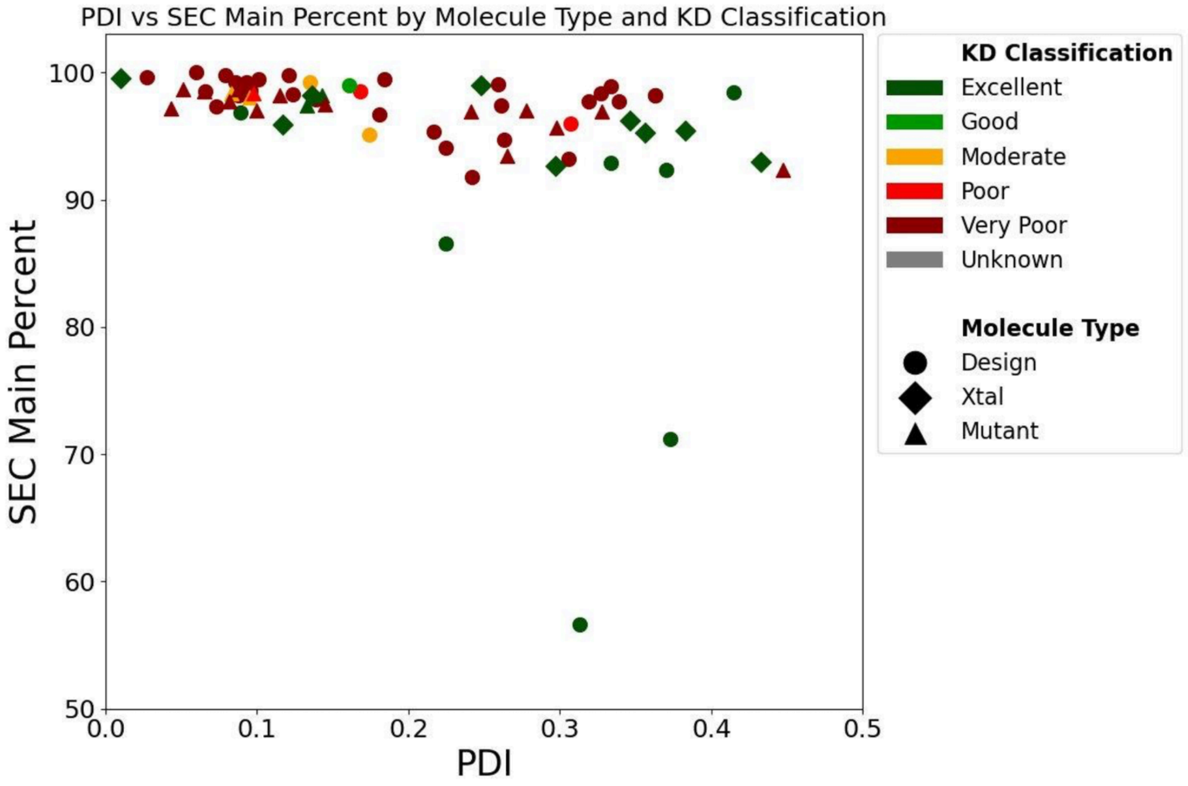

**Supplementary Figure S5. Quality metrics for validated common light chain designs.** PDI versus SEC main peak percentage scatter plot for all tested designs, colored by KD classification (excellent <3-fold, good 3-10-fold, moderate 10-30-fold, poor >30-fold). Design types indicated by symbols (circles: computational designs, diamonds: crystal structures, triangles: rescue mutants).

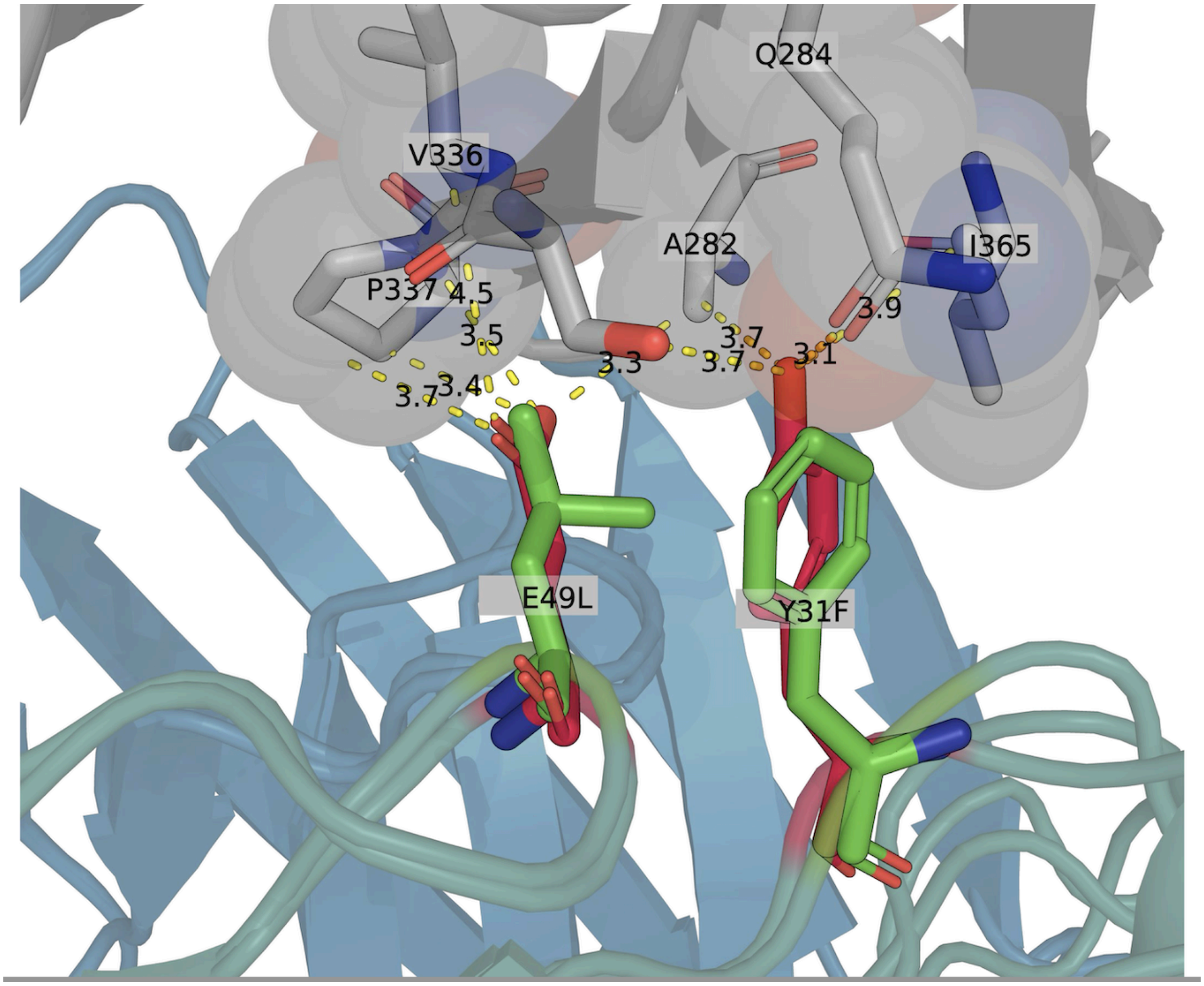

**Supplementary Figure S6. Molecular basis for structure-guided rescue mutations.** Structural visualization showing E49L mutation eliminating unfavorable buried charges within hydrophobic pockets (green: wild-type glutamate, red: mutated leucine) and Y31F mutation optimizing aromatic interactions. Distances shown in Angstroms.

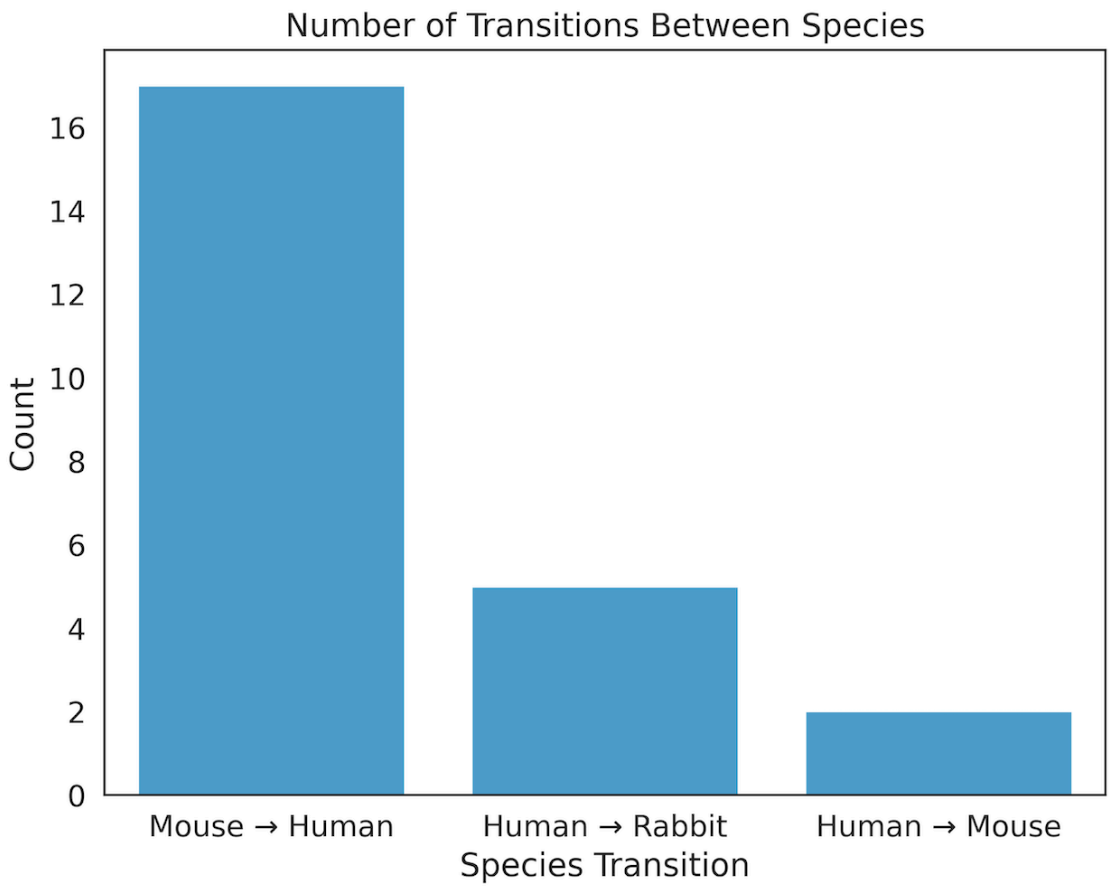

**Supplementary Figure S7. Species transitions in common light chain engineering.** Bar chart showing successful cross-species light chain conversions: mouse-to-human (n=17), human-to-rabbit (n=5), and human-to-mouse (n=2), demonstrating platform applicability for humanization efforts.

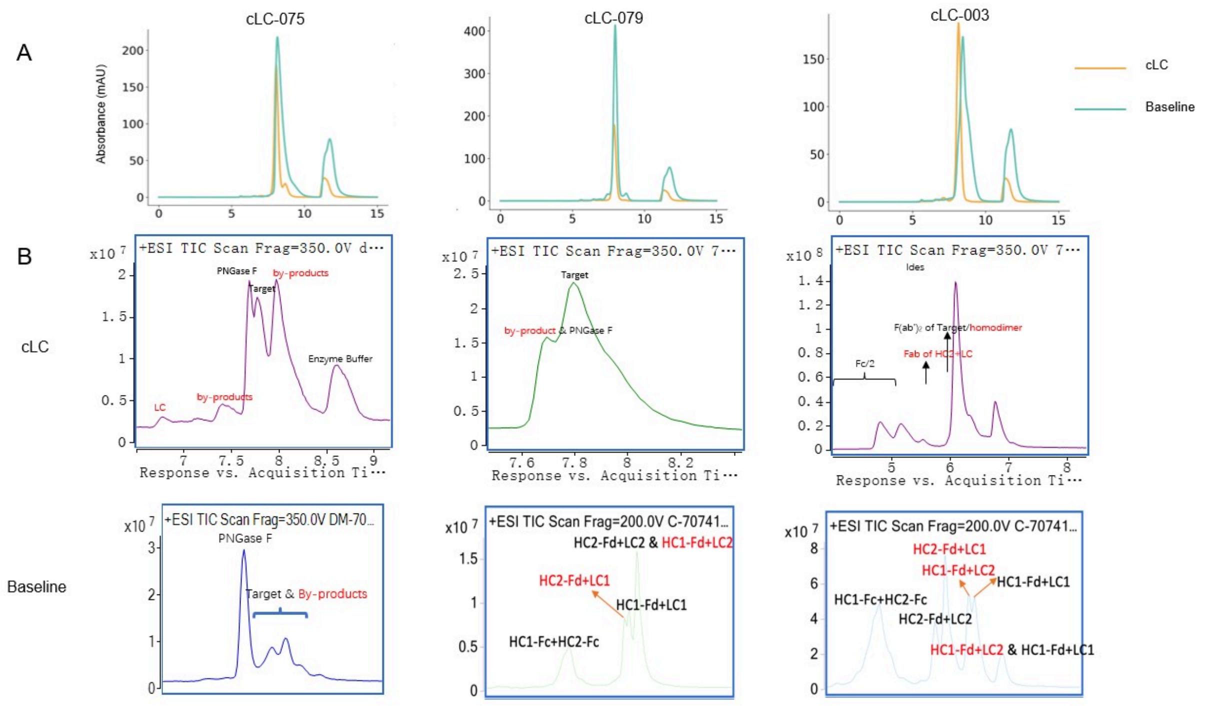

Supplementary Figure S8. **cLC improves bispecific assembly and purity across representative pairs (cLC-075, -079, -083)**. (A) SEC profiles: cLC constructs exhibit a higher main-peak fraction with reduced aggregate and low-molecular-weight species compared with baseline builds. (B) Intact-mass TIC scans: baseline shows multiple mispaired species (e.g., heavy-chain homodimers and half-antibody), whereas cLC yields a predominant intended heterotetramer; annotated masses indicate HC1/HC2/LC combinations, with off-target species flagged in the baseline panels.

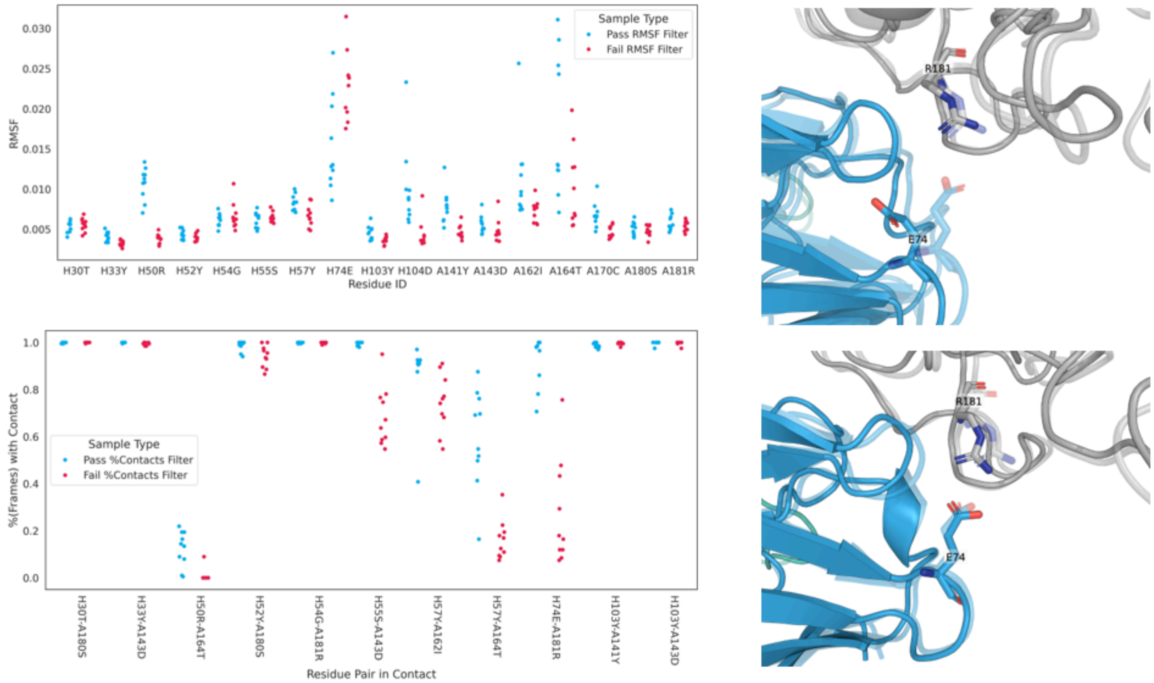

**Supplementary Figure S9. Molecular dynamics trajectory analysis.** (To be generated) RMSF distributions, contact persistence maps, and interaction stability metrics across 10 independent 6 ns simulations for representative successful designs.

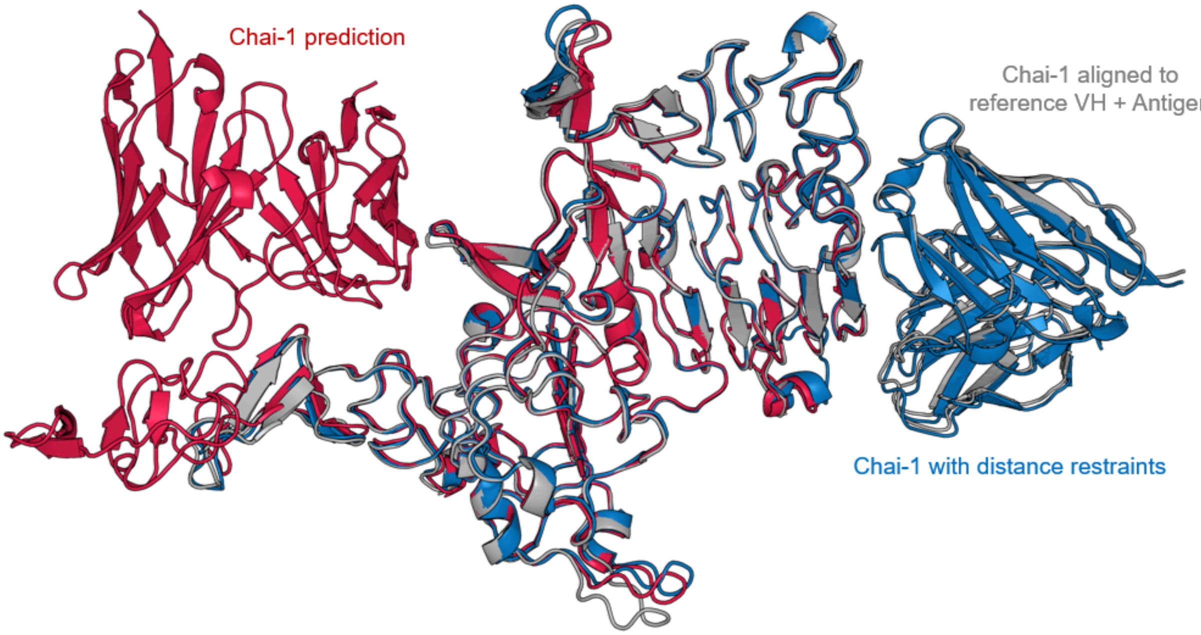

Supplementary Figure S10 **Structural validation of predicted antibody-antigen complexes.** (To be generated) Superposition of Chai-1 predicted structures with experimental crystal structures for validated designs, showing backbone RMSD values and interface preservation.

### Supplementary Tables

| Validation Tier | VH ID (PDB:Chain) | Target | Antigen Class | Tested (n) | Passed Expression (n) | Binding (n) | Success (%) | Best KD (nM) | Fold Change |
|---|---|---|---|---|---|---|---|---|---|
| 2 | 5j13 | TSLP | Cytokine | 15 | 15.0 | 8.0 | 53.3 | 1.1 | 153.6 |
| 1 | 6bft | VEGFA | Growth Factor | 13 | 12.0 | 6.0 | 46.2 | 6.2 | 787.3 |
| 2 | 4ag4 | DDR1 | Receptor Tyrosine Kinase | 11 | 11.0 | 2.0 | 18.2 | 21.4 | 10.4 |
| 1 | 5o4g | ERBB2 | Receptor Tyrosine Kinase | 10 | 7.0 | 7.0 | 70.0 | 0.8 | 1.1 |
| 1 | 7duo | CD38 | Plasma cell marker | 8 | 5.0 | 0.0 | 0.0 | ND | ND |
| 2 | 3q1s | Z13-IL22-2 | Viral | 5 | 5.0 | 1.0 | 20.0 | 630 | 58.9 |

| | | | | | | | | | |
|---|---|---|---|---|---|---|---|---|---|
| 1 | 4zff | VEGFA | Growth Factor | 5 | 5.0 | 5.0 | 100.0 | 0.4 | 2.1 |
| 1 | 7urx | CD19 | B-cell marker | 5 | 5.0 | 0.0 | 0.0 | ND | ND |
| 1 | 6al5 | CD19 | B-cell marker | 4 | 4.0 | 2.0 | 50.0 | 32.8 | 124.7 |
| 1 | 7xjf | CD47 | Checkpoint | 4 | 4.0 | 0.0 | 0.0 | ND | ND |
| 2 | 5o4g | ERBB2 | Receptor Tyrosine Kinase | 2 | 2.0 | 2.0 | 100.0 | 0.7 | 0.9 |
| 2 | 8gye | TNR9 | Costimulatory | 2 | 2.0 | 0.0 | 0.0 | ND | ND |
| 2 | 7kql | HAVR2 | Checkpoint | 1 | 1.0 | 0.0 | 0.0 | ND | ND |
| 1 | 1sy6 | CD3EG | T-cell receptor (TCR) co-receptor | 0 | | | | | |
| 1 | 2fjg | VEGFA | Growth Factor | 0 | | | | | |
| 1 | 2uzi | RASH | Small GTPase (RAS family) | 0 | | | | | |
| 1 | 2vh5 | RASH | Small GTPase (RAS family) | 0 | | | | | |
| 1 | 3bn9 | ST14 | Serine Protease (Matriptase) | 0 | | | | | |
| 1 | 3eoa | ITAL | Integrin αL (LFA-1, CD11a) | 0 | | | | | |
| 1 | 3l5x | IL13 | Cytokine | 0 | | | | | |
| 1 | 3skj | THRB | Receptor Tyrosine Kinase | 0 | | | | | |
| 1 | 5e8e | THRB | Coagulation Protease | 0 | | | | | |
| 1 | 6ban | ROR1 | Receptor Tyrosine Kinase | 0 | | | | | |
| 1 | 6o39 | FZD5 | G-protein coupled receptor (GPCR) | 0 | | | | | |
| 1 | 7joo | ICOS | Costimulatory Receptor | 0 | | | | | |
| 1 | 7yv1 | RASK | Small GTPase | 0 | | | | | |

| | | | | |
|---|---|---|---|---|
| | | | (RAS family) | |
| 1 | 8f0l | CD3E | T-cell receptor (TCR) component | 0 |
| 1 | 8ol9 | ICOS | Coagulation Protease | 0 |
| 1 | 8pg0 | SO1B3 | Solute Carrier (SLC) Transporter | 0 |
| 1 | NA (Mosunetuzumab) | CD3E | T-cell receptor (TCR) component | 0 |
| 1 | NA (Odronextamab) | CD20 | B-cell marker | 0 |
| 1 | NA (Petosemtamab) | EGFR | Receptor Tyrosine Kinase | 0 |

Supplementary Table S1 — Target panel and validation summary.
Overview of all VH–antigen targets included in the study, grouped by validation tier. For each VH (PDB ID:chain) and target/antigen class, the table lists constructs tested, expression pass count, binders observed, per-target success rate, best KD (nM), and fold-change vs parent where applicable. Success (%) is calculated as Binding(n)/Tested(n)×100.
Abbreviations: KD, equilibrium dissociation constant; PDB, Protein Data Bank.

| Design ID | SEC Main Peak (%) | KD (M) | Target | VH ID (PDB:Chain) | VL ID (PDB:Chain) |
|---|---|---|---|---|---|
| T1-A1 | 86.21 | 8.01e-10 | VEGFA | 4zff:A | 2uzi:L |
| T1-A2 | 95.07 | 1.78e-09 | VEGFA | 4zff:A | 3bn9:C |
| T1-A3 | 94.09 | 7.66e-09 | VEGFA | 4zff:A | 3eoa:L |
| T1-A4 | 86.56 | 3.64e-10 | VEGFA | 4zff:A | 7yv1:L |
| T1-G1 | 71.21 | | ERBB2 | 5o4g:B | 2fjg:A |
| T1-G2 | 92.33 | 9.94e-10 | ERBB2 | 5o4g:B | 2vh5:L |
| T1-G3 | 98.22 | 1.05e-09 | ERBB2 | 5o4g:B | 3bn9:C |
| T1-G4 | 92.91 | 8.5e-10 | ERBB2 | 5o4g:B | 7xjf:B |
| T1-C1 | 99.22 | 5.46e-07 | CD19 | 6al5:H | 3bn9:C |
| T1-C2 | 99.67 | | CD19 | 6al5:H | 4zff:B |
| T1-C3 | 95.82 | | CD19 | 6al5:H | 6ban:A |
| T1-C4 | 84.81 | 3.28e-08 | CD19 | 6al5:H | 6o39:A |
| T1-F1 | 86.35 | | CD38 | 7duo:H | 3l5x:L |
| T1-E1 | 88.59 | | CD47 | 7xjf:A | 5o4g:A |
| T1-E2 | 96.13 | | CD47 | 7xjf:A | 6o39:A |

| | | | | | |
|---|---|---|---|---|---|
| T1-E3 | 89.02 | | CD47 | 7xjf:A | 7yv1:L |
| T1-G5 | 56.59 | | ERBB2 | 5o4g:B | 1sy6:L |
| T1-G6 | 84.69 | 8.07e-08 | ERBB2 | 5o4g:B | 8f0l:L |
| T1-G7 | 74.07 | | ERBB2 | 5o4g:B | Mosunetuzumab |
| T1-F2 | 75.7 | | CD38 | 7duo:H | 1sy6:L |
| T1-F3 | 75.14 | | CD38 | 7duo:H | 8f0l:L |
| T1-F4 | 67.9 | | CD38 | 7duo:H | Mosunetuzumab |
| T1-D1 | 94.86 | | CD19 | 7urx:H | 1sy6:L |
| T1-D2 | 87.35 | | CD19 | 7urx:H | 8f0l:L |
| T1-D3 | 84.08 | | CD19 | 7urx:H | Mosunetuzumab |
| T1-G8 | 98.42 | 1.23e-09 | ERBB2 | 5o4g:B | Odronextamab |
| T1-F5 | 86.66 | | CD38 | 7duo:H | Odronextamab |
| T1-F6 | 81.6 | | CD38 | 7duo:H | Petosemtamab |
| T1-D4 | 91.62 | | CD19 | 7urx:H | Odronextamab |
| T1-D5 | 91.5 | | CD19 | 7urx:H | Petosemtamab |
| T1-A5 | 84.25 | 9.69e-09 | VEGFA | 4zff:A | 6ban:A |
| T1-G9 | 84.57 | 3.66e-09 | ERBB2 | 5o4g:B | 6al5:L |
| T1-G10 | 84.2 | 5.04e-07 | ERBB2 | 5o4g:B | 6ban:A |
| T1-B1 | 93.91 | 8.88e-09 | VEGFA | 6bft:H | 2uzi:L |
| T1-B2 | 94.62 | 6.22e-09 | VEGFA | 6bft:H | 2vh5:L |
| T1-B3 | 96.43 | 6.27e-09 | VEGFA | 6bft:H | 3bn9:C |
| T1-B4 | 94.57 | | VEGFA | 6bft:H | 3eoa:L |
| T1-B5 | 97.63 | | VEGFA | 6bft:H | 3skj:L |
| T1-B6 | 96.5 | | VEGFA | 6bft:H | 5o4g:A |
| T1-B7 | 92.9 | 8.76e-08 | VEGFA | 6bft:H | 6al5:L |
| T1-B8 | 87.91 | | VEGFA | 6bft:H | 6ban:A |
| T1-B9 | 68.97 | | VEGFA | 6bft:H | 6o39:A |
| T1-B10 | 92.41 | 4.2e-08 | VEGFA | 6bft:H | 7xjf:B |
| T1-B11 | 92.48 | | VEGFA | 6bft:H | 7yv1:L |
| T1-B12 | 93.96 | 3.36e-08 | VEGFA | 6bft:H | 8ol9:A |
| T1-B13 | 93.92 | | VEGFA | 6bft:H | 8pg0:L |
| T1-F7 | 93.65 | | CD38 | 7duo:H | 5e8e:A |
| T1-F8 | 97.53 | | CD38 | 7duo:H | 7joo:L |
| T1-E4 | 97.18 | | CD47 | 7xjf:A | 6ban:A |
| T2-K1 | 98.48 | 1.259e-07 | DDR1 | 4ag4:H | 3ulu:C |
| T2-M1 | 84.71 | 6.11e-09 | TSLP | 5j13:C | 3ulu:C |
| T2-I1 | 97.69 | | Z13-IL22-2 | 3q1s:H | 5bv7:L |
| T2-M2 | 94.72 | | TSLP | 5j13:C | 5bv7:B |
| T2-K2 | 93.25 | | DDR1 | 4ag4:H | 5lbs:L |
| T2-M3 | 85.29 | | TSLP | 5j13:C | 5lbs:L |
| T2-K3 | 91.79 | | DDR1 | 4ag4:H | 5yoy:F |

| | | | | | |
|---|---|---|---|---|---|
| T2-M4 | 97.7 | | TSLP | 5j13:C | 5yoy:F |
| T2-M5 | 98.23 | | TSLP | 5j13:C | 6nms:L |
| T2-L1 | 96.83 | 7.013e-10 | ERBB2 | 5o4g:B | 6nms:L |
| T2-K4 | 86.95 | | DDR1 | 4ag4:H | 6q1z:L |
| T2-M6 | 95.34 | 3.91e-08 | TSLP | 5j13:C | 6q1z:L |
| T2-K5 | 97.9 | | DDR1 | 4ag4:H | 7nx8:L |
| T2-M7 | 99.59 | 3.49e-08 | TSLP | 5j13:C | 7nx8:L |
| T2-K6 | 87.56 | | DDR1 | 4ag4:H | 7wnb:L |
| T2-M8 | 99.49 | 8.381e-09 | TSLP | 5j13:C | 7wnb:L |
| T2-J1 | 98.59 | | HAVR2 | 7kql:H | 7wnb:L |
| T2-H1 | 94.11 | | TNR9 | 8gye:E | 7wnb:L |
| T2-K7 | 97.31 | | DDR1 | 4ag4:H | 7y0w:L |
| T2-M9 | 100.0 | | TSLP | 5j13:C | 7y0w:B |
| T2-I2 | 98.22 | | Z13-IL22-2 | 3q1s:H | 7zfe:L |
| T2-M10 | 99.07 | 1.12e-09 | TSLP | 5j13:C | 7zfe:L |
| T2-K8 | 98.31 | | DDR1 | 4ag4:H | 7zrc:K |
| T2-M11 | 99.79 | 7.867e-09 | TSLP | 5j13:C | 7zrc:K |
| T2-I3 | 98.88 | | Z13-IL22-2 | 3q1s:H | 8c89:L |
| T2-M12 | 99.79 | | TSLP | 5j13:C | 8c89:L |
| T2-I4 | 98.36 | | Z13-IL22-2 | 3q1s:H | 8dlr:L |
| T2-K9 | 99.24 | 2.137e-08 | DDR1 | 4ag4:H | 8dlr:L |
| T2-L2 | 99.02 | 5.161e-09 | ERBB2 | 5o4g:B | 8dlr:L |
| T2-K10 | 99.19 | | DDR1 | 4ag4:H | 8dyy:B |
| T2-M13 | 97.44 | | TSLP | 5j13:C | 8dyy:B |
| T2-I5 | 96.01 | 6.3e-07 | Z13-IL22-2 | 3q1s:H | 8tnj:C |
| T2-M14 | 99.5 | 5.261e-08 | TSLP | 5j13:C | 8tnj:C |
| T2-H2 | 98.54 | | TNR9 | 8gye:E | 8tnj:C |
| T2-K11 | 96.69 | | DDR1 | 4ag4:H | 8vqd:B |
| T2-M15 | 99.26 | 1.877e-09 | TSLP | 5j13:C | 8vqd:B |

Supplementary Table S2 — Monospecific arm designs: expression, binding, and provenance.
Per-design metrics for single-arm constructs prior to bispecific assembly, including SEC main peak (%), measured KD (M), target, and the structural provenance (VH/VL PDB IDs:chains) used as inputs. This table enumerates the 85 monospecific designs underlying downstream cLC optimization.
Abbreviations: SEC, size-exclusion chromatography; KD, equilibrium dissociation constant.

| cLC ID | Validation Tier | Arm1 Design ID | Arm2 Design ID (or VH id from S1) | Arm1 Target | Arm2 Target |
|---|---|---|---|---|---|
| cLC-001 | 1 | T1-A1 | 2uzi | VEGFA | RASH |
| cLC-002 | 1 | T1-A2 | 3bn9 | VEGFA | ST14 |
| cLC-003 | 1 | T1-A3 | 3eoa | VEGFA | ITAL |
| cLC-004 | 1 | T1-A4 | 7yv1 | VEGFA | RASK |

| | | | | | |
|---|---|---|---|---|---|
| cLC-005 | 1 | T1-G1 | 2fjg | ERBB2 | VEGFA |
| cLC-006 | 1 | T1-G2 | 2vh5 | ERBB2 | RASH |
| cLC-007 | 1 | T1-G3 | 3bn9 | ERBB2 | ST14 |
| cLC-008 | 1 | T1-G4 | T1-Ee | ERBB2 | CD47 |
| cLC-009 | 1 | T1-C1 | 3bn9 | CD19 | ST14 |
| cLC-010 | 1 | T1-C2 | T1-Aa | CD19 | VEGFA |
| cLC-011 | 1 | T1-C3 | 6ban | CD19 | ROR1 |
| cLC-012 | 1 | T1-C4 | 6o39 | CD19 | FZD5 |
| cLC-013 | 1 | T1-F1 | 3l5x | CD38 | IL13 |
| cLC-014 | 1 | T1-E1 | T1-Gg | CD47 | ERBB2 |
| cLC-015 | 1 | T1-E2 | 6o39 | CD47 | FZD5 |
| cLC-016 | 1 | T1-E3 | 7yv1 | CD47 | RASK |
| cLC-017 | 1 | T1-G5 | 1sy6 | ERBB2 | CD3EG |
| cLC-018 | 1 | T1-G6 | 8f0l | ERBB2 | CD3E |
| cLC-019 | 1 | T1-G7 | mosunetuzumab | ERBB2 | CD3E |
| cLC-020 | 1 | T1-F2 | 1sy6 | CD38 | CD3EG |
| cLC-021 | 1 | T1-F3 | 8f0l | CD38 | CD3E |
| cLC-022 | 1 | T1-F4 | mosunetuzumab | CD38 | CD3E |
| cLC-023 | 1 | T1-D1 | 1sy6 | CD19 | CD3EG |
| cLC-024 | 1 | T1-D2 | 8f0l | CD19 | CD3E |
| cLC-025 | 1 | T1-D3 | mosunetuzumab | CD19 | CD3E |
| cLC-026 | 1 | T1-G8 | Odronextamab | ERBB2 | CD20 |
| cLC-027 | 1 | T1-F5 | Odronextamab | CD38 | CD20 |
| cLC-028 | 1 | T1-F6 | Petosemtamab | CD38 | EGFR |
| cLC-029 | 1 | T1-D4 | Odronextamab | CD19 | CD20 |
| cLC-030 | 1 | T1-D5 | Petosemtamab | CD19 | EGFR |
| cLC-031 | 1 | T1-A5 | 6ban | VEGFA | ROR1 |
| cLC-032 | 1 | T1-G9 | T1-Cc | ERBB2 | CD19 |
| cLC-033 | 1 | T1-G10 | 6ban | ERBB2 | ROR1 |
| cLC-034 | 1 | T1-B1 | 2uzi | VEGFA | RASH |
| cLC-035 | 1 | T1-B2 | 2vh5 | VEGFA | RASH |
| cLC-036 | 1 | T1-B3 | 3bn9 | VEGFA | ST14 |
| cLC-037 | 1 | T1-B4 | 3eoa | VEGFA | ITAL |
| cLC-038 | 1 | T1-B5 | 3skj | VEGFA | THRB |
| cLC-039 | 1 | T1-B6 | T1-Gg | VEGFA | ERBB2 |
| cLC-040 | 1 | T1-B7 | T1-Cc | VEGFA | CD19 |
| cLC-041 | 1 | T1-B8 | 6ban | VEGFA | ROR1 |
| cLC-042 | 1 | T1-B9 | 6o39 | VEGFA | FZD5 |
| cLC-043 | 1 | T1-B10 | T1-Ee | VEGFA | CD47 |
| cLC-044 | 1 | T1-B11 | 7yv1 | VEGFA | RASK |
| cLC-045 | 1 | T1-B12 | 8ol9 | VEGFA | ICOS |

| | | | | | |
|---|---|---|---|---|---|
| cLC-046 | 1 | T1-B13 | 8pg0 | VEGFA | SO1B3 |
| cLC-047 | 1 | T1-F7 | 5e8e | CD38 | THRB |
| cLC-048 | 1 | T1-F8 | 7joo | CD38 | ICOS |
| cLC-049 | 1 | T1-E4 | 6ban | CD47 | ROR1 |
| cLC-050 | 2 | T1-B1 | T1-A1 | VEGFA | VEGFA |
| cLC-051 | 2 | T1-G3 | T1-A2 | ERBB2 | VEGFA |
| cLC-052 | 2 | T1-C1 | T1-A2 | CD19 | VEGFA |
| cLC-053 | 2 | T1-B3 | T1-A2 | VEGFA | VEGFA |
| cLC-054 | 2 | T1-B4 | T1-A3 | VEGFA | VEGFA |
| cLC-055 | 2 | T1-E3 | T1-A4 | CD47 | VEGFA |
| cLC-056 | 2 | T1-B11 | T1-A4 | VEGFA | VEGFA |
| cLC-057 | 2 | T1-B2 | T1-G2 | VEGFA | ERBB2 |
| cLC-058 | 2 | T1-C1 | T1-G3 | CD19 | ERBB2 |
| cLC-059 | 2 | T1-B3 | T1-G3 | VEGFA | ERBB2 |
| cLC-060 | 2 | T1-B10 | T1-G4 | VEGFA | ERBB2 |
| cLC-061 | 2 | T1-B3 | T1-C1 | VEGFA | CD19 |
| cLC-062 | 2 | T1-C3 | T1-A5 | CD19 | VEGFA |
| cLC-063 | 2 | T1-C3 | T1-G10 | CD19 | ERBB2 |
| cLC-064 | 2 | T1-B8 | T1-C3 | VEGFA | CD19 |
| cLC-065 | 2 | T1-E4 | T1-C3 | CD47 | CD19 |
| cLC-066 | 2 | T1-E2 | T1-C4 | CD47 | CD19 |
| cLC-067 | 2 | T1-B9 | T1-C4 | VEGFA | CD19 |
| cLC-068 | 2 | T1-E1 | T1-F6 | CD47 | CD38 |
| cLC-069 | 2 | T1-E1 | T1-D5 | CD47 | CD19 |
| cLC-070 | 2 | T1-E1 | T1-B6 | CD47 | VEGFA |
| cLC-071 | 2 | T1-E2 | T1-B9 | CD47 | VEGFA |
| cLC-072 | 2 | T1-E3 | T1-B11 | CD47 | VEGFA |
| cLC-073 | 2 | T1-F2 | T1-G5 | CD38 | ERBB2 |
| cLC-074 | 2 | T1-D1 | T1-G5 | CD19 | ERBB2 |
| cLC-075 | 2 | T1-F3 | T1-G6 | CD38 | ERBB2 |
| cLC-076 | 2 | T1-D2 | T1-G6 | CD19 | ERBB2 |
| cLC-077 | 2 | T1-F4 | T1-G7 | CD38 | ERBB2 |
| cLC-078 | 2 | T1-D3 | T1-G7 | CD19 | ERBB2 |
| cLC-079 | 2 | T1-D1 | T1-F2 | CD19 | CD38 |
| cLC-080 | 2 | T1-D2 | T1-F3 | CD19 | CD38 |
| cLC-081 | 2 | T1-D3 | T1-F4 | CD19 | CD38 |
| cLC-082 | 2 | T1-F5 | T1-G8 | CD38 | ERBB2 |
| cLC-083 | 2 | T1-D4 | T1-G8 | CD19 | ERBB2 |
| cLC-084 | 2 | T1-D4 | T1-F5 | CD19 | CD38 |
| cLC-085 | 2 | T1-D5 | T1-F6 | CD19 | CD38 |
| cLC-086 | 2 | T1-F6 | T1-B6 | CD38 | VEGFA |

| | | | | | |
|---|---|---|---|---|---|
| cLC-087 | 2 | T1-D5 | T1-B6 | CD19 | VEGFA |
| cLC-088 | 2 | T1-G10 | T1-A5 | ERBB2 | VEGFA |
| cLC-089 | 2 | T1-B8 | T1-A5 | VEGFA | VEGFA |
| cLC-090 | 2 | T1-E4 | T1-A5 | CD47 | VEGFA |
| cLC-091 | 2 | T1-B7 | T1-G9 | VEGFA | ERBB2 |
| cLC-092 | 2 | T1-B8 | T1-G10 | VEGFA | ERBB2 |
| cLC-093 | 2 | T1-E4 | T1-G10 | CD47 | ERBB2 |
| cLC-094 | 2 | T1-E4 | T1-B8 | CD47 | VEGFA |
| cLC-095 | 2 | T2-M1 | T2-K1 | TSLP | DDR1 |
| cLC-096 | 2 | T2-M2 | T2-I1 | TSLP | Z13-IL22-2 |
| cLC-097 | 2 | T2-M3 | T2-K2 | TSLP | DDR1 |
| cLC-098 | 2 | T2-M4 | T2-K3 | TSLP | DDR1 |
| cLC-099 | 2 | T2-L1 | T2-M5 | ERBB2 | TSLP |
| cLC-100 | 2 | T2-M6 | T2-K4 | TSLP | DDR1 |
| cLC-101 | 2 | T2-M7 | T2-K5 | TSLP | DDR1 |
| cLC-102 | 2 | T2-M8 | T2-K6 | TSLP | DDR1 |
| cLC-103 | 2 | T2-J1 | T2-K6 | HAVR2 | DDR1 |
| cLC-104 | 2 | T2-H1 | T2-K6 | TNR9 | DDR1 |
| cLC-105 | 2 | T2-J1 | T2-M8 | HAVR2 | TSLP |
| cLC-106 | 2 | T2-H1 | T2-M8 | TNR9 | TSLP |
| cLC-107 | 2 | T2-H1 | T2-J1 | TNR9 | HAVR2 |
| cLC-108 | 2 | T2-M9 | T2-K7 | TSLP | DDR1 |
| cLC-109 | 2 | T2-M10 | T2-I2 | TSLP | Z13-IL22-2 |
| cLC-110 | 2 | T2-M11 | T2-K8 | TSLP | DDR1 |
| cLC-111 | 2 | T2-M12 | T2-I3 | TSLP | Z13-IL22-2 |
| cLC-112 | 2 | T2-K9 | T2-I4 | DDR1 | Z13-IL22-2 |
| cLC-113 | 2 | T2-L2 | T2-I4 | ERBB2 | Z13-IL22-2 |
| cLC-114 | 2 | T2-L2 | T2-K9 | ERBB2 | DDR1 |
| cLC-115 | 2 | T2-M13 | T2-K10 | TSLP | DDR1 |
| cLC-116 | 2 | T2-M14 | T2-I5 | TSLP | Z13-IL22-2 |
| cLC-117 | 2 | T2-H2 | T2-I5 | TNR9 | Z13-IL22-2 |
| cLC-118 | 2 | T2-H2 | T2-M14 | TNR9 | TSLP |
| cLC-119 | 2 | T2-M15 | T2-K11 | TSLP | DDR1 |

Supplementary Table S3 — cLC build matrix and per-arm outcomes.
Complete mapping of cLC builds linking Arm1/Arm2 design IDs (or originating VH IDs), targets, and per-arm design outcomes alongside the overall cLC outcome. “cLC Success” indicates that the common light chain supported productive binding across both arms under the study’s pass criteria.
Columns: cLC ID, Validation Tier, Arm1 Design ID, Arm2 Design ID (or VH from S1), Arm1/Arm2 Targets, Arm1/Arm2 Design Success, cLC Success.

| Design ID | Seed Desi | Mutation | Optimization | Binding | SEC Main Percent | KD (M) | Target | Original Desi | Seed KD (M) | KD Fold Change |
|---|---|---|---|---|---|---|---|---|---|---|

| | gn ID | | Success | Category | | | | gn Success | | |
|---|---|---|---|---|---|---|---|---|---|---|
| T2-K1-1 | T2-K1 | E49L_L; E52N_L | Yes | Improved | 89.81374275411373 | 1.4e-09 | DDR1 | Yes | 1.259e-07 | 0.01111993645750596 |
| T2-M1-1 | T2-M1 | E49L_L; E52N_L | No | Lost | 79.61467158376972 | | TSLP | Yes | 6.11e-09 | |
| T2-K1-2 | T2-K1 | E49L_L; E52N_L; Y31F_L | Yes | Improved | 98.19523361122148 | 8.76e-10 | DDR1 | Yes | 1.259e-07 | 0.006957903097696584 |
| T2-M1-2 | T2-M1 | E49L_L; E52N_L; Y31F_L | Yes | Maintained | 81.48562297564578 | 2.84e-08 | TSLP | Yes | 6.11e-09 | 4.648117839607202 |
| T2-K1-3 | T2-K1 | E49L_L; Y31F_L | Yes | Improved | 97.43501819988208 | 6.45e-10 | DDR1 | Yes | 1.259e-07 | 0.005123113582208101 |
| T2-M1-3 | T2-M1 | E49L_L; Y31F_L | Yes | Maintained | 85.72533251329571 | 2.06e-08 | TSLP | Yes | 6.11e-09 | 3.37152209492635 |
| T2-K1-4 | T2-K1 | E52N_L | Yes | Maintained | 98.3665669390411 | 1.65e-07 | DDR1 | Yes | 1.259e-07 | 1.310563939634631 |
| T2-M1-4 | T2-M1 | E52N_L | Yes | Maintained | 81.04572153952017 | 1.34e-08 | TSLP | Yes | 6.11e-09 | 2.193126022913257 |
| T2-K1-5 | T2-K1 | E52N_L; Y31F_L | Yes | Improved | 98.35871651431984 | 7.95e-08 | DDR1 | Yes | 1.259e-07 | 0.6314535345512311 |
| T2-M1-5 | T2-M1 | E52N_L; Y31F_L | Yes | Maintained | 81.14260422779726 | 9.07e-09 | TSLP | Yes | 6.11e-09 | 1.484451718494272 |
| T2-K1-6 | T2-K1 | Y31F_L | Yes | Improved | 98.0658618629556 | 3.09e-08 | DDR1 | Yes | 1.259e-07 | 0.2454328832406672 |
| T2-M1-6 | T2-M1 | Y31F_L | Yes | Improved | 84.13947888764058 | 5.37e-09 | TSLP | Yes | 6.11e-09 | 0.8788870703764321 |
| T2-K2-1 | T2-K2 | T53N_L | No | Nonbinding | 93.44071581300852 | | DDR1 | No | | |
| T2-M3-1 | T2-M3 | T53N_L | Yes | Rescued | 85.24609408189345 | 8.65e-08 | TSLP | No | | |
| T2-K3-1 | T2-K3 | W94N_L | No | Nonbinding | 89.4188942290084 | | DDR1 | No | | |
| T2-M4-1 | T2-M4 | W94N_L | Yes | Rescued | 96.89784953733157 | 4.74e-08 | TSLP | No | | |
| T2-K3-2 | T2-K3 | Y30T_L; W94N_L | No | Nonbinding | 82.81669344699371 | | DDR1 | No | | |
| T2-M4-2 | T2-M4 | Y30T_L; W94N_L | Yes | Rescued | 95.65464782321448 | 1.84e-07 | TSLP | No | | |
| T2-K3-3 | T2-K3 | Y32F_L | No | Nonbinding | 61.39680406418295 | | DDR1 | No | | |
| T2-M4-3 | T2-M4 | Y32F_L | Yes | Rescued | 97.02051559419004 | 6.58e-08 | TSLP | No | | |

| | | | | | | | | | | |
|---|---|---|---|---|---|---|---|---|---|---|
| T2-K3-4 | T2-K3 | Y32F_L; W94N_L | No | Nonbinding | 84.01687078412301 | | DDR1 | No | | |
| T2-M4-4 | T2-M4 | Y32F_L; W94N_L | Yes | Rescued | 96.90196038335468 | 1.41e-07 | TSLP | No | | |
| T2-K3-5 | T2-K3 | Y32F_L; Y30T_L; W94N_L | No | Nonbinding | 83.67017679930521 | | DDR1 | No | | |
| T2-M4-5 | T2-M4 | Y32F_L; Y30T_L; W94N_L | Yes | Rescued | 92.30111538353692 | 5.09e-08 | TSLP | No | | |
| T2-K5-1 | T2-K5 | A50D_L | No | Nonbinding | 96.96971996114844 | | DDR1 | No | | |
| T2-M7-1 | T2-M7 | A50D_L | Yes | Improved | 97.2039395363182 | 5.77e-09 | TSLP | Yes | 3.49e-08 | 0.1653295128939828 |
| T2-K5-2 | T2-K5 | A50D_L; L95H_L | No | Nonbinding | 97.6936363236023 | | DDR1 | No | | |
| T2-M7-2 | T2-M7 | A50D_L; L95H_L | Yes | Improved | 98.49114365833309 | 5.48e-09 | TSLP | Yes | 3.49e-08 | 0.1570200573065902 |
| T2-K5-3 | T2-K5 | L95H_L | No | Nonbinding | 98.18233066570733 | | DDR1 | No | | |
| T2-M7-3 | T2-M7 | L95H_L | Yes | Improved | 98.68728480883712 | 2.87e-08 | TSLP | Yes | 3.49e-08 | 0.822349570200573 |
| T2-K5-4 | T2-K5 | S53N_L | No | Nonbinding | 97.44480685588154 | | DDR1 | No | | |
| T2-M7-4 | T2-M7 | S53N_L | Yes | Improved | 98.65360090795632 | 2.91e-08 | TSLP | Yes | 3.49e-08 | 0.8338108882521489 |
| T2-K7-1 | T2-K7 | D92S_L | No | Nonbinding | 96.887465967942 | | DDR1 | No | | |
| T2-M9-1 | T2-M9 | D92S_L | No | Nonbinding | 98.42825832023654 | | TSLP | No | | |
| T2-K7-2 | T2-K7 | D92S_L; D93S_L | No | Nonbinding | 96.24691400609206 | | DDR1 | No | | |
| T2-M9-2 | T2-M9 | D92S_L; D93S_L | No | Nonbinding | 92.86796565669736 | | TSLP | No | | |
| T2-K7-3 | T2-K7 | D93S_L | No | Nonbinding | 96.19878815028817 | | DDR1 | No | | |
| T2-M9-3 | T2-M9 | D93S_L | No | Nonbinding | 99.09637917881896 | | TSLP | No | | |
| T2-I2-1 | T2-I2 | R97H_L | No | Nonbinding | 95.63463734801708 | | Z13-IL22-2 | No | | |
| T2-M10-1 | T2-M10 | R97H_L | Yes | Maintained | 98.66859773666155 | 4.16e-09 | TSLP | Yes | 1.12e-09 | 3.714285714285714 |

| | | | | | | | | | | |
|---|---|---|---|---|---|---|---|---|---|---|
| T2-K8-1 | T2-K8 | Y94S_L | No | Nonbinding | 98.09639369772856 | | DDR1 | No | | |
| T2-M11-1 | T2-M11 | Y94S_L | Yes | Improved | 99.8102298901628 | 5.55e-09 | TSLP | Yes | 7.867e-09 | 0.7054785814160417 |
| T2-I4-1 | T2-I4 | C33A_L | No | Nonbinding | 96.25046261529936 | | Z13-IL22-2 | No | | |
| T2-K9-1 | T2-K9 | C33A_L | Yes | Improved | 99.24682157590117 | 1.14e-08 | DDR1 | Yes | 2.137e-08 | 0.5334581188582125 |
| T2-L2-1 | T2-L2 | C33A_L | Yes | Maintained | 98.02982771121572 | 1.2e-08 | ERBB2 | Yes | 5.161e-09 | 2.325130788606859 |
| T2-I4-2 | T2-I4 | C33S_L | No | Nonbinding | 96.84748860308092 | | Z13-IL22-2 | No | | |
| T2-K9-2 | T2-K9 | C33S_L | Yes | Improved | 99.4465013266391 | 1.9e-08 | DDR1 | Yes | 2.137e-08 | 0.8890968647636875 |
| T2-L2-2 | T2-L2 | C33S_L | Yes | Maintained | 97.73577947034664 | 1.03e-08 | ERBB2 | Yes | 5.161e-09 | 1.995737260220887 |
| T2-I4-3 | T2-I4 | Y31F_L | No | Nonbinding | 94.76212229710536 | | Z13-IL22-2 | No | | |
| T2-L2-3 | T2-L2 | Y31F_L | Yes | Maintained | 97.28349556117873 | 6.74e-09 | ERBB2 | Yes | 5.161e-09 | 1.305948459600853 |
| T2-I4-4 | T2-I4 | Y31F_L; C33A_L | No | Nonbinding | 95.48549888518056 | | Z13-IL22-2 | No | | |
| T2-L2-4 | T2-L2 | Y31F_L; C33A_L | Yes | Maintained | 98.36270053043 | 5.67e-09 | ERBB2 | Yes | 5.161e-09 | 1.098624297616741 |
| T2-I4-5 | T2-I4 | Y31F_L; C33S_L | No | Nonbinding | 96.21537046988811 | | Z13-IL22-2 | No | | |
| T2-L2-5 | T2-L2 | Y31F_L; C33S_L | Yes | Maintained | 97.96005166665093 | 9.75e-09 | ERBB2 | Yes | 5.161e-09 | 1.889168765743073 |
| T2-K10-1 | T2-K10 | Y94S_L | No | Nonbinding | 99.72695231111244 | | DDR1 | No | | |
| T2-M13-1 | T2-M13 | Y94S_L | No | Nonbinding | 99.31215501766869 | | TSLP | No | | |
| T2-M14-1 | T2-M14 | S50D_L | Yes | Improved | 99.54933900552788 | 1.52e-08 | TSLP | Yes | 5.261e-08 | 0.2889184565671926 |
| T2-H2-1 | T2-H2 | S50D_L | No | Nonbinding | 98.95367040946743 | | TNR9 | No | | |

| | | | | | | | | | | |
|---|---|---|---|---|---|---|---|---|---|---|
| T2-I5-1 | T2-I5 | W92S_L | No | Lost | 94.28843282661732 | | Z13-IL22-2 | Yes | 6.3e-07 | |
| T2-M14-2 | T2-M14 | W92S_L | No | Lost | 99.3177521739763 | | TSLP | Yes | 5.261e-08 | |
| T2-H2-2 | T2-H2 | W92S_L | Yes | Rescued | 98.40128130436052 | 2.28e-08 | TNR9 | No | | |
| T2-K11-1 | T2-K11 | L96H_L | No | Nonbinding | 96.86270623747492 | | DDR1 | No | | |
| T2-M15-1 | T2-M15 | L96H_L | Yes | Weak | 98.79423667621232 | 3.46e-08 | TSLP | Yes | 1.877e-09 | 18.43367075119872 |
| T2-K11-2 | T2-K11 | W93S_L | No | Nonbinding | 98.917208001164 | | DDR1 | No | | |
| T2-M15-2 | T2-M15 | W93S_L | Yes | Weak | 100.0 | 4.1e-08 | TSLP | Yes | 1.877e-09 | 21.84336707511987 |
| T2-M14-3 | T2-M14 | S50D_L; A32Y_L | Yes | Maintained | 100.0 | 3.75e-07 | TSLP | Yes | 5.261e-08 | 7.127922448203764 |
| T2-H2-3 | T2-H2 | S50D_L; A32Y_L | Yes | Rescued | 97.45 | 6.2e-07 | TNR9 | No | | |
| T2-M14-4 | T2-M14 | S50D_L; A32Y_L; W92S_L | Yes | Maintained | 99.18 | 3.75e-07 | TSLP | Yes | 5.261e-08 | 7.127922448203764 |
| T2-H2-4 | T2-H2 | S50D_L; A32Y_L; W92S_L | Yes | Rescued | 97.15 | 2.25e-07 | TNR9 | No | | |
| T2-M14-5 | T2-M14 | S50D_L; W92S_L | Yes | Maintained | 99.56 | 3.99e-07 | TSLP | Yes | 5.261e-08 | 7.584109484888804 |
| T2-H2-5 | T2-H2 | S50D_L; W92S_L | Yes | Rescued | 97.77 | 2.83e-07 | TNR9 | No | | |
| T2-K11-3 | T2-K11 | W93S_L; L96H_L | No | Nonbinding | 98.98 | | DDR1 | No | | |
| T2-M15-3 | T2-M15 | W93S_L; L96H_L | Yes | Weak | 100.0 | 3.01e-08 | TSLP | Yes | 1.877e-09 | 16.03622802344166 |
| T2-K1-7 | T2-K1 | F95H_L | Yes | Weak | 99.28 | 1.46e-06 | DDR1 | Yes | 1.259e-07 | 11.59650516282764 |
| T2-M1-7 | T2-M1 | F95H_L | Yes | Maintained | 86.5 | 9.22e-09 | TSLP | Yes | 6.11e-09 | 1.509001636661211 |
| T2-K1-8 | T2-K1 | S55A_L | Yes | Maintained | 99.13 | 1.05e-06 | DDR1 | Yes | 1.259e-07 | 8.339952343129466 |
| T2-M1-8 | T2-M1 | S55A_L | Yes | Maintained | 89.6 | 8.75e-09 | TSLP | Yes | 6.11e-09 | 1.432078559738134 |
| T2-K1-9 | T2-K1 | S55E_L | Yes | Maintained | 99.16 | 9.53e-07 | DDR1 | Yes | 1.259e-07 | 7.569499602859412 |

| | | | | | | | | | | |
|---|---|---|---|---|---|---|---|---|---|---|
| T2-M1-9 | T2-M1 | S55E_L | Yes | Maintained | 87.68 | 1.38e-08 | TSLP | Yes | 6.11e-09 | 2.258592471358429 |
| T2-K1-10 | T2-K1 | S55E_L; F95H_L | Yes | Maintained | 99.47 | 9.51e-07 | DDR1 | Yes | 1.259e-07 | 7.553613979348689 |
| T2-M1-10 | T2-M1 | S55E_L; F95H_L | Yes | Maintained | 92.01 | 1.38e-08 | TSLP | Yes | 6.11e-09 | 2.258592471358429 |
| T2-K1-11 | T2-K1 | S55E_L; Y30K_L | Yes | Maintained | 99.57 | 1.04e-06 | DDR1 | Yes | 1.259e-07 | 8.260524225575853 |
| T2-M1-11 | T2-M1 | S55E_L; Y30K_L | Yes | Maintained | 89.73 | 1.63e-08 | TSLP | Yes | 6.11e-09 | 2.667757774140753 |
| T2-K1-12 | T2-K1 | S55T_L | Yes | Weak | 99.2 | 1.27e-06 | DDR1 | Yes | 1.259e-07 | 10.08737092930897 |
| T2-M1-12 | T2-M1 | S55T_L | Yes | Maintained | 86.61 | 7.75e-09 | TSLP | Yes | 6.11e-09 | 1.268412438625205 |
| T2-M5-1 | T2-M5 | D93S_L; R94T_L; L97T_L; T98F_L | No | Nonbinding | 61.35 | | TSLP | No | | |
| T2-L1-1 | T2-L1 | D93S_L; R94T_L; L97T_L; T98F_L | No | Lost | 70.64 | | ERBB2 | Yes | 7.013e-10 | |
| T2-M5-2 | T2-M5 | L97del_L | Yes | Rescued | 82.84 | 9.38e-08 | TSLP | No | | |
| T2-L1-2 | T2-L1 | L97del_L | Yes | Maintained | 93.87 | 3.4e-09 | ERBB2 | Yes | 7.013e-10 | 4.848139170112647 |
| T2-M5-3 | T2-M5 | R94T_L; Y30S_L; D93S_L; P95S_L | Yes | Rescued | 91.4 | 1.47e-07 | TSLP | No | | |
| T2-L1-3 | T2-L1 | R94T_L; Y30S_L; D93S_L; P95S_L | Yes | Maintained | 96.27 | 3.64e-09 | ERBB2 | Yes | 7.013e-10 | 5.190360758591187 |
| T2-M5-4 | T2-M5 | R94T_L; Y30S_L; P95S_L | Yes | Rescued | 89.19 | 1.41e-07 | TSLP | No | | |
| T2-L1-4 | T2-L1 | R94T_L; Y30S_L; P95S_L | Yes | Maintained | 97.06 | 4.44e-09 | ERBB2 | Yes | 7.013e-10 | 6.331099386852987 |
| T2-M5-5 | T2-M5 | R94T_L; Y30S_L; Y92S_L; P95S_L | Yes | Rescued | 86.63 | 1.05e-07 | TSLP | No | | |
| T2-L1-5 | T2-L1 | R94T_L; Y30S_L; Y92S_L; P95S_L | Yes | Maintained | 94.28 | 5.96e-09 | ERBB2 | Yes | 7.013e-10 | 8.498502780550407 |
| T2-M5-6 | T2-M5 | Y30S_L | Yes | Rescued | 94.7 | 1e-08 | TSLP | No | | |

| | | | | | | | | | | |
|---|---|---|---|---|---|---|---|---|---|---|
| T2-L1-6 | T2-L1 | Y30S_L | Yes | Maintained | 86.56 | 5.39e-09 | ERBB2 | Yes | 7.013e-10 | 7.685726507913873 |
| T2-M5-7 | T2-M5 | Y30S_L; D93S_L | Yes | Rescued | 83.02 | 6.81e-08 | TSLP | No | | |
| T2-L1-7 | T2-L1 | Y30S_L; D93S_L | No | Lost | 77.26 | | ERBB2 | Yes | 7.013e-10 | |
| T2-M5-8 | T2-M5 | Y30S_L; D93S_L; L97T_L | Yes | Rescued | 83.91 | 2.89e-07 | TSLP | No | | |
| T2-L1-8 | T2-L1 | Y30S_L; D93S_L; L97T_L | Yes | Weak | 95.2 | 7.54e-09 | ERBB2 | Yes | 7.013e-10 | 10.75146157136746 |
| T2-M5-9 | T2-M5 | Y30S_L; D93S_L; P95S_L | Yes | Rescued | 91.41 | 4.27e-07 | TSLP | No | | |
| T2-L1-9 | T2-L1 | Y30S_L; D93S_L; P95S_L | Yes | Maintained | 94.59 | 6.08e-09 | ERBB2 | Yes | 7.013e-10 | 8.669613574789675 |
| T2-M5-10 | T2-M5 | Y30S_L; D93S_L; Y92S_L | Yes | Rescued | 91.28 | 1.93e-07 | TSLP | No | | |
| T2-L1-10 | T2-L1 | Y30S_L; D93S_L; Y92S_L | Yes | Maintained | 95.06 | 4.28e-09 | ERBB2 | Yes | 7.013e-10 | 6.102951661200627 |
| T2-M5-11 | T2-M5 | Y30S_L; D93S_L; Y92S_L; P95S_L | No | Nonbinding | 78.64 | | TSLP | No | | |
| T2-L1-11 | T2-L1 | Y30S_L; D93S_L; Y92S_L; P95S_L | Yes | Maintained | 94.91 | 4.96e-09 | ERBB2 | Yes | 7.013e-10 | 7.072579495223157 |
| T2-M5-12 | T2-M5 | Y30S_L; L97T_L | Yes | Rescued | 95.3 | 9.52e-09 | TSLP | No | | |
| T2-L1-12 | T2-L1 | Y30S_L; L97T_L | Yes | Weak | 96.16 | 7.76e-09 | ERBB2 | Yes | 7.013e-10 | 11.06516469413945 |
| T2-M5-13 | T2-M5 | Y30S_L; L97T_L; Y92S_L | Yes | Rescued | 87.33 | 4.54e-07 | TSLP | No | | |
| T2-L1-13 | T2-L1 | Y30S_L; L97T_L; Y92S_L | Yes | Weak | 91.71 | 1.21e-08 | ERBB2 | Yes | 7.013e-10 | 17.25367175245972 |
| T2-M5-14 | T2-M5 | Y30S_L; P95S_L | Yes | Rescued | 97.27 | 4.54e-07 | TSLP | No | | |
| T2-L1-14 | T2-L1 | Y30S_L; P95S_L | Yes | Maintained | 98.39 | 6.72e-09 | ERBB2 | Yes | 7.013e-10 | 9.582204477399115 |
| T2-M5-15 | T2-M5 | Y30S_L; Y92S_L | Yes | Rescued | 84.12 | 6.31e-08 | TSLP | No | | |
| T2-L1-15 | T2-L1 | Y30S_L; Y92S_L | No | Lost | 68.23 | | ERBB2 | Yes | 7.013e-10 | |

| | | | | | | | | | | |
|---|---|---|---|---|---|---|---|---|---|---|
| T2-M5-16 | T2-M5 | Y30S_L; Y92S_L; P95S_L | Yes | Rescued | 81.24 | 1.04e-07 | TSLP | No | | |
| T2-L1-16 | T2-L1 | Y30S_L; Y92S_L; P95S_L | No | Lost | 66.21 | | ERBB2 | Yes | 7.013e-10 | |
| T2-K5-5 | T2-K5 | A50S_L | No | Nonbinding | 98.61 | | DDR1 | No | | |
| T2-M7-5 | T2-M7 | A50S_L | Yes | Maintained | 99.46 | 3.75e-08 | TSLP | Yes | 3.49e-08 | 1.074498567335243 |
| | | 55.999999999999986 | | | | | | | | |

Supplementary Table S4 — Structure-guided LC optimization (single-arm rescue set).
Per-design record of structure-guided rescue campaigns, listing the seed design, specific LC mutations, optimization outcome, binding category, SEC main %, KD (M), target, original design success state, seed KD (M), and KD fold-change. Fold-change is reported as KD(seed)/KD(optimized), with >1 indicating affinity improvement. Abbreviations: SEC, size-exclusion chromatography; KD, equilibrium dissociation constant.

| cLC ID | Arm1 Design ID | Arm2 Design ID | Seed cLC ID | Mutation | Arm1 Target | Arm2 Target | Arm1 KD Fold Change | Arm2 KD Fold Change |
|---|---|---|---|---|---|---|---|---|
| cLC-120 | T2-M1-1 | T2-K1-1 | cLC-095 | E49L_L; E52N_L | TSLP | DDR1 | | 0.01111993645750596 |
| cLC-121 | T2-M1-2 | T2-K1-2 | cLC-095 | E49L_L; E52N_L; Y31F_L | TSLP | DDR1 | 4.648117839607202 | 0.006957903097696584 |
| cLC-122 | T2-M1-3 | T2-K1-3 | cLC-095 | E49L_L; Y31F_L | TSLP | DDR1 | 3.37152209492635 | 0.005123113582208101 |
| cLC-123 | T2-M1-4 | T2-K1-4 | cLC-095 | E52N_L | TSLP | DDR1 | 2.193126022913257 | 1.310563939634631 |
| cLC-124 | T2-M1-5 | T2-K1-5 | cLC-095 | E52N_L; Y31F_L | TSLP | DDR1 | 1.484451718494272 | 0.6314535345512311 |
| cLC-125 | T2-M1-6 | T2-K1-6 | cLC-095 | Y31F_L | TSLP | DDR1 | 0.8788870703764321 | 0.2454328832406672 |
| cLC-126 | T2-M3-1 | T2-K2-1 | cLC-097 | T53N_L | TSLP | DDR1 | | |
| cLC-127 | T2-M4-1 | T2-K3-1 | cLC-098 | W94N_L | TSLP | DDR1 | | |
| cLC-128 | T2-M4-2 | T2-K3-2 | cLC-098 | Y30T_L; W94N_L | TSLP | DDR1 | | |
| cLC-129 | T2-M4-3 | T2-K3-3 | cLC-098 | Y32F_L | TSLP | DDR1 | | |
| cLC-130 | T2-M4-4 | T2-K3-4 | cLC-098 | Y32F_L; W94N_L | TSLP | DDR1 | | |
| cLC-131 | T2-M4-5 | T2-K3-5 | cLC-098 | Y32F_L; Y30T_L; W94N_L | TSLP | DDR1 | | |
| cLC-132 | T2-M7-1 | T2-K5-1 | cLC-101 | A50D_L | TSLP | DDR1 | 0.1653295128939828 | |
| cLC-133 | T2-M7-2 | T2-K5-2 | cLC-101 | A50D_L; L95H_L | TSLP | DDR1 | 0.1570200573065902 | |
| cLC-134 | T2-M7-3 | T2-K5-3 | cLC-101 | L95H_L | TSLP | DDR1 | 0.822349570200573 | |

| | | | | | | | | |
|---|---|---|---|---|---|---|---|---|
| cLC-135 | T2-M7-4 | T2-K5-4 | cLC-101 | S53N_L | TSLP | DDR1 | 0.8338108882521489 | |
| cLC-136 | T2-M9-1 | T2-K7-1 | cLC-108 | D92S_L | TSLP | DDR1 | | |
| cLC-137 | T2-M9-2 | T2-K7-2 | cLC-108 | D92S_L; D93S_L | TSLP | DDR1 | | |
| cLC-138 | T2-M9-3 | T2-K7-3 | cLC-108 | D93S_L | TSLP | DDR1 | | |
| cLC-139 | T2-M10-1 | T2-I2-1 | cLC-109 | R97H_L | TSLP | Z13-IL22-2 | 3.714285714285714 | |
| cLC-140 | T2-M11-1 | T2-K8-1 | cLC-110 | Y94S_L | TSLP | DDR1 | 0.7054785814160417 | |
| cLC-141 | T2-K9-1 | T2-I4-1 | cLC-112 | C33A_L | DDR1 | Z13-IL22-2 | 0.5334581188582125 | |
| cLC-142 | T2-L2-1 | T2-I4-1 | cLC-113 | C33A_L | ERBB2 | Z13-IL22-2 | 2.325130788606859 | |
| cLC-143 | T2-L2-1 | T2-K9-1 | cLC-114 | C33A_L | ERBB2 | DDR1 | 2.325130788606859 | 0.5334581188582125 |
| cLC-144 | T2-K9-2 | T2-I4-2 | cLC-112 | C33S_L | DDR1 | Z13-IL22-2 | 0.8890968647636875 | |
| cLC-145 | T2-L2-2 | T2-I4-2 | cLC-113 | C33S_L | ERBB2 | Z13-IL22-2 | 1.995737260220887 | |
| cLC-146 | T2-L2-2 | T2-K9-2 | cLC-114 | C33S_L | ERBB2 | DDR1 | 1.995737260220887 | 0.8890968647636875 |
| cLC-147 | T2-L2-3 | T2-I4-3 | cLC-113 | Y31F_L | ERBB2 | Z13-IL22-2 | 1.305948459600853 | |
| cLC-148 | T2-L2-4 | T2-I4-4 | cLC-113 | Y31F_L; C33A_L | ERBB2 | Z13-IL22-2 | 1.098624297616741 | |
| cLC-149 | T2-L2-5 | T2-I4-5 | cLC-113 | Y31F_L; C33S_L | ERBB2 | Z13-IL22-2 | 1.889168765743073 | |
| cLC-150 | T2-M13-1 | T2-K10-1 | cLC-115 | Y94S_L | TSLP | DDR1 | | |
| cLC-151 | T2-H2-1 | T2-M14-1 | cLC-118 | S50D_L | TNR9 | TSLP | | 0.2889184565671926 |
| cLC-152 | T2-M14-2 | T2-I5-1 | cLC-116 | W92S_L | TSLP | Z13-IL22-2 | | |
| cLC-153 | T2-H2-2 | T2-I5-1 | cLC-117 | W92S_L | TNR9 | Z13-IL22-2 | | |
| cLC-154 | T2-H2-2 | T2-M14-2 | cLC-118 | W92S_L | TNR9 | TSLP | | |
| cLC-155 | T2-M15-1 | T2-K11-1 | cLC-119 | L96H_L | TSLP | DDR1 | 18.43367075119872 | |
| cLC-156 | T2-M15-2 | T2-K11-2 | cLC-119 | W93S_L | TSLP | DDR1 | 21.84336707511987 | |
| cLC-157 | T2-H2-3 | T2-M14-3 | cLC-118 | S50D_L; A32Y_L | TNR9 | TSLP | | 7.127922448203764 |
| cLC-158 | T2-H2-4 | T2-M14-4 | cLC-118 | S50D_L; A32Y_L; W92S_L | TNR9 | TSLP | | 7.127922448203764 |
| cLC-159 | T2-H2-5 | T2-M14-5 | cLC-118 | S50D_L; W92S_L | TNR9 | TSLP | | 7.584109484888804 |
| cLC-160 | T2-M15-3 | T2-K11-3 | cLC-119 | W93S_L; L96H_L | TSLP | DDR1 | 16.03622802344166 | |
| cLC-161 | T2-M1-7 | T2-K1-7 | cLC-095 | F95H_L | TSLP | DDR1 | 1.509001636661211 | 11.59650516282764 |
| cLC-162 | T2-M1-8 | T2-K1-8 | cLC-095 | S55A_L | TSLP | DDR1 | 1.432078559738134 | 8.339952343129466 |

| | | | | | | | | |
|---|---|---|---|---|---|---|---|---|
| cLC-163 | T2-M1-9 | T2-K1-9 | cLC-095 | S55E_L | TSLP | DDR1 | 2.258592471358429 | 7.569499602859412 |
| cLC-164 | T2-M1-10 | T2-K1-10 | cLC-095 | S55E_L; F95H_L | TSLP | DDR1 | 2.258592471358429 | 7.553613979348689 |
| cLC-165 | T2-M1-11 | T2-K1-11 | cLC-095 | S55E_L; Y30K_L | TSLP | DDR1 | 2.667757774140753 | 8.260524225575853 |
| cLC-166 | T2-M1-12 | T2-K1-12 | cLC-095 | S55T_L | TSLP | DDR1 | 1.268412438625205 | 10.08737092930897 |
| cLC-167 | T2-L1-1 | T2-M5-1 | cLC-099 | D93S_L; R94T_L; L97T_L; T98F_L | ERBB2 | TSLP | | |
| cLC-168 | T2-L1-2 | T2-M5-2 | cLC-099 | L97del_L | ERBB2 | TSLP | 4.848139170112647 | |
| cLC-169 | T2-L1-3 | T2-M5-3 | cLC-099 | R94T_L; Y30S_L; D93S_L; P95S_L | ERBB2 | TSLP | 5.190360758591187 | |
| cLC-170 | T2-L1-4 | T2-M5-4 | cLC-099 | R94T_L; Y30S_L; P95S_L | ERBB2 | TSLP | 6.331099386852987 | |
| cLC-171 | T2-L1-5 | T2-M5-5 | cLC-099 | R94T_L; Y30S_L; Y92S_L; P95S_L | ERBB2 | TSLP | 8.498502780550407 | |
| cLC-172 | T2-L1-6 | T2-M5-6 | cLC-099 | Y30S_L | ERBB2 | TSLP | 7.685726507913873 | |
| cLC-173 | T2-L1-7 | T2-M5-7 | cLC-099 | Y30S_L; D93S_L | ERBB2 | TSLP | | |
| cLC-174 | T2-L1-8 | T2-M5-8 | cLC-099 | Y30S_L; D93S_L; L97T_L | ERBB2 | TSLP | 10.75146157136746 | |
| cLC-175 | T2-L1-9 | T2-M5-9 | cLC-099 | Y30S_L; D93S_L; P95S_L | ERBB2 | TSLP | 8.669613574789675 | |
| cLC-176 | T2-L1-10 | T2-M5-10 | cLC-099 | Y30S_L; D93S_L; Y92S_L | ERBB2 | TSLP | 6.102951661200627 | |
| cLC-177 | T2-L1-11 | T2-M5-11 | cLC-099 | Y30S_L; D93S_L; Y92S_L; P95S_L | ERBB2 | TSLP | 7.072579495223157 | |
| cLC-178 | T2-L1-12 | T2-M5-12 | cLC-099 | Y30S_L; L97T_L | ERBB2 | TSLP | 11.06516469413945 | |
| cLC-179 | T2-L1-13 | T2-M5-13 | cLC-099 | Y30S_L; L97T_L; Y92S_L | ERBB2 | TSLP | 17.25367175245972 | |
| cLC-180 | T2-L1-14 | T2-M5-14 | cLC-099 | Y30S_L; P95S_L | ERBB2 | TSLP | 9.582204477399115 | |
| cLC-181 | T2-L1-15 | T2-M5-15 | cLC-099 | Y30S_L; Y92S_L | ERBB2 | TSLP | | |
| cLC-182 | T2-L1-16 | T2-M5-16 | cLC-099 | Y30S_L; Y92S_L; P95S_L | ERBB2 | TSLP | | |

| | | | | | | | |
|---|---|---|---|---|---|---|---|
| cLC-183 | T2-M7-5 | T2-K5-5 | cLC-101 | A50S_L | TSLP | DDR1 | 1.074498567335243 |

Supplementary Table S5 — cLC optimization across both arms and bispecific context.
Paired-arm view of cLC optimization, including cLC/seed IDs, mutations applied, Arm1/Arm2 targets, post-optimization binding categories per arm, overall cLC binding category/success, and per-arm KD fold-change relative to seed. This table links single-arm improvements to dual-arm (cLC) performance.
Abbreviations: KD, equilibrium dissociation constant.

| Arm1 Target | Arm2 Target | Original Design cLC Success | Optimized cLC Success |
|---|---|---|---|
| CD19 | CD20 | 0/1 | |
| CD19 | CD3E | 0/2 | |
| CD19 | CD3EG | 0/1 | |
| CD19 | EGFR | 0/1 | |
| CD19 | FZD5 | 1/1 | |
| CD19 | ROR1 | 0/1 | |
| CD19 | ST14 | 1/1 | |
| CD38 | CD20 | 0/1 | |
| CD38 | CD3E | 0/2 | |
| CD38 | CD3EG | 0/1 | |
| CD38 | EGFR | 0/1 | |
| CD38 | ICOS | 0/1 | |
| CD38 | IL13 | 0/1 | |
| CD38 | THRB | 0/1 | |
| CD47 | FZD5 | 0/1 | |
| CD47 | RASK | 0/1 | |
| CD47 | ROR1 | 0/1 | |
| ERBB2 | CD20 | 1/1 | |
| ERBB2 | CD3E | 1/2 | |
| ERBB2 | CD3EG | 0/1 | |
| ERBB2 | RASH | 1/1 | |
| ERBB2 | ROR1 | 1/1 | |
| ERBB2 | ST14 | 1/1 | |
| VEGFA | FZD5 | 0/1 | |
| VEGFA | ICOS | 1/1 | |
| VEGFA | ITAL | 1/2 | |
| VEGFA | RASH | 3/3 | |
| VEGFA | RASK | 1/2 | |
| VEGFA | ROR1 | 1/2 | |
| VEGFA | SO1B3 | 0/1 | |
| VEGFA | ST14 | 2/2 | |
| VEGFA | THRB | 0/1 | |
| CD19 | CD38 | 0/5 | |

| | | | |
|---|---|---|---|
| CD19 | ERBB2 | 2/7 | |
| CD38 | ERBB2 | 0/4 | |
| CD38 | VEGFA | 0/1 | |
| CD47 | CD19 | 0/3 | |
| CD47 | CD38 | 0/1 | |
| CD47 | ERBB2 | 1/3 | |
| CD47 | VEGFA | 1/7 | |
| DDR1 | Z13-IL22-2 | 0/1 | 0/2 |
| ERBB2 | DDR1 | 1/1 | 0/2 |
| ERBB2 | Z13-IL22-2 | 0/1 | 0/5 |
| ERBB2 | TSLP | 0/1 | 11/16 |
| HAVR2 | DDR1 | 0/1 | |
| HAVR2 | TSLP | 0/1 | |
| TNR9 | DDR1 | 0/1 | |
| TNR9 | HAVR2 | 0/1 | |
| TNR9 | Z13-IL22-2 | 0/1 | 0/1 |
| TNR9 | TSLP | 0/2 | 3/5 |
| TSLP | DDR1 | 1/10 | 3/31 |
| TSLP | Z13-IL22-2 | 1/4 | 0/2 |
| VEGFA | CD19 | 3/8 | |
| VEGFA | ERBB2 | 6/9 | |
| VEGFA | VEGFA | 2/5 | |
| | | 34/119 | 17/64 |

Supplementary Table S6 — Target-pair success rates before and after optimization.
Summary of cLC success across target pairs (Arm1 target × Arm2 target), reported as success rates for the original designs and after structure-guided optimization. This highlights target-pair-specific permissivity and the gains attributable to optimization.
Columns: Arm1 Target, Arm2 Target, Original Design cLC Success, Optimized cLC Success.

| Target | Success rate | PDB | Charged-AA involved contacts | Other polar-AA involved contacts |
|---|---|---|---|---|
| VEGFA | 61.1% (11/18) | 4zff | | L-Y89-OH/H-Q42-NE2 |
| | | | | L-S94-OG/H-T52-OG1 |
| | | | | L-Y87-OH/H-Q39-NE2 |
| | | | | L-Q38-NE2/H-Y91-OH |
| | | 6bft | | L-Q38-OE1/H-Q39-NE2 |
| | | | | L-Q38-NE2/H-Q39-OE1 |
| | | | | L-Y87-OH/H-Q39-NE2 |

| | | | | |
|---|---|---|---|---|
| | | | | L-Y91-OH/H-H107-ND1 |
| | | | | L-Q89-NE2/H-W108-O |
| ERBB2 | 75% (9/12) | 5o4g | | L-Q38-OE1/H-Q39-NE2 |
| | | | | L-Q38-NE2/H-Q39-OE1 |
| | | | | L-Y87-OH/H-Q39-NE2 |
| | | | | L-Q89-NE2/H-A106-O |
| CD19 | 22.2% (2/9) | 7urx (all no-binding) | L-Y115-OH/H-K118-NZ | L-Q57-OE1/H-Q58-NE2 |
| | | | L-S69-OG/H-D125-OD1 | L-Q57-NE2/H-Q58-OE1 |
| | | | L-K28-NZ/H-G61-O | L-Y115-OH/H-Q69-OE1 |
| | | | | L-N74-OD1/H-Y127-OH |
| | | | | L-S117-OG/H-L64-O |
| | | 6al5 | | L-Q42-NE2/H39-OE1 |
| | | | | L-Q42-OE1/H39-NE2 |
| | | | | L-Y31-OH/H-Y107-OH |
| | | | | L-N98-ND2/H-Y50-O |
| | | | | L-S95-OG/H-Y-109-N |
| DDR1 | 18.1% (2/11) | 4ag4 | | L-Y96-OH/H-Y96-OH |
| TSLP | 53.3% (8/15) | 5j13 | | L-Y89-OH/H-Q42-NE2 |
| CD38 | 0% (0/8) | 7duo | l_R91-NH1/H-E106-O | L-Q38-OE1/H-Q39-NE2 |
| | | | | L-Q38-NE2/H-Q39-OE1 |
| | | | | L-Q89-NE2/H-P107-O |
| CD47 | 0% (0/4) | 7xjf | L-N34-ND2/H-R108-O | L-Q38-OE1/H-Q39-NE2 |
| | | | L-G91-O/H-R108-NE | L-Q38-NE2/H-Q39-OE1 |
| | | | L-A92-O/H-R108-NH2 | L-Q89-NE2/H-G107-O |
| | | | L-M4-O/H-R44-NH1 | |
| | | | L-F98-O/H-R44-NH1 | |

| | | | | |
|---|---|---|---|---|
| Z13-IL22-2 | 20% (1/5) | 3q1s | | L-L33-N/H-Y111-OH |
| HAVR2 | 0% (0/1) | 7kql | | L-Q39-NE2/H-Y-96-OH |
| | | | | L-Y88-OH/H-Q41-NE2 |
| TNR9 | 50% (1/2) | 8gye | | |

Supplementary Table S7 — Interface chemistry features associated with success.
Target-level aggregation of structural contact features observed in successful designs, including source PDB references and counts of charged and other polar amino acids at the VH–VL and/or VL–antigen interfaces. These features contextualize observed success rates by interface polarity/charge composition.
Abbreviations: PDB, Protein Data Bank; AA, amino acid.

| Gene Type | Germline Gene | TheraSAbDab (%) | PDB Input (%) | Validated cLCs (%) | Parental Antibodies (%) |
|---|---|---|---|---|---|
| V | IGKV1-39 | 142 (13.37%) | 275 (11.36%) | 27 (31.76%) | 4 (30.77%) |
| V | IGKV3-11 | 69 (6.5%) | 97 (4.01%) | 6 (7.06%) | 2 (15.38%) |
| V | IGKV3-20 | 70 (6.59%) | 212 (8.76%) | 7 (8.24%) | 1 (7.69%) |
| V | IGKV1-12 | 44 (4.14%) | 61 (2.52%) | 5 (5.88%) | 1 (7.69%) |
| V | IGKV1-33 | 67 (6.31%) | 102 (4.21%) | 2 (2.35%) | 1 (7.69%) |
| V | IGLV3-21 | 21 (1.98%) | 70 (2.89%) | 2 (2.35%) | 1 (7.69%) |
| V | IGKV10-94 | 5 (0.47%) | 15 (0.62%) | | 1 (7.69%) |
| V | IGKV6-15 | 2 (0.19%) | 15 (0.62%) | | 1 (7.69%) |
| V | IGKV4-68 | 2 (0.19%) | 4 (0.17%) | | 1 (7.69%) |
| V | IGKV1-5 | 38 (3.58%) | 89 (3.68%) | 6 (7.06%) | |
| V | IGKV4-1 | 69 (6.5%) | 58 (2.4%) | 6 (7.06%) | |
| V | IGKV1-9 | 19 (1.79%) | 58 (2.4%) | 6 (7.06%) | |
| V | IGLV3-1 | 11 (1.04%) | 35 (1.45%) | 5 (5.88%) | |
| V | IGKV3-15 | 31 (2.92%) | 80 (3.31%) | 3 (3.53%) | |
| V | IGKV4-59 | 3 (0.28%) | 21 (0.87%) | 3 (3.53%) | |
| V | IGKV1-16 | 28 (2.64%) | 22 (0.91%) | 2 (2.35%) | |
| V | IGKV1S5 | | 1 (0.04%) | 5 (5.88%) | |
| J | IGKJ1 | 293 (27.59%) | 593 (24.5%) | 29 (34.12%) | 8 (61.54%) |
| J | IGKJ2 | 266 (25.05%) | 551 (22.77%) | 9 (10.59%) | 2 (15.38%) |
| J | IGKJ4 | 260 (24.48%) | 337 (13.93%) | 21 (24.71%) | 1 (7.69%) |
| J | IGKJ5 | 60 (5.65%) | 243 (10.04%) | 9 (10.59%) | 1 (7.69%) |
| J | IGLJ2 | 72 (6.78%) | 285 (11.78%) | 5 (5.88%) | 1 (7.69%) |
| J | IGKJ3 | 37 (3.48%) | 109 (4.5%) | 5 (5.88%) | |
| J | IGLJ3 | 53 (4.99%) | 183 (7.56%) | 2 (2.35%) | |
| J | IGKJ1-2 | | 1 (0.04%) | 5 (5.88%) | |

Supplementary Table S8 — Germline distribution and enrichment analysis.

Comparison of V-gene usage across reference sets and study outputs: TheraSAbDab (%), PDB input templates (%), validated cLCs (%), and parental antibodies (%). Enrichment factors are computed as (Validated cLCs %)/(Reference %) for each germline, stratified by gene type (κ/λ).
Abbreviations: PDB, Protein Data Bank; TheraSAbDab, Therapeutic Structural Antibody Database.

Design ID, mutations, benchmark_design_id, ag_uniprot, original_vl_type, original_vl_v_gene, original_vl_j_gene, original_vl_species, new_vl_type, new_vl_v_gene, new_vl_j_gene, new_vl_species
T1-A1, [], T1-Aa, VEGFA_HUMAN, K, IGKV1-12*01, IGKJ1*01, human, K, IGKV1-39*01, IGKJ1*01, human
T1-A2, [], T1-Aa, VEGFA_HUMAN, K, IGKV1-12*01, IGKJ1*01, human, K, IGKV1-12*01, IGKJ2*01, human
T1-A3, [], T1-Aa, VEGFA_HUMAN, K, IGKV1-12*01, IGKJ1*01, human, K, IGKV1-16*01, IGKJ1*01, human
T1-A4, [], T1-Aa, VEGFA_HUMAN, K, IGKV1-12*01, IGKJ1*01, human, K, IGKV1-39*01, IGKJ4*01, human
T1-G1, [], T1-Gg, ERBB2_HUMAN, K, IGKV1-39*01, IGKJ1*01, human, K, IGKV1-39*01, IGKJ1*01, human
T1-G2, [], T1-Gg, ERBB2_HUMAN, K, IGKV1-39*01, IGKJ1*01, human, K, IGKV1-39*01, IGKJ1*01, human
T1-G3, [], T1-Gg, ERBB2_HUMAN, K, IGKV1-39*01, IGKJ1*01, human, K, IGKV1-12*01, IGKJ2*01, human
T1-G4, [], T1-Gg, ERBB2_HUMAN, K, IGKV1-39*01, IGKJ1*01, human, K, IGKV1-39*01, IGKJ4*01, human
T1-C1, [], T1-Cc, CD19_HUMAN, K, IGKV1-39*01, IGKJ1*01, human, K, IGKV1-12*01, IGKJ2*01, human
T1-C2, [], T1-Cc, CD19_HUMAN, K, IGKV1-39*01, IGKJ1*01, human, K, IGKV1-12*01, IGKJ1*01, human
T1-C3, [], T1-Cc, CD19_HUMAN, K, IGKV1-39*01, IGKJ1*01, human, K, IGKV1S5*01, IGKJ1-2*01, rabbit
T1-C4, [], T1-Cc, CD19_HUMAN, K, IGKV1-39*01, IGKJ1*01, human, K, IGKV1-39*01, IGKJ1*01, human
T1-F1, [], T1-Ff, CD38_HUMAN, K, IGKV3-11*01, IGKJ1*01, human, K, IGKV3-11*01, IGKJ2*01, human
T1-E1, [], T1-Ee, CD47_HUMAN, K, IGKV1-39*01, IGKJ4*01, human, K, IGKV1-39*01, IGKJ1*01, human
T1-E2, [], T1-Ee, CD47_HUMAN, K, IGKV1-39*01, IGKJ4*01, human, K, IGKV1-39*01, IGKJ1*01, human
T1-E3, [], T1-Ee, CD47_HUMAN, K, IGKV1-39*01, IGKJ4*01, human, K, IGKV1-39*01, IGKJ4*01, human
T1-G5, [], T1-Gg, ERBB2_HUMAN, K, IGKV1-39*01, IGKJ1*01, human, K, IGKV4-59*01, IGKJ4*01, mouse
T1-G6, [], T1-Gg, ERBB2_HUMAN, K, IGKV1-39*01, IGKJ1*01, human, K, IGKV4-1*01, IGKJ4*01, human
T1-G7, [], T1-Gg, ERBB2_HUMAN, K, IGKV1-39*01, IGKJ1*01, human, K, IGKV4-1*01, IGKJ1*01, human
T1-F2, [], T1-Ff, CD38_HUMAN, K, IGKV3-11*01, IGKJ1*01, human, K, IGKV4-59*01, IGKJ4*01, mouse
T1-F3, [], T1-Ff, CD38_HUMAN, K, IGKV3-11*01, IGKJ1*01, human, K, IGKV4-1*01, IGKJ4*01, human
T1-F4, [], T1-Ff, CD38_HUMAN, K, IGKV3-11*01, IGKJ1*01, human, K, IGKV4-1*01, IGKJ1*01, human
T1-D1, [], T1-Dd, CD19_HUMAN, K, IGKV6-15*01, IGKJ2*01, mouse, K, IGKV4-59*01, IGKJ4*01, mouse
T1-D2, [], T1-Dd, CD19_HUMAN, K, IGKV6-15*01, IGKJ2*01, mouse, K, IGKV4-1*01, IGKJ4*01, human
T1-D3, [], T1-Dd, CD19_HUMAN, K, IGKV6-15*01, IGKJ2*01, mouse, K, IGKV4-1*01, IGKJ1*01, human
T1-G8, [], T1-Gg, ERBB2_HUMAN, K, IGKV1-39*01, IGKJ1*01, human, K, IGKV3-15*01, IGKJ4*01, human
T1-F5, [], T1-Ff, CD38_HUMAN, K, IGKV3-11*01, IGKJ1*01, human, K, IGKV3-15*01, IGKJ4*01, human
T1-F6, [], T1-Ff, CD38_HUMAN, K, IGKV3-11*01, IGKJ1*01, human, K, IGKV1-39*01, IGKJ1*01, human
T1-D4, [], T1-Dd, CD19_HUMAN, K, IGKV6-15*01, IGKJ2*01, mouse, K, IGKV3-15*01, IGKJ4*01, human
T1-D5, [], T1-Dd, CD19_HUMAN, K, IGKV6-15*01, IGKJ2*01, mouse, K, IGKV1-39*01, IGKJ1*01, human
T1-A5, [], T1-Aa, VEGFA_HUMAN, K, IGKV1-12*01, IGKJ1*01, human, K, IGKV1S5*01, IGKJ1-2*01, rabbit
T1-G9, [], T1-Gg, ERBB2_HUMAN, K, IGKV1-39*01, IGKJ1*01, human, K, IGKV1-39*01, IGKJ1*01, human
T1-G10, [], T1-Gg, ERBB2_HUMAN, K, IGKV1-39*01, IGKJ1*01, human, K, IGKV1S5*01, IGKJ1-2*01, rabbit
T1-B1, [], T1-Bb, VEGFA_HUMAN, K, IGKV1-33*01, IGKJ1*01, human, K, IGKV1-39*01, IGKJ1*01, human
T1-B2, [], T1-Bb, VEGFA_HUMAN, K, IGKV1-33*01, IGKJ1*01, human, K, IGKV1-39*01, IGKJ1*01, human
T1-B3, [], T1-Bb, VEGFA_HUMAN, K, IGKV1-33*01, IGKJ1*01, human, K, IGKV1-12*01, IGKJ2*01, human
T1-B4, [], T1-Bb, VEGFA_HUMAN, K, IGKV1-33*01, IGKJ1*01, human, K, IGKV1-16*01, IGKJ1*01, human
T1-B5, [], T1-Bb, VEGFA_HUMAN, K, IGKV1-33*01, IGKJ1*01, human, K, IGKV1-5*03, IGKJ1*01, human
T1-B6, [], T1-Bb, VEGFA_HUMAN, K, IGKV1-33*01, IGKJ1*01, human, K, IGKV1-39*01, IGKJ1*01, human
T1-B7, [], T1-Bb, VEGFA_HUMAN, K, IGKV1-33*01, IGKJ1*01, human, K, IGKV1-39*01, IGKJ1*01, human
T1-B8, [], T1-Bb, VEGFA_HUMAN, K, IGKV1-33*01, IGKJ1*01, human, K, IGKV1S5*01, IGKJ1-2*01, rabbit
T1-B9, [], T1-Bb, VEGFA_HUMAN, K, IGKV1-33*01, IGKJ1*01, human, K, IGKV1-39*01, IGKJ1*01, human
T1-B10, [], T1-Bb, VEGFA_HUMAN, K, IGKV1-33*01, IGKJ1*01, human, K, IGKV1-39*01, IGKJ4*01, human
T1-B11, [], T1-Bb, VEGFA_HUMAN, K, IGKV1-33*01, IGKJ1*01, human, K, IGKV1-39*01, IGKJ4*01, human
T1-B12, [], T1-Bb, VEGFA_HUMAN, K, IGKV1-33*01, IGKJ1*01, human, K, IGKV1-5*03, IGKJ1*01, human
T1-B13, [], T1-Bb, VEGFA_HUMAN, K, IGKV1-33*01, IGKJ1*01, human, K, IGKV1-39*01, IGKJ5*01, human
T1-F7, [], T1-Ff, CD38_HUMAN, K, IGKV3-11*01, IGKJ1*01, human, K, IGKV3-11*01, IGKJ4*01, human
T1-F8, [], T1-Ff, CD38_HUMAN, K, IGKV3-11*01, IGKJ1*01, human, K, IGKV3-20*01, IGKJ3*01, human
T1-E4, [], T1-Ee, CD47_HUMAN, K, IGKV1-39*01, IGKJ4*01, human, K, IGKV1S5*01, IGKJ1-2*01, rabbit
T1-Aa, [], T1-Aa, VEGFA_HUMAN, K, IGKV1-12*01, IGKJ1*01, human, K, IGKV1-12*01, IGKJ1*01, human

T1-Cc, [], T1-Cc, CD19_HUMAN, K, IGKV1-39*01, IGKJ1*01, human, K, IGKV1-39*01, IGKJ1*01, human
T1-Bb, [], T1-Bb, VEGFA_HUMAN, K, IGKV1-33*01, IGKJ1*01, human, K, IGKV1-33*01, IGKJ1*01, human
T1-Dd, [], T1-Dd, CD19_HUMAN, K, IGKV6-15*01, IGKJ2*01, mouse, K, IGKV6-15*01, IGKJ2*01, mouse
T1-Ee, [], T1-Ee, CD47_HUMAN, K, IGKV1-39*01, IGKJ4*01, human, K, IGKV1-39*01, IGKJ4*01, human
T1-Ff, [], T1-Ff, CD38_HUMAN, K, IGKV3-11*01, IGKJ1*01, human, K, IGKV3-11*01, IGKJ1*01, human
T1-Gg, [], T1-Gg, ERBB2_HUMAN, K, IGKV1-39*01, IGKJ1*01, human, K, IGKV1-39*01, IGKJ1*01, human
T2-K1, [], T2-Kk, DDR1_HUMAN, K, IGKV4-68*01, IGKJ2*01, mouse, L, IGLV3-21*02, IGLJ2*01, human
T2-M1, [], T2-Mm, TSLP_HUMAN, L, IGLV3-21*02, IGLJ2*01, human, L, IGLV3-21*02, IGLJ2*01, human
T2-I1, [], T2-Ii, IL22_HUMAN, K, IGKV3-11*01, IGKJ1*01, human, L, IGLV3-1*01, IGLJ3*02, human
T2-M2, [], T2-Mm, TSLP_HUMAN, L, IGLV3-21*02, IGLJ2*01, human, L, IGLV3-1*01, IGLJ3*02, human
T2-K2, [], T2-Kk, DDR1_HUMAN, K, IGKV4-68*01, IGKJ2*01, mouse, K, IGKV3-11*01, IGKJ2*01, human
T2-M3, [], T2-Mm, TSLP_HUMAN, L, IGLV3-21*02, IGLJ2*01, human, K, IGKV3-11*01, IGKJ2*01, human
T2-K3, [], T2-Kk, DDR1_HUMAN, K, IGKV4-68*01, IGKJ2*01, mouse, K, IGKV3-11*01, IGKJ3*01, human
T2-M4, [], T2-Mm, TSLP_HUMAN, L, IGLV3-21*02, IGLJ2*01, human, K, IGKV3-11*01, IGKJ3*01, human
T2-M5, [], T2-Mm, TSLP_HUMAN, L, IGLV3-21*02, IGLJ2*01, human, K, IGKV3-20*01, IGKJ4*01, human
T2-L1, [], T2-Ll, ERBB2_HUMAN, K, IGKV1-39*01, IGKJ1*01, human, K, IGKV3-20*01, IGKJ4*01, human
T2-K4, [], T2-Kk, DDR1_HUMAN, K, IGKV4-68*01, IGKJ2*01, mouse, K, IGKV1-9*01, IGKJ3*01, human
T2-M6, [], T2-Mm, TSLP_HUMAN, L, IGLV3-21*02, IGLJ2*01, human, K, IGKV1-9*01, IGKJ3*01, human
T2-K5, [], T2-Kk, DDR1_HUMAN, K, IGKV4-68*01, IGKJ2*01, mouse, K, IGKV1-39*01, IGKJ4*01, human
T2-M7, [], T2-Mm, TSLP_HUMAN, L, IGLV3-21*02, IGLJ2*01, human, K, IGKV1-39*01, IGKJ4*01, human
T2-K6, [], T2-Kk, DDR1_HUMAN, K, IGKV4-68*01, IGKJ2*01, mouse, K, IGKV1-9*03, IGKJ5*01, human
T2-M8, [], T2-Mm, TSLP_HUMAN, L, IGLV3-21*02, IGLJ2*01, human, K, IGKV1-9*03, IGKJ5*01, human
T2-J1, [], T2-Jj, HAVR2_HUMAN, K, IGKV3-20*01, IGKJ5*01, human, K, IGKV1-9*03, IGKJ5*01, human
T2-H1, [], T2-Hh, TNR9_HUMAN, K, IGKV10-94*01, IGKJ1*01, mouse, K, IGKV1-9*03, IGKJ5*01, human
T2-K7, [], T2-Kk, DDR1_HUMAN, K, IGKV4-68*01, IGKJ2*01, mouse, K, IGKV1-33*01, IGKJ2*01, human
T2-M9, [], T2-Mm, TSLP_HUMAN, L, IGLV3-21*02, IGLJ2*01, human, K, IGKV1-33*01, IGKJ2*01, human
T2-I2, [], T2-Ii, IL22_HUMAN, K, IGKV3-11*01, IGKJ1*01, human, K, IGKV3-20*01, IGKJ4*01, human
T2-M10, [], T2-Mm, TSLP_HUMAN, L, IGLV3-21*02, IGLJ2*01, human, K, IGKV3-20*01, IGKJ4*01, human
T2-K8, [], T2-Kk, DDR1_HUMAN, K, IGKV4-68*01, IGKJ2*01, mouse, K, IGKV1-5*01, IGKJ5*01, human
T2-M11, [], T2-Mm, TSLP_HUMAN, L, IGLV3-21*02, IGLJ2*01, human, K, IGKV1-5*01, IGKJ5*01, human
T2-I3, [], T2-Ii, IL22_HUMAN, K, IGKV3-11*01, IGKJ1*01, human, K, IGKV3-20*01, IGKJ1*01, human
T2-M12, [], T2-Mm, TSLP_HUMAN, L, IGLV3-21*02, IGLJ2*01, human, K, IGKV3-20*01, IGKJ1*01, human
T2-I4, [], T2-Ii, IL22_HUMAN, K, IGKV3-11*01, IGKJ1*01, human, L, IGLV3-1*01, IGLJ2*01, human
T2-K9, [], T2-Kk, DDR1_HUMAN, K, IGKV4-68*01, IGKJ2*01, mouse, L, IGLV3-1*01, IGLJ2*01, human
T2-L2, [], T2-Ll, ERBB2_HUMAN, K, IGKV1-39*01, IGKJ1*01, human, L, IGLV3-1*01, IGLJ2*01, human
T2-K10, [], T2-Kk, DDR1_HUMAN, K, IGKV4-68*01, IGKJ2*01, mouse, K, IGKV1-5*01, IGKJ1*01, human
T2-M13, [], T2-Mm, TSLP_HUMAN, L, IGLV3-21*02, IGLJ2*01, human, K, IGKV1-5*01, IGKJ1*01, human
T2-I5, [], T2-Ii, IL22_HUMAN, K, IGKV3-11*01, IGKJ1*01, human, K, IGKV1-39*01, IGKJ1*01, human
T2-M14, [], T2-Mm, TSLP_HUMAN, L, IGLV3-21*02, IGLJ2*01, human, K, IGKV1-39*01, IGKJ1*01, human
T2-H2, [], T2-Hh, TNR9_HUMAN, K, IGKV10-94*01, IGKJ1*01, mouse, K, IGKV1-39*01, IGKJ1*01, human
T2-K11, [], T2-Kk, DDR1_HUMAN, K, IGKV4-68*01, IGKJ2*01, mouse, K, IGKV1-39*01, IGKJ5*01, human
T2-M15, [], T2-Mm, TSLP_HUMAN, L, IGLV3-21*02, IGLJ2*01, human, K, IGKV1-39*01, IGKJ5*01, human
T2-Ii, [], T2-Ii, IL22_HUMAN, K, IGKV3-11*01, IGKJ1*01, human, K, IGKV3-11*01, IGKJ1*01, human
T2-Kk, [], T2-Kk, DDR1_HUMAN, K, IGKV4-68*01, IGKJ2*01, mouse, K, IGKV4-68*01, IGKJ2*01, mouse
T2-Mm, [], T2-Mm, TSLP_HUMAN, L, IGLV3-21*02, IGLJ2*01, human, L, IGLV3-21*02, IGLJ2*01, human
T2-Ll, [], T2-Ll, ERBB2_HUMAN, K, IGKV1-39*01, IGKJ1*01, human, K, IGKV1-39*01, IGKJ1*01, human
T2-Jj, [], T2-Jj, HAVR2_HUMAN, K, IGKV3-20*01, IGKJ5*01, human, K, IGKV3-20*01, IGKJ5*01, human
T2-Hh, [], T2-Hh, TNR9_HUMAN, K, IGKV10-94*01, IGKJ1*01, mouse, K, IGKV10-94*01, IGKJ1*01, mouse
T2-K1, ['E49L_L', 'E52N_L'], T2-Kk, DDR1_HUMAN, K, IGKV4-68*01, IGKJ2*01, mouse, L, IGLV3-21*01, IGLJ2*01, human
T2-M1, ['E49L_L', 'E52N_L'], T2-Mm, TSLP_HUMAN, L, IGLV3-21*02, IGLJ2*01, human, L, IGLV3-21*01, IGLJ2*01, human
T2-K1, ['E49L_L', 'E52N_L', 'Y31F_L'], T2-Kk, DDR1_HUMAN, K, IGKV4-68*01, IGKJ2*01, mouse, L, IGLV3-21*01, IGLJ2*01, human
T2-M1, ['E49L_L', 'E52N_L', 'Y31F_L'], T2-Mm, TSLP_HUMAN, L, IGLV3-21*02, IGLJ2*01, human, L, IGLV3-21*01, IGLJ2*01, human

T2-K1, ['E49L_L', 'Y31F_L'], T2-Kk, DDR1_HUMAN, K, IGKV4-68*01, IGKJ2*01, mouse, L, IGLV3-21*01, IGLJ2*01, human
T2-M1, ['E49L_L', 'Y31F_L'], T2-Mm, TSLP_HUMAN, L, IGLV3-21*02, IGLJ2*01, human, L, IGLV3-21*01, IGLJ2*01, human
T2-K1, ['E52N_L'], T2-Kk, DDR1_HUMAN, K, IGKV4-68*01, IGKJ2*01, mouse, L, IGLV3-21*02, IGLJ2*01, human
T2-M1, ['E52N_L'], T2-Mm, TSLP_HUMAN, L, IGLV3-21*02, IGLJ2*01, human, L, IGLV3-21*02, IGLJ2*01, human
T2-K1, ['E52N_L', 'Y31F_L'], T2-Kk, DDR1_HUMAN, K, IGKV4-68*01, IGKJ2*01, mouse, L, IGLV3-21*02, IGLJ2*01, human
T2-M1, ['E52N_L', 'Y31F_L'], T2-Mm, TSLP_HUMAN, L, IGLV3-21*02, IGLJ2*01, human, L, IGLV3-21*02, IGLJ2*01, human
T2-K1, ['Y31F_L'], T2-Kk, DDR1_HUMAN, K, IGKV4-68*01, IGKJ2*01, mouse, L, IGLV3-21*02, IGLJ2*01, human
T2-M1, ['Y31F_L'], T2-Mm, TSLP_HUMAN, L, IGLV3-21*02, IGLJ2*01, human, L, IGLV3-21*02, IGLJ2*01, human
T2-K2, ['T53N_L'], T2-Kk, DDR1_HUMAN, K, IGKV4-68*01, IGKJ2*01, mouse, K, IGKV3-11*01, IGKJ2*01, human
T2-M3, ['T53N_L'], T2-Mm, TSLP_HUMAN, L, IGLV3-21*02, IGLJ2*01, human, K, IGKV3-11*01, IGKJ2*01, human
T2-K3, ['W94N_L'], T2-Kk, DDR1_HUMAN, K, IGKV4-68*01, IGKJ2*01, mouse, K, IGKV3-11*01, IGKJ3*01, human
T2-M4, ['W94N_L'], T2-Mm, TSLP_HUMAN, L, IGLV3-21*02, IGLJ2*01, human, K, IGKV3-11*01, IGKJ3*01, human
T2-K3, ['Y30T_L', 'W94N_L'], T2-Kk, DDR1_HUMAN, K, IGKV4-68*01, IGKJ2*01, mouse, K, IGKV3-11*01, IGKJ3*01, human
T2-M4, ['Y30T_L', 'W94N_L'], T2-Mm, TSLP_HUMAN, L, IGLV3-21*02, IGLJ2*01, human, K, IGKV3-11*01, IGKJ3*01, human
T2-K3, ['Y32F_L'], T2-Kk, DDR1_HUMAN, K, IGKV4-68*01, IGKJ2*01, mouse, K, IGKV3-11*01, IGKJ3*01, human
T2-M4, ['Y32F_L'], T2-Mm, TSLP_HUMAN, L, IGLV3-21*02, IGLJ2*01, human, K, IGKV3-11*01, IGKJ3*01, human
T2-K3, ['Y32F_L', 'W94N_L'], T2-Kk, DDR1_HUMAN, K, IGKV4-68*01, IGKJ2*01, mouse, K, IGKV3-11*01, IGKJ3*01, human
T2-M4, ['Y32F_L', 'W94N_L'], T2-Mm, TSLP_HUMAN, L, IGLV3-21*02, IGLJ2*01, human, K, IGKV3-11*01, IGKJ3*01, human
T2-K3, ['Y32F_L', 'Y30T_L', 'W94N_L'], T2-Kk, DDR1_HUMAN, K, IGKV4-68*01, IGKJ2*01, mouse, K, IGKV3-11*01, IGKJ3*01, human
T2-M4, ['Y32F_L', 'Y30T_L', 'W94N_L'], T2-Mm, TSLP_HUMAN, L, IGLV3-21*02, IGLJ2*01, human, K, IGKV3-11*01, IGKJ3*01, human
T2-K5, ['A50D_L'], T2-Kk, DDR1_HUMAN, K, IGKV4-68*01, IGKJ2*01, mouse, K, IGKV1-39*01, IGKJ4*01, human
T2-M7, ['A50D_L'], T2-Mm, TSLP_HUMAN, L, IGLV3-21*02, IGLJ2*01, human, K, IGKV1-39*01, IGKJ4*01, human
T2-K5, ['A50D_L', 'L95H_L'], T2-Kk, DDR1_HUMAN, K, IGKV4-68*01, IGKJ2*01, mouse, K, IGKV1-39*01, IGKJ4*01, human
T2-M7, ['A50D_L', 'L95H_L'], T2-Mm, TSLP_HUMAN, L, IGLV3-21*02, IGLJ2*01, human, K, IGKV1-39*01, IGKJ4*01, human
T2-K5, ['L95H_L'], T2-Kk, DDR1_HUMAN, K, IGKV4-68*01, IGKJ2*01, mouse, K, IGKV1-39*01, IGKJ4*01, human
T2-M7, ['L95H_L'], T2-Mm, TSLP_HUMAN, L, IGLV3-21*02, IGLJ2*01, human, K, IGKV1-39*01, IGKJ4*01, human
T2-K5, ['S53N_L'], T2-Kk, DDR1_HUMAN, K, IGKV4-68*01, IGKJ2*01, mouse, K, IGKV1-39*01, IGKJ4*01, human
T2-M7, ['S53N_L'], T2-Mm, TSLP_HUMAN, L, IGLV3-21*02, IGLJ2*01, human, K, IGKV1-39*01, IGKJ4*01, human
T2-K7, ['D92S_L'], T2-Kk, DDR1_HUMAN, K, IGKV4-68*01, IGKJ2*01, mouse, K, IGKV1-33*01, IGKJ2*01, human
T2-M9, ['D92S_L'], T2-Mm, TSLP_HUMAN, L, IGLV3-21*02, IGLJ2*01, human, K, IGKV1-33*01, IGKJ2*01, human
T2-K7, ['D92S_L', 'D93S_L'], T2-Kk, DDR1_HUMAN, K, IGKV4-68*01, IGKJ2*01, mouse, K, IGKV1-33*01, IGKJ2*01, human
T2-M9, ['D92S_L', 'D93S_L'], T2-Mm, TSLP_HUMAN, L, IGLV3-21*02, IGLJ2*01, human, K, IGKV1-33*01, IGKJ2*01, human
T2-K7, ['D93S_L'], T2-Kk, DDR1_HUMAN, K, IGKV4-68*01, IGKJ2*01, mouse, K, IGKV1-33*01, IGKJ2*01, human
T2-M9, ['D93S_L'], T2-Mm, TSLP_HUMAN, L, IGLV3-21*02, IGLJ2*01, human, K, IGKV1-33*01, IGKJ2*01, human
T2-I2, ['R97H_L'], T2-Ii, IL22_HUMAN, K, IGKV3-11*01, IGKJ1*01, human, K, IGKV3-20*01, IGKJ4*01, human
T2-M10, ['R97H_L'], T2-Mm, TSLP_HUMAN, L, IGLV3-21*02, IGLJ2*01, human, K, IGKV3-20*01, IGKJ4*01, human
T2-K8, ['Y94S_L'], T2-Kk, DDR1_HUMAN, K, IGKV4-68*01, IGKJ2*01, mouse, K, IGKV1-5*01, IGKJ5*01, human
T2-M11, ['Y94S_L'], T2-Mm, TSLP_HUMAN, L, IGLV3-21*02, IGLJ2*01, human, K, IGKV1-5*01, IGKJ5*01, human
T2-I4, ['C33A_L'], T2-Ii, IL22_HUMAN, K, IGKV3-11*01, IGKJ1*01, human, L, IGLV3-1*01, IGLJ2*01, human
T2-K9, ['C33A_L'], T2-Kk, DDR1_HUMAN, K, IGKV4-68*01, IGKJ2*01, mouse, L, IGLV3-1*01, IGLJ2*01, human
T2-L2, ['C33A_L'], T2-Ll, ERBB2_HUMAN, K, IGKV1-39*01, IGKJ1*01, human, L, IGLV3-1*01, IGLJ2*01, human
T2-I4, ['C33S_L'], T2-Ii, IL22_HUMAN, K, IGKV3-11*01, IGKJ1*01, human, L, IGLV3-1*01, IGLJ2*01, human
T2-K9, ['C33S_L'], T2-Kk, DDR1_HUMAN, K, IGKV4-68*01, IGKJ2*01, mouse, L, IGLV3-1*01, IGLJ2*01, human

T2-L2, ['C33S_L'], T2-Ll, ERBB2_HUMAN, K, IGKV1-39*01, IGKJ1*01, human, L, IGLV3-1*01, IGLJ2*01, human
T2-I4, ['Y31F_L'], T2-Ii, IL22_HUMAN, K, IGKV3-11*01, IGKJ1*01, human, L, IGLV3-1*01, IGLJ2*01, human
T2-L2, ['Y31F_L'], T2-Ll, ERBB2_HUMAN, K, IGKV1-39*01, IGKJ1*01, human, L, IGLV3-1*01, IGLJ2*01, human
T2-I4, ['Y31F_L', 'C33A_L'], T2-Ii, IL22_HUMAN, K, IGKV3-11*01, IGKJ1*01, human, L, IGLV3-1*01, IGLJ2*01, human
T2-L2, ['Y31F_L', 'C33A_L'], T2-Ll, ERBB2_HUMAN, K, IGKV1-39*01, IGKJ1*01, human, L, IGLV3-1*01, IGLJ2*01, human
T2-I4, ['Y31F_L', 'C33S_L'], T2-Ii, IL22_HUMAN, K, IGKV3-11*01, IGKJ1*01, human, L, IGLV3-1*01, IGLJ2*01, human
T2-L2, ['Y31F_L', 'C33S_L'], T2-Ll, ERBB2_HUMAN, K, IGKV1-39*01, IGKJ1*01, human, L, IGLV3-1*01, IGLJ2*01, human
T2-K10, ['Y94S_L'], T2-Kk, DDR1_HUMAN, K, IGKV4-68*01, IGKJ2*01, mouse, K, IGKV1-5*01, IGKJ1*01, human
T2-M13, ['Y94S_L'], T2-Mm, TSLP_HUMAN, L, IGLV3-21*02, IGLJ2*01, human, K, IGKV1-5*01, IGKJ1*01, human
T2-M14, ['S50D_L'], T2-Mm, TSLP_HUMAN, L, IGLV3-21*02, IGLJ2*01, human, K, IGKV1-39*01, IGKJ1*01, human
T2-H2, ['S50D_L'], T2-Hh, TNR9_HUMAN, K, IGKV10-94*01, IGKJ1*01, mouse, K, IGKV1-39*01, IGKJ1*01, human
T2-I5, ['W92S_L'], T2-Ii, IL22_HUMAN, K, IGKV3-11*01, IGKJ1*01, human, K, IGKV1-39*01, IGKJ1*01, human
T2-M14, ['W92S_L'], T2-Mm, TSLP_HUMAN, L, IGLV3-21*02, IGLJ2*01, human, K, IGKV1-39*01, IGKJ1*01, human
T2-H2, ['W92S_L'], T2-Hh, TNR9_HUMAN, K, IGKV10-94*01, IGKJ1*01, mouse, K, IGKV1-39*01, IGKJ1*01, human
T2-K11, ['L96H_L'], T2-Kk, DDR1_HUMAN, K, IGKV4-68*01, IGKJ2*01, mouse, K, IGKV1-39*01, IGKJ5*01, human
T2-M15, ['L96H_L'], T2-Mm, TSLP_HUMAN, L, IGLV3-21*02, IGLJ2*01, human, K, IGKV1-39*01, IGKJ5*01, human
T2-K11, ['W93S_L'], T2-Kk, DDR1_HUMAN, K, IGKV4-68*01, IGKJ2*01, mouse, K, IGKV1-39*01, IGKJ5*01, human
T2-M15, ['W93S_L'], T2-Mm, TSLP_HUMAN, L, IGLV3-21*02, IGLJ2*01, human, K, IGKV1-39*01, IGKJ5*01, human
T2-M14, ['S50D_L', 'A32Y_L'], T2-Mm, TSLP_HUMAN, L, IGLV3-21*02, IGLJ2*01, human, K, IGKV1-39*01, IGKJ1*01, human
T2-H2, ['S50D_L', 'A32Y_L'], T2-Hh, TNR9_HUMAN, K, IGKV10-94*01, IGKJ1*01, mouse, K, IGKV1-39*01, IGKJ1*01, human
T2-M14, ['S50D_L', 'A32Y_L', 'W92S_L'], T2-Mm, TSLP_HUMAN, L, IGLV3-21*02, IGLJ2*01, human, K, IGKV1-39*01, IGKJ1*01, human
T2-H2, ['S50D_L', 'A32Y_L', 'W92S_L'], T2-Hh, TNR9_HUMAN, K, IGKV10-94*01, IGKJ1*01, mouse, K, IGKV1-39*01, IGKJ1*01, human
T2-M14, ['S50D_L', 'W92S_L'], T2-Mm, TSLP_HUMAN, L, IGLV3-21*02, IGLJ2*01, human, K, IGKV1-13*02, IGKJ1*01, human
T2-H2, ['S50D_L', 'W92S_L'], T2-Hh, TNR9_HUMAN, K, IGKV10-94*01, IGKJ1*01, mouse, K, IGKV1-13*02, IGKJ1*01, human
T2-K11, ['W93S_L', 'L96H_L'], T2-Kk, DDR1_HUMAN, K, IGKV4-68*01, IGKJ2*01, mouse, K, IGKV1-39*01, IGKJ5*01, human
T2-M15, ['W93S_L', 'L96H_L'], T2-Mm, TSLP_HUMAN, L, IGLV3-21*02, IGLJ2*01, human, K, IGKV1-39*01, IGKJ5*01, human
T2-K1, ['F95H_L'], T2-Kk, DDR1_HUMAN, K, IGKV4-68*01, IGKJ2*01, mouse, L, IGLV3-21*02, IGLJ2*01, human
T2-M1, ['F95H_L'], T2-Mm, TSLP_HUMAN, L, IGLV3-21*02, IGLJ2*01, human, L, IGLV3-21*02, IGLJ2*01, human
T2-K1, ['S55A_L'], T2-Kk, DDR1_HUMAN, K, IGKV4-68*01, IGKJ2*01, mouse, L, IGLV3-21*02, IGLJ2*01, human
T2-M1, ['S55A_L'], T2-Mm, TSLP_HUMAN, L, IGLV3-21*02, IGLJ2*01, human, L, IGLV3-21*02, IGLJ2*01, human
T2-K1, ['S55E_L'], T2-Kk, DDR1_HUMAN, K, IGKV4-68*01, IGKJ2*01, mouse, L, IGLV3-21*02, IGLJ2*01, human
T2-M1, ['S55E_L'], T2-Mm, TSLP_HUMAN, L, IGLV3-21*02, IGLJ2*01, human, L, IGLV3-21*02, IGLJ2*01, human
T2-K1, ['S55E_L', 'F95H_L'], T2-Kk, DDR1_HUMAN, K, IGKV4-68*01, IGKJ2*01, mouse, L, IGLV3-21*02, IGLJ2*01, human
T2-M1, ['S55E_L', 'F95H_L'], T2-Mm, TSLP_HUMAN, L, IGLV3-21*02, IGLJ2*01, human, L, IGLV3-21*02, IGLJ2*01, human
T2-K1, ['S55E_L', 'Y30K_L'], T2-Kk, DDR1_HUMAN, K, IGKV4-68*01, IGKJ2*01, mouse, L, IGLV3-21*02, IGLJ2*01, human
T2-M1, ['S55E_L', 'Y30K_L'], T2-Mm, TSLP_HUMAN, L, IGLV3-21*02, IGLJ2*01, human, L, IGLV3-21*02, IGLJ2*01, human
T2-K1, ['S55T_L'], T2-Kk, DDR1_HUMAN, K, IGKV4-68*01, IGKJ2*01, mouse, L, IGLV3-21*02, IGLJ2*01, human
T2-M1, ['S55T_L'], T2-Mm, TSLP_HUMAN, L, IGLV3-21*02, IGLJ2*01, human, L, IGLV3-21*02, IGLJ2*01, human
T2-M5, ['D93S_L', 'R94T_L', 'L97T_L', 'T98F_L'], T2-Mm, TSLP_HUMAN, L, IGLV3-21*02, IGLJ2*01, human, K, IGKV3-20*01, IGKJ4*01, human

T2-L1, ['D93S_L', 'R94T_L', 'L97T_L', 'T98F_L'], T2-Ll, ERBB2_HUMAN, K, IGKV1-39*01, IGKJ1*01, human, K, IGKV3-20*01, IGKJ4*01, human
T2-M5, ['L97del_L'], T2-Mm, TSLP_HUMAN, L, IGLV3-21*02, IGLJ2*01, human, K, IGKV3-20*01, IGKJ4*01, human
T2-L1, ['L97del_L'], T2-Ll, ERBB2_HUMAN, K, IGKV1-39*01, IGKJ1*01, human, K, IGKV3-20*01, IGKJ4*01, human
T2-M5, ['R94T_L', 'Y30S_L', 'D93S_L', 'P95S_L'], T2-Mm, TSLP_HUMAN, L, IGLV3-21*02, IGLJ2*01, human, K, IGKV3-20*01, IGKJ4*01, human
T2-L1, ['R94T_L', 'Y30S_L', 'D93S_L', 'P95S_L'], T2-Ll, ERBB2_HUMAN, K, IGKV1-39*01, IGKJ1*01, human, K, IGKV3-20*01, IGKJ4*01, human
T2-M5, ['R94T_L', 'Y30S_L', 'P95S_L'], T2-Mm, TSLP_HUMAN, L, IGLV3-21*02, IGLJ2*01, human, K, IGKV3-20*01, IGKJ4*01, human
T2-L1, ['R94T_L', 'Y30S_L', 'P95S_L'], T2-Ll, ERBB2_HUMAN, K, IGKV1-39*01, IGKJ1*01, human, K, IGKV3-20*01, IGKJ4*01, human
T2-M5, ['R94T_L', 'Y30S_L', 'Y92S_L', 'P95S_L'], T2-Mm, TSLP_HUMAN, L, IGLV3-21*02, IGLJ2*01, human, K, IGKV3-20*01, IGKJ4*01, human
T2-L1, ['R94T_L', 'Y30S_L', 'Y92S_L', 'P95S_L'], T2-Ll, ERBB2_HUMAN, K, IGKV1-39*01, IGKJ1*01, human, K, IGKV3-20*01, IGKJ4*01, human
T2-M5, ['Y30S_L'], T2-Mm, TSLP_HUMAN, L, IGLV3-21*02, IGLJ2*01, human, K, IGKV3-20*01, IGKJ4*01, human
T2-L1, ['Y30S_L'], T2-Ll, ERBB2_HUMAN, K, IGKV1-39*01, IGKJ1*01, human, K, IGKV3-20*01, IGKJ4*01, human
T2-M5, ['Y30S_L', 'D93S_L'], T2-Mm, TSLP_HUMAN, L, IGLV3-21*02, IGLJ2*01, human, K, IGKV3-20*01, IGKJ4*01, human
T2-L1, ['Y30S_L', 'D93S_L'], T2-Ll, ERBB2_HUMAN, K, IGKV1-39*01, IGKJ1*01, human, K, IGKV3-20*01, IGKJ4*01, human
T2-M5, ['Y30S_L', 'D93S_L', 'L97T_L'], T2-Mm, TSLP_HUMAN, L, IGLV3-21*02, IGLJ2*01, human, K, IGKV3-20*01, IGKJ4*01, human
T2-L1, ['Y30S_L', 'D93S_L', 'L97T_L'], T2-Ll, ERBB2_HUMAN, K, IGKV1-39*01, IGKJ1*01, human, K, IGKV3-20*01, IGKJ4*01, human
T2-M5, ['Y30S_L', 'D93S_L', 'P95S_L'], T2-Mm, TSLP_HUMAN, L, IGLV3-21*02, IGLJ2*01, human, K, IGKV3-20*01, IGKJ4*01, human
T2-L1, ['Y30S_L', 'D93S_L', 'P95S_L'], T2-Ll, ERBB2_HUMAN, K, IGKV1-39*01, IGKJ1*01, human, K, IGKV3-20*01, IGKJ4*01, human
T2-M5, ['Y30S_L', 'D93S_L', 'Y92S_L'], T2-Mm, TSLP_HUMAN, L, IGLV3-21*02, IGLJ2*01, human, K, IGKV3-20*01, IGKJ4*01, human
T2-L1, ['Y30S_L', 'D93S_L', 'Y92S_L'], T2-Ll, ERBB2_HUMAN, K, IGKV1-39*01, IGKJ1*01, human, K, IGKV3-20*01, IGKJ4*01, human
T2-M5, ['Y30S_L', 'D93S_L', 'Y92S_L', 'P95S_L'], T2-Mm, TSLP_HUMAN, L, IGLV3-21*02, IGLJ2*01, human, K, IGKV3-20*01, IGKJ4*01, human
T2-L1, ['Y30S_L', 'D93S_L', 'Y92S_L', 'P95S_L'], T2-Ll, ERBB2_HUMAN, K, IGKV1-39*01, IGKJ1*01, human, K, IGKV3-20*01, IGKJ4*01, human
T2-M5, ['Y30S_L', 'L97T_L'], T2-Mm, TSLP_HUMAN, L, IGLV3-21*02, IGLJ2*01, human, K, IGKV3-20*01, IGKJ4*01, human
T2-L1, ['Y30S_L', 'L97T_L'], T2-Ll, ERBB2_HUMAN, K, IGKV1-39*01, IGKJ1*01, human, K, IGKV3-20*01, IGKJ4*01, human
T2-M5, ['Y30S_L', 'L97T_L', 'Y92S_L'], T2-Mm, TSLP_HUMAN, L, IGLV3-21*02, IGLJ2*01, human, K, IGKV3-20*01, IGKJ4*01, human
T2-L1, ['Y30S_L', 'L97T_L', 'Y92S_L'], T2-Ll, ERBB2_HUMAN, K, IGKV1-39*01, IGKJ1*01, human, K, IGKV3-20*01, IGKJ4*01, human
T2-M5, ['Y30S_L', 'P95S_L'], T2-Mm, TSLP_HUMAN, L, IGLV3-21*02, IGLJ2*01, human, K, IGKV3-20*01, IGKJ4*01, human
T2-L1, ['Y30S_L', 'P95S_L'], T2-Ll, ERBB2_HUMAN, K, IGKV1-39*01, IGKJ1*01, human, K, IGKV3-20*01, IGKJ4*01, human
T2-M5, ['Y30S_L', 'Y92S_L'], T2-Mm, TSLP_HUMAN, L, IGLV3-21*02, IGLJ2*01, human, K, IGKV3-20*01, IGKJ4*01, human
T2-L1, ['Y30S_L', 'Y92S_L'], T2-Ll, ERBB2_HUMAN, K, IGKV1-39*01, IGKJ1*01, human, K, IGKV3-20*01, IGKJ4*01, human
T2-M5, ['Y30S_L', 'Y92S_L', 'P95S_L'], T2-Mm, TSLP_HUMAN, L, IGLV3-21*02, IGLJ2*01, human, K, IGKV3-20*01, IGKJ4*01, human

T2-L1, ['Y30S_L', 'Y92S_L', 'P95S_L'], T2-Ll, ERBB2_HUMAN, K, IGKV1-39*01, IGKJ1*01, human, K, IGKV3-20*01, IGKJ4*01, human
T2-K5, ['A50S_L'], T2-Kk, DDR1_HUMAN, K, IGKV4-68*01, IGKJ2*01, mouse, K, IGKV1-39*01, IGKJ4*01, human
T2-M7, ['A50S_L'], T2-Mm, TSLP_HUMAN, L, IGLV3-21*02, IGLJ2*01, human, K, IGKV1-39*01, IGKJ4*01, human

Supplementary Table S9 — Lineage/provenance of light chains for each design.
Per-design lineage metadata covering benchmark design linkage, antigen UniProt accession, original vs new VL types (κ/λ), V/J gene calls, and species. This table documents cLC swaps relative to parental lineages and supports immunogenetic traceability.
Abbreviations: UniProt, Universal Protein Resource; VL, light chain variable region.

| Design ID | Mutation | VH Sequence | VL Sequence |
|---|---|---|---|
| T1-A1 | | EVQLVESGGGLVQPGGSLRLSCAASGFTISDYWIHWVRQAPGKGLEWVAGITPAGGYTYYADSVKGRFTISADTSKNTAYLQMNSLRAEDTAVYYCARFVFFLPYAMDYWGQGTLVTVSS | DIQMTQSPSSLSASVGDRVTITCRASQSISSYLNWYQQKPGEAPKLLIYSASVLQSGVPSRFSGSGSGTDFTLTISSLQPEDFATYYCQQSVMIPMTFGQGTKVEIK |
| T1-A2 | | EVQLVESGGGLVQPGGSLRLSCAASGFTISDYWIHWVRQAPGKGLEWVAGITPAGGYTYYADSVKGRFTISADTSKNTAYLQMNSLRAEDTAVYYCARFVFFLPYAMDYWGQGTLVTVSS | DIQMTQSPSSLSASVGDRVTITCRASQGISSYLAWYQQKPGKAPKLLIYAASSLQSGVPSRFSGSGSGTDFTLTISSLQPEDFAVYYCQQHGNLPYTFGDGTKVEIK |
| T1-A3 | | EVQLVESGGGLVQPGGSLRLSCAASGFTISDYWIHWVRQAPGKGLEWVAGITPAGGYTYYADSVKGRFTISADTSKNTAYLQMNSLRAEDTAVYYCARFVFFLPYAMDYWGQGTLVTVSS | DIQMTQSPSSLSASVGDRVTITCRASKTISKYLAWYQQKPGKAPKLLIYSGSTLQSGVPSRFSGSGSGTDFTLTISSLQPEDFATYYCQQHNEYPLTFGQGTKVEIK |
| T1-A4 | | EVQLVESGGGLVQPGGSLRLSCAASGFTISDYWIHWVRQAPGKGLEWVAGITPAGGYTYYADSVKGRFTISADTSKNTAYLQMNSLRAEDTAVYYCARFVFFLPYAMDYWGQGTLVTVSS | DIQMTQSPSSLSASVGDRVTITCRASQSISSYLNWYQQKPGKAPKLLIYAASSLQSGVPSRFSGSGSGTDFTLTISSLQPEDFATYYCQQSDSYPLTFGGGTKVEIK |
| T1-G1 | | QVQLVQSGAEVKKPGASVKLSCKASGYTFTAYYINWVRQAPGQGLEWIGRIYPGSGYTSYAQKFQGRATLTADESTSTAYMELSSLRSEDTAVYFCARPPVYYDSAWFAYWGQGTLVTVSS | DIQMTQSPSSLSASVGDRVTITCRASQDVSTAVAWYQQKPGKAPKLLIYSASFLYSGVPSRFSGSGSGTDFTLTISSLQPEDFATYYCQQSYTTPPTFGQGTKVEIK |
| T1-G2 | | QVQLVQSGAEVKKPGASVKLSCKASGYTFTAYYINWVRQAPGQGLEWIGRIYPGSGYTSYAQKFQGRATLTADESTSTAYMELSSLRSEDTAVYFCARPPVYYDSAWFAYWGQGTLVTVSS | DIQMTQSPSSLSASVGDRVTITVRASQSISSYLNWYQQKPGEAPKLLIYSASVLQSGVPSRFSGSGSGTDFTLTISSLQPEDFATYYAQQSVMIPMTFGQGTKVEIK |
| T1-G3 | | QVQLVQSGAEVKKPGASVKLSCKASGYTFTAYYINWVRQAPGQGLEWIGRIYPGSGYTSYAQKFQGRATLTADESTSTAYMELSSLRSEDTAVYFCARPPVYYDSAWFAYWGQGTLVTVSS | DIQMTQSPSSLSASVGDRVTITCRASQGISSYLAWYQQKPGKAPKLLIYAASSLQSGVPSRFSGSGSGTDFTLTISSLQPEDFAVYYCQQHGNLPYTFGDGTKVEIK |
| T1-G4 | | QVQLVQSGAEVKKPGASVKLSCKASGYTFTAYYINWVRQAPGQGLEWIGRIYPGSGYTSYAQKFQGRATLTADESTSTAYMELSSLRSEDTAVYFCARPPVYYDSAWFAYWGQGTLVTVSS | DIQMTQSPSSLSASVGDRVTITCRASQDISNYLNWYQQKPGKAPKLLLYYTSRLHSGVPSRFSGSGSGTDYTLTISSLQPEDFATYFCQQGAGFPYTFGGGTKVEIK |
| T1-C1 | | QVQLVQSGAEVKKPGSSVKVSCKASGYAFSSYWMNWVRQAPGQGLEWMGQIWPGDSDTNYAQKFQGRVTITADESTSTAYMELSSLRSEDTAVYYCARRETTTVGRYYYAMDYWGQGTTVTVSS | DIQMTQSPSSLSASVGDRVTITCRASQGISSYLAWYQQKPGKAPKLLIYAASSLQSGVPSRFSGSGSGTDFTLTISSLQPEDFAVYYCQQHGNLPYTFGDGTKVEIK |
| T1-C2 | | QVQLVQSGAEVKKPGSSVKVSCKASGYAFSSYWMNWVRQAPGQGLEWMGQIWPGDSDTNYAQKFQGRVTITADESTSTAYMELSSLRSEDTAVYYCARRETTTVGRYYYAMDYWGQGTTVTVSS | DIQMTQSPSSLSASVGDRVTITCRASQFLSSFGVAWYQQKPGKAPKLLIYGASSLYSGVPSRFSGSGSGTDFTLTISSLQPEDFATYYCQQGLLSPLTFGQGTKVEIK |
| T1-C3 | | QVQLVQSGAEVKKPGSSVKVSCKASGYAFSSYWMNWVRQAPGQGLEWMGQIWPGDSDTNYAQKFQG | ELVMTQTPSSTSGAVGGTVTINCQASQSIDSNLAWFQQKPGQPPTLLIYRASNLASGVPSR |

| | | | |
|---|---|---|---|
| | | RVTITADESTSTAYMELSSLRSEDTAVYYCARRETTTVGRYYYAMDYWGQGTTVTVSS | FSGSRSGTEYTLTISGVQREDAATYYCLGGVGNVSYRTSFGGGTEVVVK |
| T1-C4 | | QVQLVQSGAEVKKPGSSVKVSCKASGYAFSSYWMNWVRQAPGQGLEWMGQIWPGDSDTNYAQKFQGRVTITADESTSTAYMELSSLRSEDTAVYYCARRETTTVGRYYYAMDYWGQGTTVTVSS | DIQMTQSPSSLSASVGDRVTITCRASQSVSSAVAWYQQKPGKAPKLLIYSASSLYSGVPSRFSGSRSGTDFTLTISSLQPEDFATYYCQQGVYLFTFGQGTKVEIK |
| T1-F1 | | EVQLLESGGGLVQPGGSLRLSCAVSGFTFNSFAMSWVRQAPGKGLEWVSAISGSGGGTYYADSVKGRFTISRDNSKNTLYLQMNSLRAEDTAVYFCAKDKILWFGEPVFDYWGQGTLVTVSS | EIVLTQSPATLSLSPGERATLSCRASKSISKYLAWYQQKPGQAPRLLIYSGSTLQSGIPARFSGSGSGTDFTLTISSLEPEDFAVYYCQQHNEYPYTFGQGTKLEIK |
| T1-E1 | | EVQLVQSGAEVKKPGASVKVSCKASGYKFTNYVMSWVRQAPGQRLEWMGYINPYNDAIKYNEKFTGRVTITRDTSASTAYMELSSLRSEDTAVYYCAREGDFYANYGRLGFAYWGQGTLVTVSS | DIQMTQSPSSLSASVGDRVTITCRASQSISSYLNWYQQKPGKAPKLLIYAASSLQSGVPSRFSGSGSGTDFTLTISSLQPEDFATYYCQQSYSTPPTFGQGTKVEIK |
| T1-E2 | | EVQLVQSGAEVKKPGASVKVSCKASGYKFTNYVMSWVRQAPGQRLEWMGYINPYNDAIKYNEKFTGRVTITRDTSASTAYMELSSLRSEDTAVYYCAREGDFYANYGRLGFAYWGQGTLVTVSS | DIQMTQSPSSLSASVGDRVTITCRASQSVSSAVAWYQQKPGKAPKLLIYSASSLYSGVPSRFSGSRSGTDFTLTISSLQPEDFATYYCQQGVYLFTFGQGTKVEIK |
| T1-E3 | | EVQLVQSGAEVKKPGASVKVSCKASGYKFTNYVMSWVRQAPGQRLEWMGYINPYNDAIKYNEKFTGRVTITRDTSASTAYMELSSLRSEDTAVYYCAREGDFYANYGRLGFAYWGQGTLVTVSS | DIQMTQSPSSLSASVGDRVTITCRASQSISSYLNWYQQKPGKAPKLLIYAASSLQSGVPSRFSGSGSGTDFTLTISSLQPEDFATYYCQQSDSYPLTFGGGTKVEIK |
| T1-G5 | | QVQLVQSGAEVKKPGASVKLSCKASGYTFTAYYINWVRQAPGQGLEWIGRIYPGSGYTSYAQKFQGRATLTADESTSTAYMELSSLRSEDTAVYFCARPPVYYDSAWFAYWGQGTLVTVSS | QIVLTQSPAIMSASPGEKVTMTCSASSSVSYMNWYQQKSGTSPKRWIYDTSKLASGVPAHFRGSGSGTSYSLTISGMEAEDAATYYCQQWSSNPFTFGSGTKLEIN |
| T1-G6 | | QVQLVQSGAEVKKPGASVKLSCKASGYTFTAYYINWVRQAPGQGLEWIGRIYPGSGYTSYAQKFQGRATLTADESTSTAYMELSSLRSEDTAVYFCARPPVYYDSAWFAYWGQGTLVTVSS | DIVMTQSPDSLAVSLGERATINCKSSQSLLNARTGKNYLAWYQQKPGQPPKLLIYWASTRESGVPDRFSGSGSGTDFTLTISSLQAEDVAVYYCKQSYSRRTFGGGTKVEIK |
| T1-G7 | | QVQLVQSGAEVKKPGASVKLSCKASGYTFTAYYINWVRQAPGQGLEWIGRIYPGSGYTSYAQKFQGRATLTADESTSTAYMELSSLRSEDTAVYFCARPPVYYDSAWFAYWGQGTLVTVSS | DIVMTQSPDSLAVSLGERATINCKSSQSLLNSRTRKNYLAWYQQKPGQPPKLLIYWASTRESGVPDRFSGSGSGTDFTLTISSLQAEDVAVYYCTQSFILRTFGQGTKVEIK |
| T1-F2 | | EVQLLESGGGLVQPGGSLRLSCAVSGFTFNSFAMSWVRQAPGKGLEWVSAISGSGGGTYYADSVKGRFTISRDNSKNTLYLQMNSLRAEDTAVYFCAKDKILWFGEPVFDYWGQGTLVTVSS | QIVLTQSPAIMSASPGEKVTMTCSASSSVSYMNWYQQKSGTSPKRWIYDTSKLASGVPAHFRGSGSGTSYSLTISGMEAEDAATYYCQQWSSNPFTFGSGTKLEIN |
| T1-F3 | | EVQLLESGGGLVQPGGSLRLSCAVSGFTFNSFAMSWVRQAPGKGLEWVSAISGSGGGTYYADSVKGRFTISRDNSKNTLYLQMNSLRAEDTAVYFCAKDKILWFGEPVFDYWGQGTLVTVSS | DIVMTQSPDSLAVSLGERATINCKSSQSLLNARTGKNYLAWYQQKPGQPPKLLIYWASTRESGVPDRFSGSGSGTDFTLTISSLQAEDVAVYYCKQSYSRRTFGGGTKVEIK |
| T1-F4 | | EVQLLESGGGLVQPGGSLRLSCAVSGFTFNSFAMSWVRQAPGKGLEWVSAISGSGGGTYYADSVKGRFTISRDNSKNTLYLQMNSLRAEDTAVYFCAKDKILWFGEPVFDYWGQGTLVTVSS | DIVMTQSPDSLAVSLGERATINCKSSQSLLNSRTRKNYLAWYQQKPGQPPKLLIYWASTRESGVPDRFSGSGSGTDFTLTISSLQAEDVAVYYCTQSFILRTFGQGTKVEIK |
| T1-D1 | | EVKLQQSGAELVRPGSSVKISCKASGYAFSSYWMNWVKQRPGQGLEWIGQIYPGDGDTNYNGKFKGQATLTADKSSSTAYMQLSGLTSEDSAVYFCARKTISSVVDFYFDYWGQGTTVTVSS | QIVLTQSPAIMSASPGEKVTMTCSASSSVSYMNWYQQKSGTSPKRWIYDTSKLASGVPAHFRGSGSGTSYSLTISGMEAEDAATYYCQQWSSNPFTFGSGTKLEIN |
| T1-D2 | | EVKLQQSGAELVRPGSSVKISCKASGYAFSSYWMNWVKQRPGQGLEWIGQIYPGDGDTNYNGKFKGQATLTADKSSSTAYMQLSGLTSEDSAVYFCARKTISSVVDFYFDYWGQGTTVTVSS | DIVMTQSPDSLAVSLGERATINCKSSQSLLNARTGKNYLAWYQQKPGQPPKLLIYWASTRESGVPDRFSGSGSGTDFTLTISSLQAEDVAVYYCKQSYSRRTFGGGTKVEIK |
| T1-D3 | | EVKLQQSGAELVRPGSSVKISCKASGYAFSSYWMNWVKQRPGQGLEWIGQIYPGDGDTNYNGKFKGQ | DIVMTQSPDSLAVSLGERATINCKSSQSLLNSRTRKNYLAWYQQKPGQPPKLLIYWASTR |

| | | | |
|---|---|---|---|
| | | ATLTADKSSSTAYMQLSGLTSEDSAVYFCARKTISSVVDFYFDYWGQGTTVTVSS | ESGVPDRFSGSGSGTDFTLTISSLQAEDVAVYYCTQSFILRTFGQGTKVEIK |
| T1-G8 | | QVQLVQSGAEVKKPGASVKLSCKASGYTFTAYYINWVRQAPGQGLEWIGRIYPGSGYTSYAQKFQGRATLTADESTSTAYMELSSLRSEDTAVYFCARPPVYYDSAWFAYWGQGTLVTVSS | EIVMTQSPATLSVSPGERATLSCRASQSVSSNLAWYQQKPGQAPRLLIYGASTRATGIPARFSGSGSGTEFTLTISSLQSEDFAVYYCQHYINWPLTFGGGTKVEIK |
| T1-F5 | | EVQLLESGGGLVQPGGSLRLSCAVSGFTFNSFAMSWVRQAPGKGLEWVSAISGSGGGTYYADSVKGRFTISRDNSKNTLYLQMNSLRAEDTAVYFCAKDKILWFGEPVFDYWGQGTLVTVSS | EIVMTQSPATLSVSPGERATLSCRASQSVSSNLAWYQQKPGQAPRLLIYGASTRATGIPARFSGSGSGTEFTLTISSLQSEDFAVYYCQHYINWPLTFGGGTKVEIK |
| T1-F6 | | EVQLLESGGGLVQPGGSLRLSCAVSGFTFNSFAMSWVRQAPGKGLEWVSAISGSGGGTYYADSVKGRFTISRDNSKNTLYLQMNSLRAEDTAVYFCAKDKILWFGEPVFDYWGQGTLVTVSS | DIQMTQSPSSLSASVGDRVTITCRASQSISSYLNWYQQKPGKAPKLLIYAASSLQSGVPSRFSGSGSGTDFTLTISSLQPEDFATYYCQQSYSTPPTFGQGTKVEIK |
| T1-D4 | | EVKLQQSGAELVRPGSSVKISCKASGYAFSSYWMNWVKQRPGQGLEWIGQIYPGDGDTNYNGKFKGQATLTADKSSSTAYMQLSGLTSEDSAVYFCARKTISSVVDFYFDYWGQGTTVTVSS | EIVMTQSPATLSVSPGERATLSCRASQSVSSNLAWYQQKPGQAPRLLIYGASTRATGIPARFSGSGSGTEFTLTISSLQSEDFAVYYCQHYINWPLTFGGGTKVEIK |
| T1-D5 | | EVKLQQSGAELVRPGSSVKISCKASGYAFSSYWMNWVKQRPGQGLEWIGQIYPGDGDTNYNGKFKGQATLTADKSSSTAYMQLSGLTSEDSAVYFCARKTISSVVDFYFDYWGQGTTVTVSS | DIQMTQSPSSLSASVGDRVTITCRASQSISSYLNWYQQKPGKAPKLLIYAASSLQSGVPSRFSGSGSGTDFTLTISSLQPEDFATYYCQQSYSTPPTFGQGTKVEIK |
| T1-A5 | | EVQLVESGGGLVQPGGSLRLSCAASGFTISDYWIHWVRQAPGKGLEWVAGITPAGGYTYYADSVKGRFTISADTSKNTAYLQMNSLRAEDTAVYYCARFVFFLPYAMDYWGQGTLVTVSS | ELVMTQTPSSTSGAVGGTVTINCQASQSIDSNLAWFQQKPGQPPTLLIYRASNLASGVPSRFSGSRSGTEYTLTISGVQREDAATYYCLGGVGNVSYRTSFGGGTEVVVK |
| T1-G9 | | QVQLVQSGAEVKKPGASVKLSCKASGYTFTAYYINWVRQAPGQGLEWIGRIYPGSGYTSYAQKFQGRATLTADESTSTAYMELSSLRSEDTAVYFCARPPVYYDSAWFAYWGQGTLVTVSS | DIQLTQSPSFLSASVGDRVTITCKASQSVDYSGDSYLNWYQQKPGKAPKLLIYDASNLVSGVPSRFSGSGSGTEFTLTISSLQPEDFATYYCQQSTENPWTFGGGTKLEIK |
| T1-G10 | | QVQLVQSGAEVKKPGASVKLSCKASGYTFTAYYINWVRQAPGQGLEWIGRIYPGSGYTSYAQKFQGRATLTADESTSTAYMELSSLRSEDTAVYFCARPPVYYDSAWFAYWGQGTLVTVSS | ELVMTQTPSSTSGAVGGTVTINCQASQSIDSNLAWFQQKPGQPPTLLIYRASNLASGVPSRFSGSRSGTEYTLTISGVQREDAATYYCLGGVGNVSYRTSFGGGTEVVVK |
| T1-B1 | | EVQLVESGGGLVQPGGSLRLSCAASGYDFDNYGMNWVRQAPGKGLEWVGWINTYTGEPTYAADFKRRFTFSLDTSKSTAYLQMNSLRAEDTAVYYCAKYPHYYGSSHWYFDVWGQGTLVTVSS | DIQMTQSPSSLSASVGDRVTITCRASQSISSYLNWYQQKPGEAPKLLIYSASVLQSGVPSRFSGSGSGTDFTLTISSLQPEDFATYYCQQSVMIPMTFGQGTKVEIK |
| T1-B2 | | EVQLVESGGGLVQPGGSLRLSCAASGYDFDNYGMNWVRQAPGKGLEWVGWINTYTGEPTYAADFKRRFTFSLDTSKSTAYLQMNSLRAEDTAVYYCAKYPHYYGSSHWYFDVWGQGTLVTVSS | DIQMTQSPSSLSASVGDRVTITVRASQSISSYLNWYQQKPGEAPKLLIYSASVLQSGVPSRFSGSGSGTDFTLTISSLQPEDFATYYAQQSVMIPMTFGQGTKVEIK |
| T1-B3 | | EVQLVESGGGLVQPGGSLRLSCAASGYDFDNYGMNWVRQAPGKGLEWVGWINTYTGEPTYAADFKRRFTFSLDTSKSTAYLQMNSLRAEDTAVYYCAKYPHYYGSSHWYFDVWGQGTLVTVSS | DIQMTQSPSSLSASVGDRVTITCRASQGISSYLAWYQQKPGKAPKLLIYAASSLQSGVPSRFSGSGSGTDFTLTISSLQPEDFAVYYCQQHGNLPYTFGDGTKVEIK |
| T1-B4 | | EVQLVESGGGLVQPGGSLRLSCAASGYDFDNYGMNWVRQAPGKGLEWVGWINTYTGEPTYAADFKRRFTFSLDTSKSTAYLQMNSLRAEDTAVYYCAKYPHYYGSSHWYFDVWGQGTLVTVSS | DIQMTQSPSSLSASVGDRVTITCRASKTISKYLAWYQQKPGKAPKLLIYSGSTLQSGVPSRFSGSGSGTDFTLTISSLQPEDFATYYCQQHNEYPLTFGQGTKVEIK |
| T1-B5 | | EVQLVESGGGLVQPGGSLRLSCAASGYDFDNYGMNWVRQAPGKGLEWVGWINTYTGEPTYAADFKRRFTFSLDTSKSTAYLQMNSLRAEDTAVYYCAKYPHYYGSSHWYFDVWGQGTLVTVSS | DIQMTQSPSSLSASVGDRVTITCRASQSISTWLAWYQQKPGKAPKLLIYKASNLHTGVPSRFSGSGSGTEFSLTISGLQPDDFATYYCQQYNSYSRTFGQGTKVEIK |
| T1-B6 | | EVQLVESGGGLVQPGGSLRLSCAASGYDFDNYGMNWVRQAPGKGLEWVGWINTYTGEPTYAADFKRRF | DIQMTQSPSSLSASVGDRVTITCRASQSISSYLNWYQQKPGKAPKLLIYAASSLQSGVPSR |

| | | | |
|---|---|---|---|
| | | TFSLDTSKSTAYLQMNSLRAEDTAVYYCAKYPHYY GSSHWYFDVWGQGTLVTVSS | FSGSGSGTDFTLTISSLQPEDFATYYCQQSY STPPTFGQGTKVEIK |
| T1-B 7 | | EVQLVESGGGLVQPGGSLRLSCAASGYDFDNYGM NWVRQAPGKGLEWVGWINTYTGEPTYAADFKRRF TFSLDTSKSTAYLQMNSLRAEDTAVYYCAKYPHYY GSSHWYFDVWGQGTLVTVSS | DIQLTQSPSFLSASVGDRVTITCKASQSVDY SGDSYLNWYQQKPGKAPKLLIYDASNLVSG VPSRFSGSGSGTEFTLTISSLQPEDFATYYC QQSTENPWTFGGGTKLEIK |
| T1-B 8 | | EVQLVESGGGLVQPGGSLRLSCAASGYDFDNYGM NWVRQAPGKGLEWVGWINTYTGEPTYAADFKRRF TFSLDTSKSTAYLQMNSLRAEDTAVYYCAKYPHYY GSSHWYFDVWGQGTLVTVSS | ELVMTQTPSSTSGAVGGTVTINCQASQSIDS NLAWFQQKPGQPPTLLIYRASNLASGVPSR FSGSRSGTEYTLTISGVQREDAATYYCLGGV GNVSYRTSFGGGTEVVVK |
| T1-B 9 | | EVQLVESGGGLVQPGGSLRLSCAASGYDFDNYGM NWVRQAPGKGLEWVGWINTYTGEPTYAADFKRRF TFSLDTSKSTAYLQMNSLRAEDTAVYYCAKYPHYY GSSHWYFDVWGQGTLVTVSS | DIQMTQSPSSLSASVGDRVTITCRASQSVS SAVAWYQQKPGKAPKLLIYSASSLYSGVPS RFSGSRSGTDFTLTISSLQPEDFATYYCQQG VYLFTFGQGTKVEIK |
| T1-B 10 | | EVQLVESGGGLVQPGGSLRLSCAASGYDFDNYGM NWVRQAPGKGLEWVGWINTYTGEPTYAADFKRRF TFSLDTSKSTAYLQMNSLRAEDTAVYYCAKYPHYY GSSHWYFDVWGQGTLVTVSS | DIQMTQSPSSLSASVGDRVTITCRASQDISN YLNWYQQKPGKAPKLLLYYTSRLHSGVPS RFSGSGSGTDYTLTISSLQPEDFATYFCQQG AGFPYTFGGGTKVEIK |
| T1-B 11 | | EVQLVESGGGLVQPGGSLRLSCAASGYDFDNYGM NWVRQAPGKGLEWVGWINTYTGEPTYAADFKRRF TFSLDTSKSTAYLQMNSLRAEDTAVYYCAKYPHYY GSSHWYFDVWGQGTLVTVSS | DIQMTQSPSSLSASVGDRVTITCRASQSISS YLNWYQQKPGKAPKLLIYAASSLQSGVPSR FSGSGSGTDFTLTISSLQPEDFATYYCQQSD SYPLTFGGGTKVEIK |
| T1-B 12 | | EVQLVESGGGLVQPGGSLRLSCAASGYDFDNYGM NWVRQAPGKGLEWVGWINTYTGEPTYAADFKRRF TFSLDTSKSTAYLQMNSLRAEDTAVYYCAKYPHYY GSSHWYFDVWGQGTLVTVSS | DIQMTQSPSTLSASAGDRVTISCRASQSISS WLAWYQQKPGKAPKLLIYKASSLESGVPSR FSGSGSGTEFTLTISSLQPDDFATYYCQEYN SYIRTFGQGTKVEIK |
| T1-B 13 | | EVQLVESGGGLVQPGGSLRLSCAASGYDFDNYGM NWVRQAPGKGLEWVGWINTYTGEPTYAADFKRRF TFSLDTSKSTAYLQMNSLRAEDTAVYYCAKYPHYY GSSHWYFDVWGQGTLVTVSS | DIQMTQSPSSLSASVGDRVTITCRASQSVS SAVAWYQQKPGKAPKLLIYSSLYSGVPSRF SGSRSGTDFTLTISSLQPEDFATYYCQQSSS SLITFGQGTKVEIK |
| T1-F 7 | | EVQLLESGGGLVQPGGSLRLSCAVSGFTFNSFAM SWVRQAPGKGLEWVSAISGSGGGTYYADSVKGRF TISRDNSKNTLYLQMNSLRAEDTAVYFCAKDKILWF GEPVFDYWGQGTLVTVSS | EIVLTQSPATLSLSPGERATLSCRASQNVSS FLAWYQHKPGQAPRLLIYDASSRATDIPIRF SGSGSGTDFTLTISGLEPEDFAVYYCQQRR SWPPLTFGGGTKVEIK |
| T1-F 8 | | EVQLLESGGGLVQPGGSLRLSCAVSGFTFNSFAM SWVRQAPGKGLEWVSAISGSGGGTYYADSVKGRF TISRDNSKNTLYLQMNSLRAEDTAVYFCAKDKILWF GEPVFDYWGQGTLVTVSS | EIVLTQSPGTLSLSPGERATLSCRASQSVSR SYLAWYQQKRGQAPRLLIYGASSRATGIPD RFSGDGSGTDFTLSISRLEPEDFAVYYCHQY DMSPFTFGPGTKVDIK |
| T1-E 4 | | EVQLVQSGAEVKKPGASVKVSCKASGYKFTNYVM SWVRQAPGQRLEWMGYINPYNDAIKYNEKFTGRV TITRDTSASTAYMELSSLRSEDTAVYYCAREGDFYA NYGRLGFAYWGQGTLVTVSS | ELVMTQTPSSTSGAVGGTVTINCQASQSIDS NLAWFQQKPGQPPTLLIYRASNLASGVPSR FSGSRSGTEYTLTISGVQREDAATYYCLGGV GNVSYRTSFGGGTEVVVK |
| T2-K 1 | | QVQLQESGAELVRPGASVKLSCKASGYTFSISWIN WVKQRPGQGLEWIGNIYPSGGYTNYNQKFKDKAT LTVDKSSNTAYIQLSSPTSEDSAVYYCTRGYGHLDY WGQGTTLTVSA | QSVLTQPPSVSVAPGQTARISCSGDNIGSYY VHWYQQKPGQAPVLVIYEDSERPSGIPERF SGSNSGNTATLTISGTQAEDEADYYCSSYD DPNFQVFGGGTKLTVL |
| T2-M 1 | | QMQLVESGGGVVQPGRSLRLSCAASGFTFRTYG MHWVRQAPGKGLEWVAVIWYDGSNKHYADSVKG RFTITRDNSKNTLNLQMNSLRAEDTAVYYCARAPQ WELVHEAFDIWGQGTMVTVSS | QSVLTQPPSVSVAPGQTARISCSGDNIGSYY VHWYQQKPGQAPVLVIYEDSERPSGIPERF SGSNSGNTATLTISGTQAEDEADYYCSSYD DPNFQVFGGGTKLTVL |
| T2-I1 | | EVQLLESGPGLLKPSETLSLTCTVSGGSMINYYWS WIRQPPGERPQWLGHIIYGGTTKYNPSLESRITISR DISKNQFSLRLNSVTAADTAIYYCARVAIGVSGFLNY YYYMDVWGSGTAVTVSS | SYELTQPPSVSVSPGQTASITCSGDKLGNKF TSWYQRKPGQSPVLVIYQDTKRPSGIPERF SGSTSGNTATLTISGTQAMDEADYYCQAWD SSTAWVFGGGTKLEVL |
| T2-M 2 | | QMQLVESGGGVVQPGRSLRLSCAASGFTFRTYG MHWVRQAPGKGLEWVAVIWYDGSNKHYADSVKG | SYELTQPPSVSVSPGQTASITCSGDKLGNKF TSWYQRKPGQSPVLVIYQDTKRPSGIPERF |

| | | | |
|---|---|---|---|
| | | RFTITRDNSKNTLNLQMNSLRAEDTAVYYCARAPQ WELVHEAFDIWGQGTMVTVSS | SGSTSGNTATLTISGTQAMDEADYYCQAWD SSTAWVFGGGTKLEVL |
| T2-K 2 | | QVQLQESGAELVRPGASVKLSCKASGYTFSISWIN WVKQRPGQGLEWIGNIYPSGGYTNYNQKFKDKAT LTVDKSSNTAYIQLSSPTSEDSAVYYCTRGYGHLDY WGQGTTLTVSA | EIVLTQSPATLSLSPGERATLSCRASQSISTF LAWYQHKPGQAPRLLIYDASTRATGVPARF SGSRSGTDFTLTISTLEPEDFAVYYCQQRYN WPPYTFGQGTKVEIK |
| T2-M 3 | | QMQLVESGGGVVQPGRSLRLSCAASGFTFRTYG MHWVRQAPGKGLEWVAVIWYDGSNKHYADSVKG RFTITRDNSKNTLNLQMNSLRAEDTAVYYCARAPQ WELVHEAFDIWGQGTMVTVSS | EIVLTQSPATLSLSPGERATLSCRASQSISTF LAWYQHKPGQAPRLLIYDASTRATGVPARF SGSRSGTDFTLTISTLEPEDFAVYYCQQRYN WPPYTFGQGTKVEIK |
| T2-K 3 | | QVQLQESGAELVRPGASVKLSCKASGYTFSISWIN WVKQRPGQGLEWIGNIYPSGGYTNYNQKFKDKAT LTVDKSSNTAYIQLSSPTSEDSAVYYCTRGYGHLDY WGQGTTLTVSA | EIVLTQSPATLSLSPGERATLSCRASQSVYSY LAWYQQKPGQAPRLLIYDASNRATGIPARF SGSGSGTDFTLTISSLEPEDFAVYYCQQRSN WPPFTFGPGTKVDIK |
| T2-M 4 | | QMQLVESGGGVVQPGRSLRLSCAASGFTFRTYG MHWVRQAPGKGLEWVAVIWYDGSNKHYADSVKG RFTITRDNSKNTLNLQMNSLRAEDTAVYYCARAPQ WELVHEAFDIWGQGTMVTVSS | EIVLTQSPATLSLSPGERATLSCRASQSVYSY LAWYQQKPGQAPRLLIYDASNRATGIPARF SGSGSGTDFTLTISSLEPEDFAVYYCQQRSN WPPFTFGPGTKVDIK |
| T2-M 5 | | QMQLVESGGGVVQPGRSLRLSCAASGFTFRTYG MHWVRQAPGKGLEWVAVIWYDGSNKHYADSVKG RFTITRDNSKNTLNLQMNSLRAEDTAVYYCARAPQ WELVHEAFDIWGQGTMVTVSS | ETVLTQSPGTLTLSPGERATLTCRASQSVYT YLAWYQEKPGQAPRLLIYGASSRATGIPDRF SGSGSGTEFTLTISSLQSEDFAVYYCQQYYD RPPLTFGGGTKVEIK |
| T2-L 1 | | QVQLVQSGAEVKKPGASVKLSCKASGYTFTAYYIN WVRQAPGQGLEWIGRIYPGSGYTSYAQKFQGRAT LTADESTSTAYMELSSLRSEDTAVYFCARPPVYYDS AWFAYWGQGTLVTVSS | ETVLTQSPGTLTLSPGERATLTCRASQSVYT YLAWYQEKPGQAPRLLIYGASSRATGIPDRF SGSGSGTEFTLTISSLQSEDFAVYYCQQYYD RPPLTFGGGTKVEIK |
| T2-K 4 | | QVQLQESGAELVRPGASVKLSCKASGYTFSISWIN WVKQRPGQGLEWIGNIYPSGGYTNYNQKFKDKAT LTVDKSSNTAYIQLSSPTSEDSAVYYCTRGYGHLDY WGQGTTLTVSA | DIQLTQSPSFLSASVGDRVTITCRASQDISSF LAWYQQKPGNAPKVLIYAASLLQSGVPSRF SGSGSGTDFTLTISSLQPEDFATYYCQQLNS YPLFTFGPGTKVDIK |
| T2-M 6 | | QMQLVESGGGVVQPGRSLRLSCAASGFTFRTYG MHWVRQAPGKGLEWVAVIWYDGSNKHYADSVKG RFTITRDNSKNTLNLQMNSLRAEDTAVYYCARAPQ WELVHEAFDIWGQGTMVTVSS | DIQLTQSPSFLSASVGDRVTITCRASQDISSF LAWYQQKPGNAPKVLIYAASLLQSGVPSRF SGSGSGTDFTLTISSLQPEDFATYYCQQLNS YPLFTFGPGTKVDIK |
| T2-K 5 | | QVQLQESGAELVRPGASVKLSCKASGYTFSISWIN WVKQRPGQGLEWIGNIYPSGGYTNYNQKFKDKAT LTVDKSSNTAYIQLSSPTSEDSAVYYCTRGYGHLDY WGQGTTLTVSA | DIQMTQSPSSLSASVGDRVTITCRASQSISS YLNWYQQKPGKAPKLLIYAASSLQSGVPSR FSGSGSGTDFTLTISSLQPEDFATYYCQQSY STLALTFGGGTKVEIK |
| T2-M 7 | | QMQLVESGGGVVQPGRSLRLSCAASGFTFRTYG MHWVRQAPGKGLEWVAVIWYDGSNKHYADSVKG RFTITRDNSKNTLNLQMNSLRAEDTAVYYCARAPQ WELVHEAFDIWGQGTMVTVSS | DIQMTQSPSSLSASVGDRVTITCRASQSISS YLNWYQQKPGKAPKLLIYAASSLQSGVPSR FSGSGSGTDFTLTISSLQPEDFATYYCQQSY STLALTFGGGTKVEIK |
| T2-K 6 | | QVQLQESGAELVRPGASVKLSCKASGYTFSISWIN WVKQRPGQGLEWIGNIYPSGGYTNYNQKFKDKAT LTVDKSSNTAYIQLSSPTSEDSAVYYCTRGYGHLDY WGQGTTLTVSA | AIQLTQSPSSLSASVGDRVTITCRASQGISS HLAWYQQKPGKAPKLLIFAASTLQSGVPSR FSGSGSGTDFTLTISSLQPEDFATYYCQHLN SNPPITFGQGTRLEIK |
| T2-M 8 | | QMQLVESGGGVVQPGRSLRLSCAASGFTFRTYG MHWVRQAPGKGLEWVAVIWYDGSNKHYADSVKG RFTITRDNSKNTLNLQMNSLRAEDTAVYYCARAPQ WELVHEAFDIWGQGTMVTVSS | AIQLTQSPSSLSASVGDRVTITCRASQGISS HLAWYQQKPGKAPKLLIFAASTLQSGVPSR FSGSGSGTDFTLTISSLQPEDFATYYCQHLN SNPPITFGQGTRLEIK |
| T2-J 1 | | QLQLQESGPGLVKPSETLSLTCTVSGGSISSRSYY WGWIRQPPGKGLEWIGSIYYSGFTYYQPSLKSRVT ISVDTSKNQFSLKLSSVTAADTAVYYCATGGPYGDY AHWFEPWGQGTLVTVSS | AIQLTQSPSSLSASVGDRVTITCRASQGISS HLAWYQQKPGKAPKLLIFAASTLQSGVPSR FSGSGSGTDFTLTISSLQPEDFATYYCQHLN SNPPITFGQGTRLEIK |
| T2-H 1 | | QVQLQESGPGLVKPSETLSLTCTVSGSSLTSYGVH WVRQPPGKGLEGLGVIWPGGSTNYNSALMSRVTI | AIQLTQSPSSLSASVGDRVTITCRASQGISS HLAWYQQKPGKAPKLLIFAASTLQSGVPSR |

| | | | |
|---|---|---|---|
| | | SKDNSKSQVSLKMSSLTAADTAVYYCARVTGTWYF DVWGQGTTVTVSS | FSGSGSGTDFTLTISSLQPEDFATYYCQHLN SNPPITFGQGTRLEIK |
| T2-K 7 | | QVQLQESGAELVRPGASVKLSCKASGYTFSISWIN WVKQRPGQGLEWIGNIYPSGGYTNYNQKFKDKAT LTVDKSSNTAYIQLSSPTSEDSAVYYCTRGYGHLDY WGQGTTLTVSA | DIQMTQSPSSLSASVGDRVTITCQASQDIG NYLNWYQQKPGKAPKLLIYDASHLETGVPS RFSGSGSGTDFTFTISSLQPEDIATYYCQRY DDLPSYTFGQGTKVEIK |
| T2-M 9 | | QMQLVESGGGVVQPGRSLRLSCAASGFTFRTYG MHWVRQAPGKGLEWVAVIWYDGSNKHYADSVKG RFTITRDNSKNTLNLQMNSLRAEDTAVYYCARAPQ WELVHEAFDIWGQGTMVTVSS | DIQMTQSPSSLSASVGDRVTITCQASQDIG NYLNWYQQKPGKAPKLLIYDASHLETGVPS RFSGSGSGTDFTFTISSLQPEDIATYYCQRY DDLPSYTFGQGTKVEIK |
| T2-I2 | | EVQLLESGPGLLKPSETLSLTCTVSGGSMINYYWS WIRQPPGERPQWLGHIIYGGTTKYNPSLESRITISR DISKNQFSLRLNSVTAADTAIYYCARVAIGVSGFLNY YYYMDVWGSGTAVTVSS | AIRMTQSPGTLSLSPGERATLSCRASQSISS SFLAWYQQKPGQAPRLLIYGASSRATGIPD RFSGSGSGTDFTLTISRLEPEDFAVYYCQQY GTSPRLTFGGGTKVDIK |
| T2-M 10 | | QMQLVESGGGVVQPGRSLRLSCAASGFTFRTYG MHWVRQAPGKGLEWVAVIWYDGSNKHYADSVKG RFTITRDNSKNTLNLQMNSLRAEDTAVYYCARAPQ WELVHEAFDIWGQGTMVTVSS | AIRMTQSPGTLSLSPGERATLSCRASQSISS SFLAWYQQKPGQAPRLLIYGASSRATGIPD RFSGSGSGTDFTLTISRLEPEDFAVYYCQQY GTSPRLTFGGGTKVDIK |
| T2-K 8 | | QVQLQESGAELVRPGASVKLSCKASGYTFSISWIN WVKQRPGQGLEWIGNIYPSGGYTNYNQKFKDKAT LTVDKSSNTAYIQLSSPTSEDSAVYYCTRGYGHLDY WGQGTTLTVSA | AIRMTQSPSTLSASVGDRVTITCRASQTINS WLAWYQQKPGKAPKLLIYDASNLESGVPS RFSGSGSGTEFTLTISSLQPDDFATYYCQQY ESYSPITFGQGTRLEIK |
| T2-M 11 | | QMQLVESGGGVVQPGRSLRLSCAASGFTFRTYG MHWVRQAPGKGLEWVAVIWYDGSNKHYADSVKG RFTITRDNSKNTLNLQMNSLRAEDTAVYYCARAPQ WELVHEAFDIWGQGTMVTVSS | AIRMTQSPSTLSASVGDRVTITCRASQTINS WLAWYQQKPGKAPKLLIYDASNLESGVPS RFSGSGSGTEFTLTISSLQPDDFATYYCQQY ESYSPITFGQGTRLEIK |
| T2-I3 | | EVQLLESGPGLLKPSETLSLTCTVSGGSMINYYWS WIRQPPGERPQWLGHIIYGGTTKYNPSLESRITISR DISKNQFSLRLNSVTAADTAIYYCARVAIGVSGFLNY YYYMDVWGSGTAVTVSS | EIVLTQSPGTLSLSPGERATLSCRASQSISS NYLAWYQQKPGQAPRLLIYGASSRATGIPD RFSGSGSGTDFTLSINRLEPEDFAMYYCQH YGGLSRWTFGQGTKVEIK |
| T2-M 12 | | QMQLVESGGGVVQPGRSLRLSCAASGFTFRTYG MHWVRQAPGKGLEWVAVIWYDGSNKHYADSVKG RFTITRDNSKNTLNLQMNSLRAEDTAVYYCARAPQ WELVHEAFDIWGQGTMVTVSS | EIVLTQSPGTLSLSPGERATLSCRASQSISS NYLAWYQQKPGQAPRLLIYGASSRATGIPD RFSGSGSGTDFTLSINRLEPEDFAMYYCQH YGGLSRWTFGQGTKVEIK |
| T2-I4 | | EVQLLESGPGLLKPSETLSLTCTVSGGSMINYYWS WIRQPPGERPQWLGHIIYGGTTKYNPSLESRITISR DISKNQFSLRLNSVTAADTAIYYCARVAIGVSGFLNY YYYMDVWGSGTAVTVSS | SYELTQPPSVSVSPGQTASITCSGDKLGDKY ACWYQQKPGQSPVLVIYQDNKRPSGIPERF SGSNSGNTATLTISGTQAMDEADYYCQAW DSSTAVFGGGTKLTVL |
| T2-K 9 | | QVQLQESGAELVRPGASVKLSCKASGYTFSISWIN WVKQRPGQGLEWIGNIYPSGGYTNYNQKFKDKAT LTVDKSSNTAYIQLSSPTSEDSAVYYCTRGYGHLDY WGQGTTLTVSA | SYELTQPPSVSVSPGQTASITCSGDKLGDKY ACWYQQKPGQSPVLVIYQDNKRPSGIPERF SGSNSGNTATLTISGTQAMDEADYYCQAW DSSTAVFGGGTKLTVL |
| T2-L 2 | | QVQLVQSGAEVKKPGASVKLSCKASGYTFTAYYIN WVRQAPGQGLEWIGRIYPGSGYTSYAQKFQGRAT LTADESTSTAYMELSSLRSEDTAVYFCARPPVYYDS AWFAYWGQGTLVTVSS | SYELTQPPSVSVSPGQTASITCSGDKLGDKY ACWYQQKPGQSPVLVIYQDNKRPSGIPERF SGSNSGNTATLTISGTQAMDEADYYCQAW DSSTAVFGGGTKLTVL |
| T2-K 10 | | QVQLQESGAELVRPGASVKLSCKASGYTFSISWIN WVKQRPGQGLEWIGNIYPSGGYTNYNQKFKDKAT LTVDKSSNTAYIQLSSPTSEDSAVYYCTRGYGHLDY WGQGTTLTVSA | VIQMTQSPSTLSASVGDRVTITCRASQSVST WLAWYQQKPGQGPKLLIYEASSLESGVPS RFSGSGSGTEFTLTISSLQPDDFATYYCQQY NSYSFWTFGQGTKVEIK |
| T2-M 13 | | QMQLVESGGGVVQPGRSLRLSCAASGFTFRTYG MHWVRQAPGKGLEWVAVIWYDGSNKHYADSVKG RFTITRDNSKNTLNLQMNSLRAEDTAVYYCARAPQ WELVHEAFDIWGQGTMVTVSS | VIQMTQSPSTLSASVGDRVTITCRASQSVST WLAWYQQKPGQGPKLLIYEASSLESGVPS RFSGSGSGTEFTLTISSLQPDDFATYYCQQY NSYSFWTFGQGTKVEIK |
| T2-I5 | | EVQLLESGPGLLKPSETLSLTCTVSGGSMINYYWS WIRQPPGERPQWLGHIIYGGTTKYNPSLESRITISR | DIQMTQSPSSLSASVGDRVTITCRASQSVS SAVAWYQQKPGKAPKLLIYSASSLYSGVPS |

| | | | |
|---|---|---|---|
| | | DISKNQFSLRLNSVTAADTAIYYCARVAIGVSGFLNYYYYMDVWGSGTAVTVSS | RFSGSRSGTDFTLTISSLQPEDFATYYCQQSWSAYPFTFGQGTKVEIK |
| T2-M14 | | QMQLVESGGGVVQPGRSLRLSCAASGFTFRTYGMHWVRQAPGKGLEWVAVIWYDGSNKHYADSVKGRFTITRDNSKNTLNLQMNSLRAEDTAVYYCARAPQWELVHEAFDIWGQGTMVTVSS | DIQMTQSPSSLSASVGDRVTITCRASQSVSSAVAWYQQKPGKAPKLLIYSASSLYSGVPSRFSGSRSGTDFTLTISSLQPEDFATYYCQQSWSAYPFTFGQGTKVEIK |
| T2-H2 | | QVQLQESGPGLVKPSETLSLTCTVSGSSLTSYGVHWVRQPPGKGLEGLGVIWPGGSTNYNSALMSRVTISKDNSKSQVSLKMSSLTAADTAVYYCARVTGTWYFDVWGQGTTVTVSS | DIQMTQSPSSLSASVGDRVTITCRASQSVSSAVAWYQQKPGKAPKLLIYSASSLYSGVPSRFSGSRSGTDFTLTISSLQPEDFATYYCQQSWSAYPFTFGQGTKVEIK |
| T2-K11 | | QVQLQESGAELVRPGASVKLSCKASGYTFSISWINWVKQRPGQGLEWIGNIYPSGGYTNYNQKFKDKATLTVDKSSNTAYIQLSSPTSEDSAVYYCTRGYGHLDYWGQGTTLTVSA | DIQMTQSPSSLSASVGDRVTITCRASQSVSSAVAWYQQKPGKAPKLLIYSASSLYSGVPSRFSGSRSGTDFTLTISSLQPEDFATYYCQQSEWGGLITFGQGTKVEIK |
| T2-M15 | | QMQLVESGGGVVQPGRSLRLSCAASGFTFRTYGMHWVRQAPGKGLEWVAVIWYDGSNKHYADSVKGRFTITRDNSKNTLNLQMNSLRAEDTAVYYCARAPQWELVHEAFDIWGQGTMVTVSS | DIQMTQSPSSLSASVGDRVTITCRASQSVSSAVAWYQQKPGKAPKLLIYSASSLYSGVPSRFSGSRSGTDFTLTISSLQPEDFATYYCQQSEWGGLITFGQGTKVEIK |
| T2-K1-1 | E49L_L; E52N_L | QVQLQESGAELVRPGASVKLSCKASGYTFSISWINWVKQRPGQGLEWIGNIYPSGGYTNYNQKFKDKATLTVDKSSNTAYIQLSSPTSEDSAVYYCTRGYGHLDYWGQGTTLTVSA | QSVLTQPPSVSVAPGQTARISCSGDNIGSYYVHWYQQKPGQAPVLVIYLDSNRPSGIPERFSGSNSGNTATLTISGTQAEDEADYYCSSYDDPNFQVFGGGTKLTVL |
| T2-M1-1 | E49L_L; E52N_L | QMQLVESGGGVVQPGRSLRLSCAASGFTFRTYGMHWVRQAPGKGLEWVAVIWYDGSNKHYADSVKGRFTITRDNSKNTLNLQMNSLRAEDTAVYYCARAPQWELVHEAFDIWGQGTMVTVSS | QSVLTQPPSVSVAPGQTARISCSGDNIGSYYVHWYQQKPGQAPVLVIYLDSNRPSGIPERFSGSNSGNTATLTISGTQAEDEADYYCSSYDDPNFQVFGGGTKLTVL |
| T2-K1-2 | E49L_L; E52N_L; Y31F_L | QVQLQESGAELVRPGASVKLSCKASGYTFSISWINWVKQRPGQGLEWIGNIYPSGGYTNYNQKFKDKATLTVDKSSNTAYIQLSSPTSEDSAVYYCTRGYGHLDYWGQGTTLTVSA | QSVLTQPPSVSVAPGQTARISCSGDNIGSYFVHWYQQKPGQAPVLVIYLDSNRPSGIPERFSGSNSGNTATLTISGTQAEDEADYYCSSYDDPNFQVFGGGTKLTVL |
| T2-M1-2 | E49L_L; E52N_L; Y31F_L | QMQLVESGGGVVQPGRSLRLSCAASGFTFRTYGMHWVRQAPGKGLEWVAVIWYDGSNKHYADSVKGRFTITRDNSKNTLNLQMNSLRAEDTAVYYCARAPQWELVHEAFDIWGQGTMVTVSS | QSVLTQPPSVSVAPGQTARISCSGDNIGSYFVHWYQQKPGQAPVLVIYLDSNRPSGIPERFSGSNSGNTATLTISGTQAEDEADYYCSSYDDPNFQVFGGGTKLTVL |
| T2-K1-3 | E49L_L; Y31F_L | QVQLQESGAELVRPGASVKLSCKASGYTFSISWINWVKQRPGQGLEWIGNIYPSGGYTNYNQKFKDKATLTVDKSSNTAYIQLSSPTSEDSAVYYCTRGYGHLDYWGQGTTLTVSA | QSVLTQPPSVSVAPGQTARISCSGDNIGSYFVHWYQQKPGQAPVLVIYLDSERPSGIPERFSGSNSGNTATLTISGTQAEDEADYYCSSYDDPNFQVFGGGTKLTVL |
| T2-M1-3 | E49L_L; Y31F_L | QMQLVESGGGVVQPGRSLRLSCAASGFTFRTYGMHWVRQAPGKGLEWVAVIWYDGSNKHYADSVKGRFTITRDNSKNTLNLQMNSLRAEDTAVYYCARAPQWELVHEAFDIWGQGTMVTVSS | QSVLTQPPSVSVAPGQTARISCSGDNIGSYFVHWYQQKPGQAPVLVIYLDSERPSGIPERFSGSNSGNTATLTISGTQAEDEADYYCSSYDDPNFQVFGGGTKLTVL |
| T2-K1-4 | E52N_L | QVQLQESGAELVRPGASVKLSCKASGYTFSISWINWVKQRPGQGLEWIGNIYPSGGYTNYNQKFKDKATLTVDKSSNTAYIQLSSPTSEDSAVYYCTRGYGHLDYWGQGTTLTVSA | QSVLTQPPSVSVAPGQTARISCSGDNIGSYYVHWYQQKPGQAPVLVIYEDSNRPSGIPERFSGSNSGNTATLTISGTQAEDEADYYCSSYDDPNFQVFGGGTKLTVL |
| T2-M1-4 | E52N_L | QMQLVESGGGVVQPGRSLRLSCAASGFTFRTYGMHWVRQAPGKGLEWVAVIWYDGSNKHYADSVKGRFTITRDNSKNTLNLQMNSLRAEDTAVYYCARAPQWELVHEAFDIWGQGTMVTVSS | QSVLTQPPSVSVAPGQTARISCSGDNIGSYYVHWYQQKPGQAPVLVIYEDSNRPSGIPERFSGSNSGNTATLTISGTQAEDEADYYCSSYDDPNFQVFGGGTKLTVL |
| T2-K1-5 | E52N_L; | QVQLQESGAELVRPGASVKLSCKASGYTFSISWINWVKQRPGQGLEWIGNIYPSGGYTNYNQKFKDKAT | QSVLTQPPSVSVAPGQTARISCSGDNIGSYFVHWYQQKPGQAPVLVIYEDSNRPSGIPERF |

| | | | |
|---|---|---|---|
| | Y31F<br>_L | LTVDKSSNTAYIQLSSPTSEDSAVYYCTRGYGHLDY<br>WGQGTTLTVSA | SGSNSGNTATLTISGTQAEDEADYYCSSYD<br>DPNFQVFGGGTKLTVL |
| T2-M<br>1-5 | E52N<br>_L;<br>Y31F<br>_L | QMQLVESGGGVVQPGRSLRLSCAASGFTFRTYG<br>MHWVRQAPGKGLEWVAVIWYDGSNKHYADSVKG<br>RFTITRDNSKNTLNLQMNSLRAEDTAVYYCARAPQ<br>WELVHEAFDIWGQGTMVTVSS | QSVLTQPPSVSVAPGQTARISCSGDNIGSYF<br>VHWYQQKPGQAPVLVIYEDSNRPSGIPERF<br>SGSNSGNTATLTISGTQAEDEADYYCSSYD<br>DPNFQVFGGGTKLTVL |
| T2-K<br>1-6 | Y31F<br>_L | QVQLQESGAELVRPGASVKLSCKASGYTFSISWIN<br>WVKQRPGQGLEWIGNIYPSGGYTNYNQKFKDKAT<br>LTVDKSSNTAYIQLSSPTSEDSAVYYCTRGYGHLDY<br>WGQGTTLTVSA | QSVLTQPPSVSVAPGQTARISCSGDNIGSYF<br>VHWYQQKPGQAPVLVIYEDSERPSGIPERF<br>SGSNSGNTATLTISGTQAEDEADYYCSSYD<br>DPNFQVFGGGTKLTVL |
| T2-M<br>1-6 | Y31F<br>_L | QMQLVESGGGVVQPGRSLRLSCAASGFTFRTYG<br>MHWVRQAPGKGLEWVAVIWYDGSNKHYADSVKG<br>RFTITRDNSKNTLNLQMNSLRAEDTAVYYCARAPQ<br>WELVHEAFDIWGQGTMVTVSS | QSVLTQPPSVSVAPGQTARISCSGDNIGSYF<br>VHWYQQKPGQAPVLVIYEDSERPSGIPERF<br>SGSNSGNTATLTISGTQAEDEADYYCSSYD<br>DPNFQVFGGGTKLTVL |
| T2-K<br>2-1 | T53N<br>_L | QVQLQESGAELVRPGASVKLSCKASGYTFSISWIN<br>WVKQRPGQGLEWIGNIYPSGGYTNYNQKFKDKAT<br>LTVDKSSNTAYIQLSSPTSEDSAVYYCTRGYGHLDY<br>WGQGTTLTVSA | EIVLTQSPATLSLSPGERATLSCRASQSISTF<br>LAWYQHKPGQAPRLLIYDASNRATGVPARF<br>SGSRSGTDFTLTISTLEPEDFAVYYCQQRYN<br>WPPYTFGQGTKVEIK |
| T2-M<br>3-1 | T53N<br>_L | QMQLVESGGGVVQPGRSLRLSCAASGFTFRTYG<br>MHWVRQAPGKGLEWVAVIWYDGSNKHYADSVKG<br>RFTITRDNSKNTLNLQMNSLRAEDTAVYYCARAPQ<br>WELVHEAFDIWGQGTMVTVSS | EIVLTQSPATLSLSPGERATLSCRASQSISTF<br>LAWYQHKPGQAPRLLIYDASNRATGVPARF<br>SGSRSGTDFTLTISTLEPEDFAVYYCQQRYN<br>WPPYTFGQGTKVEIK |
| T2-K<br>3-1 | W94<br>N_L | QVQLQESGAELVRPGASVKLSCKASGYTFSISWIN<br>WVKQRPGQGLEWIGNIYPSGGYTNYNQKFKDKAT<br>LTVDKSSNTAYIQLSSPTSEDSAVYYCTRGYGHLDY<br>WGQGTTLTVSA | EIVLTQSPATLSLSPGERATLSCRASQSVYSY<br>LAWYQQKPGQAPRLLIYDASNRATGIPARF<br>SGSGSGTDFTLTISSLEPEDFAVYYCQQRSN<br>NPPFTFGPGTKVDIK |
| T2-M<br>4-1 | W94<br>N_L | QMQLVESGGGVVQPGRSLRLSCAASGFTFRTYG<br>MHWVRQAPGKGLEWVAVIWYDGSNKHYADSVKG<br>RFTITRDNSKNTLNLQMNSLRAEDTAVYYCARAPQ<br>WELVHEAFDIWGQGTMVTVSS | EIVLTQSPATLSLSPGERATLSCRASQSVYSY<br>LAWYQQKPGQAPRLLIYDASNRATGIPARF<br>SGSGSGTDFTLTISSLEPEDFAVYYCQQRSN<br>NPPFTFGPGTKVDIK |
| T2-K<br>3-2 | Y30T<br>_L;<br>W94<br>N_L | QVQLQESGAELVRPGASVKLSCKASGYTFSISWIN<br>WVKQRPGQGLEWIGNIYPSGGYTNYNQKFKDKAT<br>LTVDKSSNTAYIQLSSPTSEDSAVYYCTRGYGHLDY<br>WGQGTTLTVSA | EIVLTQSPATLSLSPGERATLSCRASQSVTSY<br>LAWYQQKPGQAPRLLIYDASNRATGIPARF<br>SGSGSGTDFTLTISSLEPEDFAVYYCQQRSN<br>NPPFTFGPGTKVDIK |
| T2-M<br>4-2 | Y30T<br>_L;<br>W94<br>N_L | QMQLVESGGGVVQPGRSLRLSCAASGFTFRTYG<br>MHWVRQAPGKGLEWVAVIWYDGSNKHYADSVKG<br>RFTITRDNSKNTLNLQMNSLRAEDTAVYYCARAPQ<br>WELVHEAFDIWGQGTMVTVSS | EIVLTQSPATLSLSPGERATLSCRASQSVTSY<br>LAWYQQKPGQAPRLLIYDASNRATGIPARF<br>SGSGSGTDFTLTISSLEPEDFAVYYCQQRSN<br>NPPFTFGPGTKVDIK |
| T2-K<br>3-3 | Y32F<br>_L | QVQLQESGAELVRPGASVKLSCKASGYTFSISWIN<br>WVKQRPGQGLEWIGNIYPSGGYTNYNQKFKDKAT<br>LTVDKSSNTAYIQLSSPTSEDSAVYYCTRGYGHLDY<br>WGQGTTLTVSA | EIVLTQSPATLSLSPGERATLSCRASQSVYSF<br>LAWYQQKPGQAPRLLIYDASNRATGIPARF<br>SGSGSGTDFTLTISSLEPEDFAVYYCQQRSN<br>WPPFTFGPGTKVDIK |
| T2-M<br>4-3 | Y32F<br>_L | QMQLVESGGGVVQPGRSLRLSCAASGFTFRTYG<br>MHWVRQAPGKGLEWVAVIWYDGSNKHYADSVKG<br>RFTITRDNSKNTLNLQMNSLRAEDTAVYYCARAPQ<br>WELVHEAFDIWGQGTMVTVSS | EIVLTQSPATLSLSPGERATLSCRASQSVYSF<br>LAWYQQKPGQAPRLLIYDASNRATGIPARF<br>SGSGSGTDFTLTISSLEPEDFAVYYCQQRSN<br>WPPFTFGPGTKVDIK |
| T2-K<br>3-4 | Y32F<br>_L;<br>W94<br>N_L | QVQLQESGAELVRPGASVKLSCKASGYTFSISWIN<br>WVKQRPGQGLEWIGNIYPSGGYTNYNQKFKDKAT<br>LTVDKSSNTAYIQLSSPTSEDSAVYYCTRGYGHLDY<br>WGQGTTLTVSA | EIVLTQSPATLSLSPGERATLSCRASQSVYSF<br>LAWYQQKPGQAPRLLIYDASNRATGIPARF<br>SGSGSGTDFTLTISSLEPEDFAVYYCQQRSN<br>NPPFTFGPGTKVDIK |
| T2-M<br>4-4 | Y32F<br>_L;<br>W94<br>N_L | QMQLVESGGGVVQPGRSLRLSCAASGFTFRTYG<br>MHWVRQAPGKGLEWVAVIWYDGSNKHYADSVKG<br>RFTITRDNSKNTLNLQMNSLRAEDTAVYYCARAPQ<br>WELVHEAFDIWGQGTMVTVSS | EIVLTQSPATLSLSPGERATLSCRASQSVYSF<br>LAWYQQKPGQAPRLLIYDASNRATGIPARF<br>SGSGSGTDFTLTISSLEPEDFAVYYCQQRSN<br>NPPFTFGPGTKVDIK |
| T2-K<br>3-5 | Y32F<br>_L; | QVQLQESGAELVRPGASVKLSCKASGYTFSISWIN<br>WVKQRPGQGLEWIGNIYPSGGYTNYNQKFKDKAT | EIVLTQSPATLSLSPGERATLSCRASQSVTSF<br>LAWYQQKPGQAPRLLIYDASNRATGIPARF |

| | | | |
|---|---|---|---|
| | Y30T_L; W94N_L | LTVDKSSNTAYIQLSSPTSEDSAVYYCTRGYGHLDYWGQGTTLTVSA | SGSGSGTDFTLTISSLEPEDFAVYYCQQRSNNPPFTFGPGTKVDIK |
| T2-M4-5 | Y32F_L; Y30T_L; W94N_L | QMQLVESGGGVVQPGRSLRLSCAASGFTFRTYGMHWVRQAPGKGLEWVAVIWYDGSNKHYADSVKGRFTITRDNSKNTLNLQMNSLRAEDTAVYYCARAPQWELVHEAFDIWGQGTMVTVSS | EIVLTQSPATLSLSPGERATLSCRASQSVTSFLAWYQQKPGQAPRLLIYDASNRATGIPARFSGSGSGTDFTLTISSLEPEDFAVYYCQQRSNNPPFTFGPGTKVDIK |
| T2-K5-1 | A50D_L | QVQLQESGAELVRPGASVKLSCKASGYTFSISWINWVKQRPGQGLEWIGNIYPSGGYTNYNQKFKDKATLTVDKSSNTAYIQLSSPTSEDSAVYYCTRGYGHLDYWGQGTTLTVSA | DIQMTQSPSSLSASVGDRVTITCRASQSISSYLNWYQQKPGKAPKLLIYDASSLQSGVPSRFSGSGSGTDFTLTISSLQPEDFATYYCQQSYSTLALTFGGGTKVEIK |
| T2-M7-1 | A50D_L | QMQLVESGGGVVQPGRSLRLSCAASGFTFRTYGMHWVRQAPGKGLEWVAVIWYDGSNKHYADSVKGRFTITRDNSKNTLNLQMNSLRAEDTAVYYCARAPQWELVHEAFDIWGQGTMVTVSS | DIQMTQSPSSLSASVGDRVTITCRASQSISSYLNWYQQKPGKAPKLLIYDASSLQSGVPSRFSGSGSGTDFTLTISSLQPEDFATYYCQQSYSTLALTFGGGTKVEIK |
| T2-K5-2 | A50D_L; L95H_L | QVQLQESGAELVRPGASVKLSCKASGYTFSISWINWVKQRPGQGLEWIGNIYPSGGYTNYNQKFKDKATLTVDKSSNTAYIQLSSPTSEDSAVYYCTRGYGHLDYWGQGTTLTVSA | DIQMTQSPSSLSASVGDRVTITCRASQSISSYLNWYQQKPGKAPKLLIYDASSLQSGVPSRFSGSGSGTDFTLTISSLQPEDFATYYCQQSYSTHALTFGGGTKVEIK |
| T2-M7-2 | A50D_L; L95H_L | QMQLVESGGGVVQPGRSLRLSCAASGFTFRTYGMHWVRQAPGKGLEWVAVIWYDGSNKHYADSVKGRFTITRDNSKNTLNLQMNSLRAEDTAVYYCARAPQWELVHEAFDIWGQGTMVTVSS | DIQMTQSPSSLSASVGDRVTITCRASQSISSYLNWYQQKPGKAPKLLIYDASSLQSGVPSRFSGSGSGTDFTLTISSLQPEDFATYYCQQSYSTHALTFGGGTKVEIK |
| T2-K5-3 | L95H_L | QVQLQESGAELVRPGASVKLSCKASGYTFSISWINWVKQRPGQGLEWIGNIYPSGGYTNYNQKFKDKATLTVDKSSNTAYIQLSSPTSEDSAVYYCTRGYGHLDYWGQGTTLTVSA | DIQMTQSPSSLSASVGDRVTITCRASQSISSYLNWYQQKPGKAPKLLIYAASSLQSGVPSRFSGSGSGTDFTLTISSLQPEDFATYYCQQSYSTHALTFGGGTKVEIK |
| T2-M7-3 | L95H_L | QMQLVESGGGVVQPGRSLRLSCAASGFTFRTYGMHWVRQAPGKGLEWVAVIWYDGSNKHYADSVKGRFTITRDNSKNTLNLQMNSLRAEDTAVYYCARAPQWELVHEAFDIWGQGTMVTVSS | DIQMTQSPSSLSASVGDRVTITCRASQSISSYLNWYQQKPGKAPKLLIYAASSLQSGVPSRFSGSGSGTDFTLTISSLQPEDFATYYCQQSYSTHALTFGGGTKVEIK |
| T2-K5-4 | S53N_L | QVQLQESGAELVRPGASVKLSCKASGYTFSISWINWVKQRPGQGLEWIGNIYPSGGYTNYNQKFKDKATLTVDKSSNTAYIQLSSPTSEDSAVYYCTRGYGHLDYWGQGTTLTVSA | DIQMTQSPSSLSASVGDRVTITCRASQSISSYLNWYQQKPGKAPKLLIYAASNLQSGVPSRFSGSGSGTDFTLTISSLQPEDFATYYCQQSYSTLALTFGGGTKVEIK |
| T2-M7-4 | S53N_L | QMQLVESGGGVVQPGRSLRLSCAASGFTFRTYGMHWVRQAPGKGLEWVAVIWYDGSNKHYADSVKGRFTITRDNSKNTLNLQMNSLRAEDTAVYYCARAPQWELVHEAFDIWGQGTMVTVSS | DIQMTQSPSSLSASVGDRVTITCRASQSISSYLNWYQQKPGKAPKLLIYAASNLQSGVPSRFSGSGSGTDFTLTISSLQPEDFATYYCQQSYSTLALTFGGGTKVEIK |
| T2-K7-1 | D92S_L | QVQLQESGAELVRPGASVKLSCKASGYTFSISWINWVKQRPGQGLEWIGNIYPSGGYTNYNQKFKDKATLTVDKSSNTAYIQLSSPTSEDSAVYYCTRGYGHLDYWGQGTTLTVSA | DIQMTQSPSSLSASVGDRVTITCQASQDIGNYLNWYQQKPGKAPKLLIYDASHLETGVPSRFSGSGSGTDFTFTISSLQPEDIATYYCQRYSDLPSYTFGQGTKVEIK |
| T2-M9-1 | D92S_L | QMQLVESGGGVVQPGRSLRLSCAASGFTFRTYGMHWVRQAPGKGLEWVAVIWYDGSNKHYADSVKGRFTITRDNSKNTLNLQMNSLRAEDTAVYYCARAPQWELVHEAFDIWGQGTMVTVSS | DIQMTQSPSSLSASVGDRVTITCQASQDIGNYLNWYQQKPGKAPKLLIYDASHLETGVPSRFSGSGSGTDFTFTISSLQPEDIATYYCQRYSDLPSYTFGQGTKVEIK |
| T2-K7-2 | D92S_L; D93S_L | QVQLQESGAELVRPGASVKLSCKASGYTFSISWINWVKQRPGQGLEWIGNIYPSGGYTNYNQKFKDKATLTVDKSSNTAYIQLSSPTSEDSAVYYCTRGYGHLDYWGQGTTLTVSA | DIQMTQSPSSLSASVGDRVTITCQASQDIGNYLNWYQQKPGKAPKLLIYDASHLETGVPSRFSGSGSGTDFTFTISSLQPEDIATYYCQRYSSLPSYTFGQGTKVEIK |
| T2-M9-2 | D92S_L; | QMQLVESGGGVVQPGRSLRLSCAASGFTFRTYGMHWVRQAPGKGLEWVAVIWYDGSNKHYADSVKG | DIQMTQSPSSLSASVGDRVTITCQASQDIGNYLNWYQQKPGKAPKLLIYDASHLETGVPS |

| | | | |
|---|---|---|---|
| | D93S_L | RFTITRDNSKNTLNLQMNSLRAEDTAVYYCARAPQWELVHEAFDIWGQGTMVTVSS | RFSGSGSGTDFTFTISSLQPEDIATYYCQRYSSLPSYTFGQGTKVEIK |
| T2-K7-3 | D93S_L | QVQLQESGAELVRPGASVKLSCKASGYTFSISWINWVKQRPGQGLEWIGNIYPSGGYTNYNQKFKDKATLTVDKSSNTAYIQLSSPTSEDSAVYYCTRGYGHLDYWGQGTTLTVSA | DIQMTQSPSSLSASVGDRVTITCQASQDIGNYLNWYQQKPGKAPKLLIYDASHLETGVPSRFSGSGSGTDFTFTISSLQPEDIATYYCQRYDSLPSYTFGQGTKVEIK |
| T2-M9-3 | D93S_L | QMQLVESGGGVVQPGRSLRLSCAASGFTFRTYGMHWVRQAPGKGLEWVAVIWYDGSNKHYADSVKGRFTITRDNSKNTLNLQMNSLRAEDTAVYYCARAPQWELVHEAFDIWGQGTMVTVSS | DIQMTQSPSSLSASVGDRVTITCQASQDIGNYLNWYQQKPGKAPKLLIYDASHLETGVPSRFSGSGSGTDFTFTISSLQPEDIATYYCQRYDSLPSYTFGQGTKVEIK |
| T2-I2-1 | R97H_L | EVQLLESGPGLLKPSETLSLTCTVSGGSMINYYWSWIRQPPGERPQWLGHIIYGGTTKYNPSLESRITISRDISKNQFSLRLNSVTAADTAIYYCARVAIGVSGFLNYYYYMDVWGSGTAVTVSS | AIRMTQSPGTLSLSPGERATLSCRASQSISSSFLAWYQQKPGQAPRLLIYGASSRATGIPDRFSGSGSGTDFTLTISRLEPEDFAVYYCQQYGTSPHLTFGGGTKVDIK |
| T2-M10-1 | R97H_L | QMQLVESGGGVVQPGRSLRLSCAASGFTFRTYGMHWVRQAPGKGLEWVAVIWYDGSNKHYADSVKGRFTITRDNSKNTLNLQMNSLRAEDTAVYYCARAPQWELVHEAFDIWGQGTMVTVSS | AIRMTQSPGTLSLSPGERATLSCRASQSISSSFLAWYQQKPGQAPRLLIYGASSRATGIPDRFSGSGSGTDFTLTISRLEPEDFAVYYCQQYGTSPHLTFGGGTKVDIK |
| T2-K8-1 | Y94S_L | QVQLQESGAELVRPGASVKLSCKASGYTFSISWINWVKQRPGQGLEWIGNIYPSGGYTNYNQKFKDKATLTVDKSSNTAYIQLSSPTSEDSAVYYCTRGYGHLDYWGQGTTLTVSA | AIRMTQSPSTLSASVGDRVTITCRASQTINSWLAWYQQKPGKAPKLLIYDASNLESGVPSRFSGSGSGTEFTLTISSLQPDDFATYYCQQYESSSPITFGQGTRLEIK |
| T2-M11-1 | Y94S_L | QMQLVESGGGVVQPGRSLRLSCAASGFTFRTYGMHWVRQAPGKGLEWVAVIWYDGSNKHYADSVKGRFTITRDNSKNTLNLQMNSLRAEDTAVYYCARAPQWELVHEAFDIWGQGTMVTVSS | AIRMTQSPSTLSASVGDRVTITCRASQTINSWLAWYQQKPGKAPKLLIYDASNLESGVPSRFSGSGSGTEFTLTISSLQPDDFATYYCQQYESSSPITFGQGTRLEIK |
| T2-I4-1 | C33A_L | EVQLLESGPGLLKPSETLSLTCTVSGGSMINYYWSWIRQPPGERPQWLGHIIYGGTTKYNPSLESRITISRDISKNQFSLRLNSVTAADTAIYYCARVAIGVSGFLNYYYYMDVWGSGTAVTVSS | SYELTQPPSVSVSPGQTASITCSGDKLGDKYAAWYQQKPGQSPVLVIYQDNKRPSGIPERFSGSNSGNTATLTISGTQAMDEADYYCQAWDSSTAVFGGGTKLTVL |
| T2-K9-1 | C33A_L | QVQLQESGAELVRPGASVKLSCKASGYTFSISWINWVKQRPGQGLEWIGNIYPSGGYTNYNQKFKDKATLTVDKSSNTAYIQLSSPTSEDSAVYYCTRGYGHLDYWGQGTTLTVSA | SYELTQPPSVSVSPGQTASITCSGDKLGDKYAAWYQQKPGQSPVLVIYQDNKRPSGIPERFSGSNSGNTATLTISGTQAMDEADYYCQAWDSSTAVFGGGTKLTVL |
| T2-L2-1 | C33A_L | QVQLVQSGAEVKKPGASVKLSCKASGYTFTAYYINWVRQAPGQGLEWIGRIYPGSGYTSYAQKFQGRATLTADESTSTAYMELSSLRSEDTAVYFCARPPVYYDSAWFAYWGQGTLVTVSS | SYELTQPPSVSVSPGQTASITCSGDKLGDKYAAWYQQKPGQSPVLVIYQDNKRPSGIPERFSGSNSGNTATLTISGTQAMDEADYYCQAWDSSTAVFGGGTKLTVL |
| T2-I4-2 | C33S_L | EVQLLESGPGLLKPSETLSLTCTVSGGSMINYYWSWIRQPPGERPQWLGHIIYGGTTKYNPSLESRITISRDISKNQFSLRLNSVTAADTAIYYCARVAIGVSGFLNYYYYMDVWGSGTAVTVSS | SYELTQPPSVSVSPGQTASITCSGDKLGDKYASWYQQKPGQSPVLVIYQDNKRPSGIPERFSGSNSGNTATLTISGTQAMDEADYYCQAWDSSTAVFGGGTKLTVL |
| T2-K9-2 | C33S_L | QVQLQESGAELVRPGASVKLSCKASGYTFSISWINWVKQRPGQGLEWIGNIYPSGGYTNYNQKFKDKATLTVDKSSNTAYIQLSSPTSEDSAVYYCTRGYGHLDYWGQGTTLTVSA | SYELTQPPSVSVSPGQTASITCSGDKLGDKYASWYQQKPGQSPVLVIYQDNKRPSGIPERFSGSNSGNTATLTISGTQAMDEADYYCQAWDSSTAVFGGGTKLTVL |
| T2-L2-2 | C33S_L | QVQLVQSGAEVKKPGASVKLSCKASGYTFTAYYINWVRQAPGQGLEWIGRIYPGSGYTSYAQKFQGRATLTADESTSTAYMELSSLRSEDTAVYFCARPPVYYDSAWFAYWGQGTLVTVSS | SYELTQPPSVSVSPGQTASITCSGDKLGDKYASWYQQKPGQSPVLVIYQDNKRPSGIPERFSGSNSGNTATLTISGTQAMDEADYYCQAWDSSTAVFGGGTKLTVL |
| T2-I4-3 | Y31F_L | EVQLLESGPGLLKPSETLSLTCTVSGGSMINYYWSWIRQPPGERPQWLGHIIYGGTTKYNPSLESRITISRDISKNQFSLRLNSVTAADTAIYYCARVAIGVSGFLNYYYYMDVWGSGTAVTVSS | SYELTQPPSVSVSPGQTASITCSGDKLGDKFACWYQQKPGQSPVLVIYQDNKRPSGIPERFSGSNSGNTATLTISGTQAMDEADYYCQAWDSSTAVFGGGTKLTVL |
| T2-L2-3 | Y31F_L | QVQLVQSGAEVKKPGASVKLSCKASGYTFTAYYINWVRQAPGQGLEWIGRIYPGSGYTSYAQKFQGRAT | SYELTQPPSVSVSPGQTASITCSGDKLGDKFACWYQQKPGQSPVLVIYQDNKRPSGIPERF |

| | | | |
|---|---|---|---|
| | | LTADESTSTAYMELSSLRSEDTAVYFCARPPVYYDS AWFAYWGQGTLVTVSS | SGSNSGNTATLTISGTQAMDEADYYCQAW DSSTAVFGGGTKLTVL |
| T2-I4 -4 | Y31F _L; C33A _L | EVQLLESGPGLLKPSETLSLTCTVSGGSMINYYWS WIRQPPGERPQWLGHIIYGGTTKYNPSLESRITISR DISKNQFSLRLNSVTAADTAIYYCARVAIGVSGFLNY YYYMDVWGSGTAVTVSS | SYELTQPPSVSVSPGQTASITCSGDKLGDKF AAWYQQKPGQSPVLVIYQDNKRPSGIPERF SGSNSGNTATLTISGTQAMDEADYYCQAW DSSTAVFGGGTKLTVL |
| T2-L 2-4 | Y31F _L; C33A _L | QVQLVQSGAEVKKPGASVKLSCKASGYTFTAYYIN WVRQAPGQGLEWIGRIYPGSGYTSYAQKFQGRAT LTADESTSTAYMELSSLRSEDTAVYFCARPPVYYDS AWFAYWGQGTLVTVSS | SYELTQPPSVSVSPGQTASITCSGDKLGDKF AAWYQQKPGQSPVLVIYQDNKRPSGIPERF SGSNSGNTATLTISGTQAMDEADYYCQAW DSSTAVFGGGTKLTVL |
| T2-I4 -5 | Y31F _L; C33S _L | EVQLLESGPGLLKPSETLSLTCTVSGGSMINYYWS WIRQPPGERPQWLGHIIYGGTTKYNPSLESRITISR DISKNQFSLRLNSVTAADTAIYYCARVAIGVSGFLNY YYYMDVWGSGTAVTVSS | SYELTQPPSVSVSPGQTASITCSGDKLGDKF ASWYQQKPGQSPVLVIYQDNKRPSGIPERF SGSNSGNTATLTISGTQAMDEADYYCQAW DSSTAVFGGGTKLTVL |
| T2-L 2-5 | Y31F _L; C33S _L | QVQLVQSGAEVKKPGASVKLSCKASGYTFTAYYIN WVRQAPGQGLEWIGRIYPGSGYTSYAQKFQGRAT LTADESTSTAYMELSSLRSEDTAVYFCARPPVYYDS AWFAYWGQGTLVTVSS | SYELTQPPSVSVSPGQTASITCSGDKLGDKF ASWYQQKPGQSPVLVIYQDNKRPSGIPERF SGSNSGNTATLTISGTQAMDEADYYCQAW DSSTAVFGGGTKLTVL |
| T2-K 10-1 | Y94S _L | QVQLQESGAELVRPGASVKLSCKASGYTFSISWIN WVKQRPGQGLEWIGNIYPSGGYTNYNQKFKDKAT LTVDKSSNTAYIQLSSPTSEDSAVYYCTRGYGHLDY WGQGTTLTVSA | VIQMTQSPSTLSASVGDRVTITCRASQSVST WLAWYQQKPGQGPKLLIYEASSLESGVPS RFSGSGSGTEFTLTISSLQPDDFATYYCQQY NSSSFWTFGQGTKVEIK |
| T2-M 13-1 | Y94S _L | QMQLVESGGGVVQPGRSLRLSCAASGFTFRTYG MHWVRQAPGKGLEWVAVIWYDGSNKHYADSVKG RFTITRDNSKNTLNLQMNSLRAEDTAVYYCARAPQ WELVHEAFDIWGQGTMVTVSS | VIQMTQSPSTLSASVGDRVTITCRASQSVST WLAWYQQKPGQGPKLLIYEASSLESGVPS RFSGSGSGTEFTLTISSLQPDDFATYYCQQY NSSSFWTFGQGTKVEIK |
| T2-M 14-1 | S50D _L | QMQLVESGGGVVQPGRSLRLSCAASGFTFRTYG MHWVRQAPGKGLEWVAVIWYDGSNKHYADSVKG RFTITRDNSKNTLNLQMNSLRAEDTAVYYCARAPQ WELVHEAFDIWGQGTMVTVSS | DIQMTQSPSSLSASVGDRVTITCRASQSVS SAVAWYQQKPGKAPKLLIYDASSLYSGVPS RFSGSRSGTDFTLTISSLQPEDFATYYCQQS WSAYPFTFGQGTKVEIK |
| T2-H 2-1 | S50D _L | QVQLQESGPGLVKPSETLSLTCTVSGSSLTSYGVH WVRQPPGKGLEGLGVIWPGGSTNYNSALMSRVTI SKDNSKSQVSLKMSSLTAADTAVYYCARVTGTWYF DVWGQGTTVTVSS | DIQMTQSPSSLSASVGDRVTITCRASQSVS SAVAWYQQKPGKAPKLLIYDASSLYSGVPS RFSGSRSGTDFTLTISSLQPEDFATYYCQQS WSAYPFTFGQGTKVEIK |
| T2-I5 -1 | W92 S_L | EVQLLESGPGLLKPSETLSLTCTVSGGSMINYYWS WIRQPPGERPQWLGHIIYGGTTKYNPSLESRITISR DISKNQFSLRLNSVTAADTAIYYCARVAIGVSGFLNY YYYMDVWGSGTAVTVSS | DIQMTQSPSSLSASVGDRVTITCRASQSVS SAVAWYQQKPGKAPKLLIYSASSLYSGVPS RFSGSRSGTDFTLTISSLQPEDFATYYCQQS SSAYPFTFGQGTKVEIK |
| T2-M 14-2 | W92 S_L | QMQLVESGGGVVQPGRSLRLSCAASGFTFRTYG MHWVRQAPGKGLEWVAVIWYDGSNKHYADSVKG RFTITRDNSKNTLNLQMNSLRAEDTAVYYCARAPQ WELVHEAFDIWGQGTMVTVSS | DIQMTQSPSSLSASVGDRVTITCRASQSVS SAVAWYQQKPGKAPKLLIYSASSLYSGVPS RFSGSRSGTDFTLTISSLQPEDFATYYCQQS SSAYPFTFGQGTKVEIK |
| T2-H 2-2 | W92 S_L | QVQLQESGPGLVKPSETLSLTCTVSGSSLTSYGVH WVRQPPGKGLEGLGVIWPGGSTNYNSALMSRVTI SKDNSKSQVSLKMSSLTAADTAVYYCARVTGTWYF DVWGQGTTVTVSS | DIQMTQSPSSLSASVGDRVTITCRASQSVS SAVAWYQQKPGKAPKLLIYSASSLYSGVPS RFSGSRSGTDFTLTISSLQPEDFATYYCQQS SSAYPFTFGQGTKVEIK |
| T2-K 11-1 | L96H _L | QVQLQESGAELVRPGASVKLSCKASGYTFSISWIN WVKQRPGQGLEWIGNIYPSGGYTNYNQKFKDKAT LTVDKSSNTAYIQLSSPTSEDSAVYYCTRGYGHLDY WGQGTTLTVSA | DIQMTQSPSSLSASVGDRVTITCRASQSVS SAVAWYQQKPGKAPKLLIYSASSLYSGVPS RFSGSRSGTDFTLTISSLQPEDFATYYCQQS EWGGHITFGQGTKVEIK |
| T2-M 15-1 | L96H _L | QMQLVESGGGVVQPGRSLRLSCAASGFTFRTYG MHWVRQAPGKGLEWVAVIWYDGSNKHYADSVKG RFTITRDNSKNTLNLQMNSLRAEDTAVYYCARAPQ WELVHEAFDIWGQGTMVTVSS | DIQMTQSPSSLSASVGDRVTITCRASQSVS SAVAWYQQKPGKAPKLLIYSASSLYSGVPS RFSGSRSGTDFTLTISSLQPEDFATYYCQQS EWGGHITFGQGTKVEIK |
| T2-K 11-2 | W93 S_L | QVQLQESGAELVRPGASVKLSCKASGYTFSISWIN WVKQRPGQGLEWIGNIYPSGGYTNYNQKFKDKAT | DIQMTQSPSSLSASVGDRVTITCRASQSVS SAVAWYQQKPGKAPKLLIYSASSLYSGVPS |

| | | | |
|---|---|---|---|
| | | LTVDKSSNTAYIQLSSPTSEDSAVYYCTRGYGHLDY<br>WGQGTTLTVSA | RFSGSRSGTDFTLTISSLQPEDFATYYCQQS<br>ESGGLITFGQGTKVEIK |
| T2-M<br>15-2 | W93<br>S_L | QMQLVESGGGVVQPGRSLRLSCAASGFTFRTYG<br>MHWVRQAPGKGLEWVAVIWYDGSNKHYADSVKG<br>RFTITRDNSKNTLNLQMNSLRAEDTAVYYCARAPQ<br>WELVHEAFDIWGQGTMVTVSS | DIQMTQSPSSLSASVGDRVTITCRASQSVS<br>SAVAWYQQKPGKAPKLLIYSASSLYSGVPS<br>RFSGSRSGTDFTLTISSLQPEDFATYYCQQS<br>ESGGLITFGQGTKVEIK |
| T2-M<br>14-3 | S50D<br>_L;<br>A32Y<br>_L | QMQLVESGGGVVQPGRSLRLSCAASGFTFRTYG<br>MHWVRQAPGKGLEWVAVIWYDGSNKHYADSVKG<br>RFTITRDNSKNTLNLQMNSLRAEDTAVYYCARAPQ<br>WELVHEAFDIWGQGTMVTVSS | DIQMTQSPSSLSASVGDRVTITCRASQSVS<br>SYVAWYQQKPGKAPKLLIYDASSLYSGVPS<br>RFSGSRSGTDFTLTISSLQPEDFATYYCQQS<br>WSAYPFTFGQGTKVEIK |
| T2-H<br>2-3 | S50D<br>_L;<br>A32Y<br>_L | QVQLQESGPGLVKPSETLSLTCTVSGSSLTSYGVH<br>WVRQPPGKGLEGLGVIWPGGSTNYNSALMSRVTI<br>SKDNSKSQVSLKMSSLTAADTAVYYCARVTGTWYF<br>DVWGQGTTVTVSS | DIQMTQSPSSLSASVGDRVTITCRASQSVS<br>SYVAWYQQKPGKAPKLLIYDASSLYSGVPS<br>RFSGSRSGTDFTLTISSLQPEDFATYYCQQS<br>WSAYPFTFGQGTKVEIK |
| T2-M<br>14-4 | S50D<br>_L;<br>A32Y<br>_L;<br>W92<br>S_L | QMQLVESGGGVVQPGRSLRLSCAASGFTFRTYG<br>MHWVRQAPGKGLEWVAVIWYDGSNKHYADSVKG<br>RFTITRDNSKNTLNLQMNSLRAEDTAVYYCARAPQ<br>WELVHEAFDIWGQGTMVTVSS | DIQMTQSPSSLSASVGDRVTITCRASQSVS<br>SYVAWYQQKPGKAPKLLIYDASSLYSGVPS<br>RFSGSRSGTDFTLTISSLQPEDFATYYCQQS<br>SSAYPFTFGQGTKVEIK |
| T2-H<br>2-4 | S50D<br>_L;<br>A32Y<br>_L;<br>W92<br>S_L | QVQLQESGPGLVKPSETLSLTCTVSGSSLTSYGVH<br>WVRQPPGKGLEGLGVIWPGGSTNYNSALMSRVTI<br>SKDNSKSQVSLKMSSLTAADTAVYYCARVTGTWYF<br>DVWGQGTTVTVSS | DIQMTQSPSSLSASVGDRVTITCRASQSVS<br>SYVAWYQQKPGKAPKLLIYDASSLYSGVPS<br>RFSGSRSGTDFTLTISSLQPEDFATYYCQQS<br>SSAYPFTFGQGTKVEIK |
| T2-M<br>14-5 | S50D<br>_L;<br>W92<br>S_L | QMQLVESGGGVVQPGRSLRLSCAASGFTFRTYG<br>MHWVRQAPGKGLEWVAVIWYDGSNKHYADSVKG<br>RFTITRDNSKNTLNLQMNSLRAEDTAVYYCARAPQ<br>WELVHEAFDIWGQGTMVTVSS | DIQMTQSPSSLSASVGDRVTITCRASQSVS<br>SAVAWYQQKPGKAPKLLIYDASSLYSGVPS<br>RFSGSRSGTDFTLTISSLQPEDFATYYCQQS<br>SSAYPFTFGQGTKVEIK |
| T2-H<br>2-5 | S50D<br>_L;<br>W92<br>S_L | QVQLQESGPGLVKPSETLSLTCTVSGSSLTSYGVH<br>WVRQPPGKGLEGLGVIWPGGSTNYNSALMSRVTI<br>SKDNSKSQVSLKMSSLTAADTAVYYCARVTGTWYF<br>DVWGQGTTVTVSS | DIQMTQSPSSLSASVGDRVTITCRASQSVS<br>SAVAWYQQKPGKAPKLLIYDASSLYSGVPS<br>RFSGSRSGTDFTLTISSLQPEDFATYYCQQS<br>SSAYPFTFGQGTKVEIK |
| T2-K<br>11-3 | W93<br>S_L;<br>L96H<br>_L | QVQLQESGAELVRPGASVKLSCKASGYTFSISWIN<br>WVKQRPGQGLEWIGNIYPSGGYTNYNQKFKDKAT<br>LTVDKSSNTAYIQLSSPTSEDSAVYYCTRGYGHLDY<br>WGQGTTLTVSA | DIQMTQSPSSLSASVGDRVTITCRASQSVS<br>SAVAWYQQKPGKAPKLLIYSASSLYSGVPS<br>RFSGSRSGTDFTLTISSLQPEDFATYYCQQS<br>ESGGHITFGQGTKVEIK |
| T2-M<br>15-3 | W93<br>S_L;<br>L96H<br>_L | QMQLVESGGGVVQPGRSLRLSCAASGFTFRTYG<br>MHWVRQAPGKGLEWVAVIWYDGSNKHYADSVKG<br>RFTITRDNSKNTLNLQMNSLRAEDTAVYYCARAPQ<br>WELVHEAFDIWGQGTMVTVSS | DIQMTQSPSSLSASVGDRVTITCRASQSVS<br>SAVAWYQQKPGKAPKLLIYSASSLYSGVPS<br>RFSGSRSGTDFTLTISSLQPEDFATYYCQQS<br>ESGGHITFGQGTKVEIK |
| T2-K<br>1-7 | F95H<br>_L | QVQLQESGAELVRPGASVKLSCKASGYTFSISWIN<br>WVKQRPGQGLEWIGNIYPSGGYTNYNQKFKDKAT<br>LTVDKSSNTAYIQLSSPTSEDSAVYYCTRGYGHLDY<br>WGQGTTLTVSA | QSVLTQPPSVSVAPGQTARISCSGDNIGSYY<br>VHWYQQKPGQAPVLVIYEDSERPSGIPERF<br>SGSNSGNTATLTISGTQAEDEADYYCSSYD<br>DPNHQVFGGGTKLTVL |
| T2-M<br>1-7 | F95H<br>_L | QMQLVESGGGVVQPGRSLRLSCAASGFTFRTYG<br>MHWVRQAPGKGLEWVAVIWYDGSNKHYADSVKG<br>RFTITRDNSKNTLNLQMNSLRAEDTAVYYCARAPQ<br>WELVHEAFDIWGQGTMVTVSS | QSVLTQPPSVSVAPGQTARISCSGDNIGSYY<br>VHWYQQKPGQAPVLVIYEDSERPSGIPERF<br>SGSNSGNTATLTISGTQAEDEADYYCSSYD<br>DPNHQVFGGGTKLTVL |
| T2-K<br>1-8 | S55A<br>_L | QVQLQESGAELVRPGASVKLSCKASGYTFSISWIN<br>WVKQRPGQGLEWIGNIYPSGGYTNYNQKFKDKAT<br>LTVDKSSNTAYIQLSSPTSEDSAVYYCTRGYGHLDY<br>WGQGTTLTVSA | QSVLTQPPSVSVAPGQTARISCSGDNIGSYY<br>VHWYQQKPGQAPVLVIYEDSERPAGIPERF<br>SGSNSGNTATLTISGTQAEDEADYYCSSYD<br>DPNFQVFGGGTKLTVL |
| T2-M<br>1-8 | S55A<br>_L | QMQLVESGGGVVQPGRSLRLSCAASGFTFRTYG<br>MHWVRQAPGKGLEWVAVIWYDGSNKHYADSVKG | QSVLTQPPSVSVAPGQTARISCSGDNIGSYY<br>VHWYQQKPGQAPVLVIYEDSERPAGIPERF |

| | | | |
|---|---|---|---|
| | | RFTITRDNSKNTLNLQMNSLRAEDTAVYYCARAPQ WELVHEAFDIWGQGTMVTVSS | SGSNSGNTATLTISGTQAEDEADYYCSSYD DPNFQVFGGGTKLTVL |
| T2-K 1-9 | S55E _L | QVQLQESGAELVRPGASVKLSCKASGYTFSISWIN WVKQRPGQGLEWIGNIYPSGGYTNYNQKFKDKAT LTVDKSSNTAYIQLSSPTSEDSAVYYCTRGYGHLDY WGQGTTLTVSA | QSVLTQPPSVSVAPGQTARISCSGDNIGSYY VHWYQQKPGQAPVLVIYEDSERPEGIPERF SGSNSGNTATLTISGTQAEDEADYYCSSYD DPNFQVFGGGTKLTVL |
| T2-M 1-9 | S55E _L | QMQLVESGGGVVQPGRSLRLSCAASGFTFRTYG MHWVRQAPGKGLEWVAVIWYDGSNKHYADSVKG RFTITRDNSKNTLNLQMNSLRAEDTAVYYCARAPQ WELVHEAFDIWGQGTMVTVSS | QSVLTQPPSVSVAPGQTARISCSGDNIGSYY VHWYQQKPGQAPVLVIYEDSERPEGIPERF SGSNSGNTATLTISGTQAEDEADYYCSSYD DPNFQVFGGGTKLTVL |
| T2-K 1-10 | S55E _L; F95H _L | QVQLQESGAELVRPGASVKLSCKASGYTFSISWIN WVKQRPGQGLEWIGNIYPSGGYTNYNQKFKDKAT LTVDKSSNTAYIQLSSPTSEDSAVYYCTRGYGHLDY WGQGTTLTVSA | QSVLTQPPSVSVAPGQTARISCSGDNIGSYY VHWYQQKPGQAPVLVIYEDSERPEGIPERF SGSNSGNTATLTISGTQAEDEADYYCSSYD DPNHQVFGGGTKLTVL |
| T2-M 1-10 | S55E _L; F95H _L | QMQLVESGGGVVQPGRSLRLSCAASGFTFRTYG MHWVRQAPGKGLEWVAVIWYDGSNKHYADSVKG RFTITRDNSKNTLNLQMNSLRAEDTAVYYCARAPQ WELVHEAFDIWGQGTMVTVSS | QSVLTQPPSVSVAPGQTARISCSGDNIGSYY VHWYQQKPGQAPVLVIYEDSERPEGIPERF SGSNSGNTATLTISGTQAEDEADYYCSSYD DPNHQVFGGGTKLTVL |
| T2-K 1-11 | S55E _L; Y30K _L | QVQLQESGAELVRPGASVKLSCKASGYTFSISWIN WVKQRPGQGLEWIGNIYPSGGYTNYNQKFKDKAT LTVDKSSNTAYIQLSSPTSEDSAVYYCTRGYGHLDY WGQGTTLTVSA | QSVLTQPPSVSVAPGQTARISCSGDNIGSKY VHWYQQKPGQAPVLVIYEDSERPEGIPERF SGSNSGNTATLTISGTQAEDEADYYCSSYD DPNFQVFGGGTKLTVL |
| T2-M 1-11 | S55E _L; Y30K _L | QMQLVESGGGVVQPGRSLRLSCAASGFTFRTYG MHWVRQAPGKGLEWVAVIWYDGSNKHYADSVKG RFTITRDNSKNTLNLQMNSLRAEDTAVYYCARAPQ WELVHEAFDIWGQGTMVTVSS | QSVLTQPPSVSVAPGQTARISCSGDNIGSKY VHWYQQKPGQAPVLVIYEDSERPEGIPERF SGSNSGNTATLTISGTQAEDEADYYCSSYD DPNFQVFGGGTKLTVL |
| T2-K 1-12 | S55T _L | QVQLQESGAELVRPGASVKLSCKASGYTFSISWIN WVKQRPGQGLEWIGNIYPSGGYTNYNQKFKDKAT LTVDKSSNTAYIQLSSPTSEDSAVYYCTRGYGHLDY WGQGTTLTVSA | QSVLTQPPSVSVAPGQTARISCSGDNIGSYY VHWYQQKPGQAPVLVIYEDSERPTGIPERF SGSNSGNTATLTISGTQAEDEADYYCSSYD DPNFQVFGGGTKLTVL |
| T2-M 1-12 | S55T _L | QMQLVESGGGVVQPGRSLRLSCAASGFTFRTYG MHWVRQAPGKGLEWVAVIWYDGSNKHYADSVKG RFTITRDNSKNTLNLQMNSLRAEDTAVYYCARAPQ WELVHEAFDIWGQGTMVTVSS | QSVLTQPPSVSVAPGQTARISCSGDNIGSYY VHWYQQKPGQAPVLVIYEDSERPTGIPERF SGSNSGNTATLTISGTQAEDEADYYCSSYD DPNFQVFGGGTKLTVL |
| T2-M 5-1 | D93S _L; R94T _L; L97T _L; T98F _L | QMQLVESGGGVVQPGRSLRLSCAASGFTFRTYG MHWVRQAPGKGLEWVAVIWYDGSNKHYADSVKG RFTITRDNSKNTLNLQMNSLRAEDTAVYYCARAPQ WELVHEAFDIWGQGTMVTVSS | ETVLTQSPGTLTLSPGERATLTCRASQSVYT YLAWYQEKPGQAPRLLIYGASSRATGIPDRF SGSGSGTEFTLTISSLQSEDFAVYYCQQYYS TPPTFFGGGTKVEIK |
| T2-L 1-1 | D93S _L; R94T _L; L97T _L; T98F _L | QVQLVQSGAEVKKPGASVKLSCKASGYTFTAYYIN WVRQAPGQGLEWIGRIYPGSGYTSYAQKFQGRAT LTADESTSTAYMELSSLRSEDTAVYFCARPPVYYDS AWFAYWGQGTLVTVSS | ETVLTQSPGTLTLSPGERATLTCRASQSVYT YLAWYQEKPGQAPRLLIYGASSRATGIPDRF SGSGSGTEFTLTISSLQSEDFAVYYCQQYYS TPPTFFGGGTKVEIK |
| T2-M 5-2 | L97d el_L | QMQLVESGGGVVQPGRSLRLSCAASGFTFRTYG MHWVRQAPGKGLEWVAVIWYDGSNKHYADSVKG RFTITRDNSKNTLNLQMNSLRAEDTAVYYCARAPQ WELVHEAFDIWGQGTMVTVSS | ETVLTQSPGTLTLSPGERATLTCRASQSVYT YLAWYQEKPGQAPRLLIYGASSRATGIPDRF SGSGSGTEFTLTISSLQSEDFAVYYCQQYYD RPPTFGGGTKVEIK |
| T2-L 1-2 | L97d el_L | QVQLVQSGAEVKKPGASVKLSCKASGYTFTAYYIN WVRQAPGQGLEWIGRIYPGSGYTSYAQKFQGRAT | ETVLTQSPGTLTLSPGERATLTCRASQSVYT YLAWYQEKPGQAPRLLIYGASSRATGIPDRF |

| | | | |
|---|---|---|---|
| | | LTADESTSTAYMELSSLRSEDTAVYFCARPPVYYDS AWFAYWGQGTLVTVSS | SGSGSGTEFTLTISSLQSEDFAVYYCQQYYD RPPTFGGGTKVEIK |
| T2-M 5-3 | R94T _L; Y30S _L; D93S _L; P95S _L | QMQLVESGGGVVQPGRSLRLSCAASGFTFRTYG MHWVRQAPGKGLEWVAVIWYDGSNKHYADSVKG RFTITRDNSKNTLNLQMNSLRAEDTAVYYCARAPQ WELVHEAFDIWGQGTMVTVSS | ETVLTQSPGTLTLSPGERATLTCRASQSVST YLAWYQEKPGQAPRLLIYGASSRATGIPDRF SGSGSGTEFTLTISSLQSEDFAVYYCQQYYS TSPLTFGGGTKVEIK |
| T2-L 1-3 | R94T _L; Y30S _L; D93S _L; P95S _L | QVQLVQSGAEVKKPGASVKLSCKASGYTFTAYYIN WVRQAPGQGLEWIGRIYPGSGYTSYAQKFQGRAT LTADESTSTAYMELSSLRSEDTAVYFCARPPVYYDS AWFAYWGQGTLVTVSS | ETVLTQSPGTLTLSPGERATLTCRASQSVST YLAWYQEKPGQAPRLLIYGASSRATGIPDRF SGSGSGTEFTLTISSLQSEDFAVYYCQQYYS TSPLTFGGGTKVEIK |
| T2-M 5-4 | R94T _L; Y30S _L; P95S _L | QMQLVESGGGVVQPGRSLRLSCAASGFTFRTYG MHWVRQAPGKGLEWVAVIWYDGSNKHYADSVKG RFTITRDNSKNTLNLQMNSLRAEDTAVYYCARAPQ WELVHEAFDIWGQGTMVTVSS | ETVLTQSPGTLTLSPGERATLTCRASQSVST YLAWYQEKPGQAPRLLIYGASSRATGIPDRF SGSGSGTEFTLTISSLQSEDFAVYYCQQYYD TSPLTFGGGTKVEIK |
| T2-L 1-4 | R94T _L; Y30S _L; P95S _L | QVQLVQSGAEVKKPGASVKLSCKASGYTFTAYYIN WVRQAPGQGLEWIGRIYPGSGYTSYAQKFQGRAT LTADESTSTAYMELSSLRSEDTAVYFCARPPVYYDS AWFAYWGQGTLVTVSS | ETVLTQSPGTLTLSPGERATLTCRASQSVST YLAWYQEKPGQAPRLLIYGASSRATGIPDRF SGSGSGTEFTLTISSLQSEDFAVYYCQQYYD TSPLTFGGGTKVEIK |
| T2-M 5-5 | R94T _L; Y30S _L; Y92S _L; P95S _L | QMQLVESGGGVVQPGRSLRLSCAASGFTFRTYG MHWVRQAPGKGLEWVAVIWYDGSNKHYADSVKG RFTITRDNSKNTLNLQMNSLRAEDTAVYYCARAPQ WELVHEAFDIWGQGTMVTVSS | ETVLTQSPGTLTLSPGERATLTCRASQSVST YLAWYQEKPGQAPRLLIYGASSRATGIPDRF SGSGSGTEFTLTISSLQSEDFAVYYCQQYSD TSPLTFGGGTKVEIK |
| T2-L 1-5 | R94T _L; Y30S _L; Y92S _L; P95S _L | QVQLVQSGAEVKKPGASVKLSCKASGYTFTAYYIN WVRQAPGQGLEWIGRIYPGSGYTSYAQKFQGRAT LTADESTSTAYMELSSLRSEDTAVYFCARPPVYYDS AWFAYWGQGTLVTVSS | ETVLTQSPGTLTLSPGERATLTCRASQSVST YLAWYQEKPGQAPRLLIYGASSRATGIPDRF SGSGSGTEFTLTISSLQSEDFAVYYCQQYSD TSPLTFGGGTKVEIK |
| T2-M 5-6 | Y30S _L | QMQLVESGGGVVQPGRSLRLSCAASGFTFRTYG MHWVRQAPGKGLEWVAVIWYDGSNKHYADSVKG RFTITRDNSKNTLNLQMNSLRAEDTAVYYCARAPQ WELVHEAFDIWGQGTMVTVSS | ETVLTQSPGTLTLSPGERATLTCRASQSVST YLAWYQEKPGQAPRLLIYGASSRATGIPDRF SGSGSGTEFTLTISSLQSEDFAVYYCQQYYD RPPLTFGGGTKVEIK |
| T2-L 1-6 | Y30S _L | QVQLVQSGAEVKKPGASVKLSCKASGYTFTAYYIN WVRQAPGQGLEWIGRIYPGSGYTSYAQKFQGRAT LTADESTSTAYMELSSLRSEDTAVYFCARPPVYYDS AWFAYWGQGTLVTVSS | ETVLTQSPGTLTLSPGERATLTCRASQSVST YLAWYQEKPGQAPRLLIYGASSRATGIPDRF SGSGSGTEFTLTISSLQSEDFAVYYCQQYYD RPPLTFGGGTKVEIK |
| T2-M 5-7 | Y30S _L; | QMQLVESGGGVVQPGRSLRLSCAASGFTFRTYG MHWVRQAPGKGLEWVAVIWYDGSNKHYADSVKG | ETVLTQSPGTLTLSPGERATLTCRASQSVST YLAWYQEKPGQAPRLLIYGASSRATGIPDRF |

| | | | |
|---|---|---|---|
| | D93S_L | RFTITRDNSKNTLNLQMNSLRAEDTAVYYCARAPQWELVHEAFDIWGQGTMVTVSS | SGSGSGTEFTLTISSLQSEDFAVYYCQQYYSRPPLTFGGGTKVEIK |
| T2-L1-7 | Y30S_L; D93S_L | QVQLVQSGAEVKKPGASVKLSCKASGYTFTAYYINWVRQAPGQGLEWIGRIYPGSGYTSYAQKFQGRATLTADESTSTAYMELSSLRSEDTAVYFCARPPVYYDSAWFAYWGQGTLVTVSS | ETVLTQSPGTLTLSPGERATLTCRASQSVSTYLAWYQEKPGQAPRLLIYGASSRATGIPDRFSGSGSGTEFTLTISSLQSEDFAVYYCQQYYSRPPLTFGGGTKVEIK |
| T2-M5-8 | Y30S_L; D93S_L; L97T_L | QMQLVESGGGVVQPGRSLRLSCAASGFTFRTYGMHWVRQAPGKGLEWVAVIWYDGSNKHYADSVKGRFTITRDNSKNTLNLQMNSLRAEDTAVYYCARAPQWELVHEAFDIWGQGTMVTVSS | ETVLTQSPGTLTLSPGERATLTCRASQSVSTYLAWYQEKPGQAPRLLIYGASSRATGIPDRFSGSGSGTEFTLTISSLQSEDFAVYYCQQYYSRPPTTFGGGTKVEIK |
| T2-L1-8 | Y30S_L; D93S_L; L97T_L | QVQLVQSGAEVKKPGASVKLSCKASGYTFTAYYINWVRQAPGQGLEWIGRIYPGSGYTSYAQKFQGRATLTADESTSTAYMELSSLRSEDTAVYFCARPPVYYDSAWFAYWGQGTLVTVSS | ETVLTQSPGTLTLSPGERATLTCRASQSVSTYLAWYQEKPGQAPRLLIYGASSRATGIPDRFSGSGSGTEFTLTISSLQSEDFAVYYCQQYYSRPPTTFGGGTKVEIK |
| T2-M5-9 | Y30S_L; D93S_L; P95S_L | QMQLVESGGGVVQPGRSLRLSCAASGFTFRTYGMHWVRQAPGKGLEWVAVIWYDGSNKHYADSVKGRFTITRDNSKNTLNLQMNSLRAEDTAVYYCARAPQWELVHEAFDIWGQGTMVTVSS | ETVLTQSPGTLTLSPGERATLTCRASQSVSTYLAWYQEKPGQAPRLLIYGASSRATGIPDRFSGSGSGTEFTLTISSLQSEDFAVYYCQQYYSRSPLTFGGGTKVEIK |
| T2-L1-9 | Y30S_L; D93S_L; P95S_L | QVQLVQSGAEVKKPGASVKLSCKASGYTFTAYYINWVRQAPGQGLEWIGRIYPGSGYTSYAQKFQGRATLTADESTSTAYMELSSLRSEDTAVYFCARPPVYYDSAWFAYWGQGTLVTVSS | ETVLTQSPGTLTLSPGERATLTCRASQSVSTYLAWYQEKPGQAPRLLIYGASSRATGIPDRFSGSGSGTEFTLTISSLQSEDFAVYYCQQYYSRSPLTFGGGTKVEIK |
| T2-M5-10 | Y30S_L; D93S_L; Y92S_L | QMQLVESGGGVVQPGRSLRLSCAASGFTFRTYGMHWVRQAPGKGLEWVAVIWYDGSNKHYADSVKGRFTITRDNSKNTLNLQMNSLRAEDTAVYYCARAPQWELVHEAFDIWGQGTMVTVSS | ETVLTQSPGTLTLSPGERATLTCRASQSVSTYLAWYQEKPGQAPRLLIYGASSRATGIPDRFSGSGSGTEFTLTISSLQSEDFAVYYCQQYSSRPPLTFGGGTKVEIK |
| T2-L1-10 | Y30S_L; D93S_L; Y92S_L | QVQLVQSGAEVKKPGASVKLSCKASGYTFTAYYINWVRQAPGQGLEWIGRIYPGSGYTSYAQKFQGRATLTADESTSTAYMELSSLRSEDTAVYFCARPPVYYDSAWFAYWGQGTLVTVSS | ETVLTQSPGTLTLSPGERATLTCRASQSVSTYLAWYQEKPGQAPRLLIYGASSRATGIPDRFSGSGSGTEFTLTISSLQSEDFAVYYCQQYSSRPPLTFGGGTKVEIK |
| T2-M5-11 | Y30S_L; D93S_L; Y92S_L; P95S_L | QMQLVESGGGVVQPGRSLRLSCAASGFTFRTYGMHWVRQAPGKGLEWVAVIWYDGSNKHYADSVKGRFTITRDNSKNTLNLQMNSLRAEDTAVYYCARAPQWELVHEAFDIWGQGTMVTVSS | ETVLTQSPGTLTLSPGERATLTCRASQSVSTYLAWYQEKPGQAPRLLIYGASSRATGIPDRFSGSGSGTEFTLTISSLQSEDFAVYYCQQYSSRSPLTFGGGTKVEIK |
| T2-L1-11 | Y30S_L; D93S_L; Y92S_L; | QVQLVQSGAEVKKPGASVKLSCKASGYTFTAYYINWVRQAPGQGLEWIGRIYPGSGYTSYAQKFQGRATLTADESTSTAYMELSSLRSEDTAVYFCARPPVYYDSAWFAYWGQGTLVTVSS | ETVLTQSPGTLTLSPGERATLTCRASQSVSTYLAWYQEKPGQAPRLLIYGASSRATGIPDRFSGSGSGTEFTLTISSLQSEDFAVYYCQQYSSRSPLTFGGGTKVEIK |

| | | | |
|---|---|---|---|
| | P95S _L | | |
| T2-M 5-12 | Y30S _L; L97T _L | QMQLVESGGGVVQPGRSLRLSCAASGFTFRTYGMHWVRQAPGKGLEWVAVIWYDGSNKHYADSVKGRFTITRDNSKNTLNLQMNSLRAEDTAVYYCARAPQWELVHEAFDIWGQGTMVTVSS | ETVLTQSPGTLTLSPGERATLTCRASQSVSTYLAWYQEKPGQAPRLLIYGASSRATGIPDRFSGSGSGTEFTLTISSLQSEDFAVYYCQQYYDRPPTTFGGGTKVEIK |
| T2-L 1-12 | Y30S _L; L97T _L | QVQLVQSGAEVKKPGASVKLSCKASGYTFTAYYINWVRQAPGQGLEWIGRIYPGSGYTSYAQKFQGRATLTADESTSTAYMELSSLRSEDTAVYFCARPPVYYDSAWFAYWGQGTLVTVSS | ETVLTQSPGTLTLSPGERATLTCRASQSVSTYLAWYQEKPGQAPRLLIYGASSRATGIPDRFSGSGSGTEFTLTISSLQSEDFAVYYCQQYYDRPPTTFGGGTKVEIK |
| T2-M 5-13 | Y30S _L; L97T _L; Y92S _L | QMQLVESGGGVVQPGRSLRLSCAASGFTFRTYGMHWVRQAPGKGLEWVAVIWYDGSNKHYADSVKGRFTITRDNSKNTLNLQMNSLRAEDTAVYYCARAPQWELVHEAFDIWGQGTMVTVSS | ETVLTQSPGTLTLSPGERATLTCRASQSVSTYLAWYQEKPGQAPRLLIYGASSRATGIPDRFSGSGSGTEFTLTISSLQSEDFAVYYCQQYSDRPPTTFGGGTKVEIK |
| T2-L 1-13 | Y30S _L; L97T _L; Y92S _L | QVQLVQSGAEVKKPGASVKLSCKASGYTFTAYYINWVRQAPGQGLEWIGRIYPGSGYTSYAQKFQGRATLTADESTSTAYMELSSLRSEDTAVYFCARPPVYYDSAWFAYWGQGTLVTVSS | ETVLTQSPGTLTLSPGERATLTCRASQSVSTYLAWYQEKPGQAPRLLIYGASSRATGIPDRFSGSGSGTEFTLTISSLQSEDFAVYYCQQYSDRPPTTFGGGTKVEIK |
| T2-M 5-14 | Y30S _L; P95S _L | QMQLVESGGGVVQPGRSLRLSCAASGFTFRTYGMHWVRQAPGKGLEWVAVIWYDGSNKHYADSVKGRFTITRDNSKNTLNLQMNSLRAEDTAVYYCARAPQWELVHEAFDIWGQGTMVTVSS | ETVLTQSPGTLTLSPGERATLTCRASQSVSTYLAWYQEKPGQAPRLLIYGASSRATGIPDRFSGSGSGTEFTLTISSLQSEDFAVYYCQQYYDRSPLTFGGGTKVEIK |
| T2-L 1-14 | Y30S _L; P95S _L | QVQLVQSGAEVKKPGASVKLSCKASGYTFTAYYINWVRQAPGQGLEWIGRIYPGSGYTSYAQKFQGRATLTADESTSTAYMELSSLRSEDTAVYFCARPPVYYDSAWFAYWGQGTLVTVSS | ETVLTQSPGTLTLSPGERATLTCRASQSVSTYLAWYQEKPGQAPRLLIYGASSRATGIPDRFSGSGSGTEFTLTISSLQSEDFAVYYCQQYYDRSPLTFGGGTKVEIK |
| T2-M 5-15 | Y30S _L; Y92S _L | QMQLVESGGGVVQPGRSLRLSCAASGFTFRTYGMHWVRQAPGKGLEWVAVIWYDGSNKHYADSVKGRFTITRDNSKNTLNLQMNSLRAEDTAVYYCARAPQWELVHEAFDIWGQGTMVTVSS | ETVLTQSPGTLTLSPGERATLTCRASQSVSTYLAWYQEKPGQAPRLLIYGASSRATGIPDRFSGSGSGTEFTLTISSLQSEDFAVYYCQQYSDRPPLTFGGGTKVEIK |
| T2-L 1-15 | Y30S _L; Y92S _L | QVQLVQSGAEVKKPGASVKLSCKASGYTFTAYYINWVRQAPGQGLEWIGRIYPGSGYTSYAQKFQGRATLTADESTSTAYMELSSLRSEDTAVYFCARPPVYYDSAWFAYWGQGTLVTVSS | ETVLTQSPGTLTLSPGERATLTCRASQSVSTYLAWYQEKPGQAPRLLIYGASSRATGIPDRFSGSGSGTEFTLTISSLQSEDFAVYYCQQYSDRPPLTFGGGTKVEIK |
| T2-M 5-16 | Y30S _L; Y92S _L; P95S _L | QMQLVESGGGVVQPGRSLRLSCAASGFTFRTYGMHWVRQAPGKGLEWVAVIWYDGSNKHYADSVKGRFTITRDNSKNTLNLQMNSLRAEDTAVYYCARAPQWELVHEAFDIWGQGTMVTVSS | ETVLTQSPGTLTLSPGERATLTCRASQSVSTYLAWYQEKPGQAPRLLIYGASSRATGIPDRFSGSGSGTEFTLTISSLQSEDFAVYYCQQYSDRSPLTFGGGTKVEIK |
| T2-L 1-16 | Y30S _L; Y92S _L; P95S _L | QVQLVQSGAEVKKPGASVKLSCKASGYTFTAYYINWVRQAPGQGLEWIGRIYPGSGYTSYAQKFQGRATLTADESTSTAYMELSSLRSEDTAVYFCARPPVYYDSAWFAYWGQGTLVTVSS | ETVLTQSPGTLTLSPGERATLTCRASQSVSTYLAWYQEKPGQAPRLLIYGASSRATGIPDRFSGSGSGTEFTLTISSLQSEDFAVYYCQQYSDRSPLTFGGGTKVEIK |
| T2-K 5-5 | A50S _L | QVQLQESGAELVRPGASVKLSCKASGYTFSISWINWVKQRPGQGLEWIGNIYPSGGYTNYNQKFKDKATLTVDKSSNTAYIQLSSPTSEDSAVYYCTRGYGHLDYWGQGTTLTVSA | DIQMTQSPSSLSASVGDRVTITCRASQSISSYLNWYQQKPGKAPKLLIYSASSLQSGVPSRFSGSGSGTDFTLTISSLQPEDFATYYCQQSYSTLALTFGGGTKVEIK |
| T2-M 7-5 | A50S _L | QMQLVESGGGVVQPGRSLRLSCAASGFTFRTYGMHWVRQAPGKGLEWVAVIWYDGSNKHYADSVKG | DIQMTQSPSSLSASVGDRVTITCRASQSISSYLNWYQQKPGKAPKLLIYSASSLQSGVPSR |

| | | | |
|---|---|---|---|
| | | RFTITRDNSKNTLNLQMNSLRAEDTAVYYCARAPQWELVHEAFDIWGQGTMVTVSS | FSGSGSGTDFTLTISSLQPEDFATYYCQQSYSTLALTFGGGTKVEIK |
| T1-Dd | | EVKLQQSGAELVRPGSSVKISCKASGYAFSSYWMNWVKQRPGQGLEWIGQIYPGDGDTNYNGKFKGQATLTADKSSSTAYMQLSGLTSEDSAVYFCARKTISSVVDFYFDYWGQGTTVTVSS | DIELTQSPKFMSTSVGDRVSVTCKASQNVGTNVAWYQQKPGQSPKPLIYSATYRNSGVPDRFTGSGSGTDFTLTITNVQSKDLADYFCQQYNRYPYTSGGGTKLEIK |
| T1-Ee | | EVQLVQSGAEVKKPGASVKVSCKASGYKFTNYVMSWVRQAPGQRLEWMGYINPYNDAIKYNEKFTGRVTITRDTSASTAYMELSSLRSEDTAVYYCAREGDFYANYGRLGFAYWGQGTLVTVSS | DIQMTQSPSSLSASVGDRVTITCRASQDISNYLNWYQQKPGKAPKLLLYYTSRLHSGVPSRFSGSGSGTDYTLTISSLQPEDFATYFCQQGAGFPYTFGGGTKVEIK |
| T1-Bb | | EVQLVESGGGLVQPGGSLRLSCAASGYDFDNYGMNWVRQAPGKGLEWVGWINTYTGEPTYAADFKRRFTFSLDTSKSTAYLQMNSLRAEDTAVYYCAKYPHYYGSSHWYFDVWGQGTLVTVSS | DIQMTQSPSSLSASVGDRVTITCSASQDISNYLNWYQQKPGKAPKVLIYFTDDLHSGVPSRFSGSGSGTDFTLTISSLQPEDFATYYCQQYSTVPWTFGQGTKVEIK |
| T1-Cc | | QVQLVQSGAEVKKPGSSVKVSCKASGYAFSSYWMNWVRQAPGQGLEWMGQIWPGDSDTNYAQKFQGRVTITADESTSTAYMELSSLRSEDTAVYYCARRETTTVGRYYYAMDYWGQGTTVTVSS | DIQLTQSPSFLSASVGDRVTITCKASQSVDYSGDSYLNWYQQKPGKAPKLLIYDASNLVSGVPSRFSGSGSGTEFTLTISSLQPEDFATYYCQQSTENPWTFGGGTKLEIK |
| T1-Ff | | EVQLLESGGGLVQPGGSLRLSCAVSGFTFNSFAMSWVRQAPGKGLEWVSAISGSGGGTYYADSVKGRFTISRDNSKNTLYLQMNSLRAEDTAVYFCAKDKILWFGEPVFDYWGQGTLVTVSS | EIVLTQSPATLSLSPGERATLSCRASQSVSSYLAWYQQKPGQAPRLLIYDASNRATGIPARFSGSGSGTDFTLTISSLEPEDFAVYYCQQRSNWPPTFGQGTKVEIK |
| T1-Gg | | QVQLVQSGAEVKKPGASVKLSCKASGYTFTAYYINWVRQAPGQGLEWIGRIYPGSGYTSYAQKFQGRATLTADESTSTAYMELSSLRSEDTAVYFCARPPVYYDSAWFAYWGQGTLVTVSS | DIQMTQSPSSLSASVGDRVTITCRASQSISSYLNWYQQKPGKAPKLLIYAASSLQSGVPSRFSGSGSGTDFTLTISSLQPEDFATYYCQQSYSTPPTFGQGTKVEIK |
| T1-Aa | | EVQLVESGGGLVQPGGSLRLSCAASGFTISDYWIHWVRQAPGKGLEWVAGITPAGGYTYYADSVKGRFTISADTSKNTAYLQMNSLRAEDTAVYYCARFVFFLPYAMDYWGQGTLVTVSS | DIQMTQSPSSLSASVGDRVTITCRASQFLSSFGVAWYQQKPGKAPKLLIYGASSLYSGVPSRFSGSGSGTDFTLTISSLQPEDFATYYCQQGLLSPLTFGQGTKVEIK |
| T2-Ii | | EVQLLESGPGLLKPSETLSLTCTVSGGSMINYYWSWIRQPPGERPQWLGHIIYGGTTKYNPSLESRITISRDISKNQFSLRLNSVTAADTAIYYCARVAIGVSGFLNYYYYMDVWGSGTAVTVSS | ELTQSPATLSLSPGERATLSCRASQSVGRNLGWYQQKPGQAPRLLIYDASNRATGIPARFSGSGSGTDFTLTISSLEPEDFAVYYCQARLLLPQTFGQGTKVEIK |
| T2-Kk | | QVQLQESGAELVRPGASVKLSCKASGYTFSISWINWVKQRPGQGLEWIGNIYPSGGYTNYNQKFKDKATLTVDKSSNTAYIQLSSPTSEDSAVYYCTRGYGHLDYWGQGTTLTVSA | DIQLTQSPALMSASPGEKVTMTCSASSSVTFMYWYQQKPRSSPKPWIYLTSNLASGVPARFSGSGSGTSYSLTISSMEAEDAATYYCQQWSSNPYTFGGGTKLELK |
| T2-Mm | | QMQLVESGGGVVQPGRSLRLSCAASGFTFRTYGMHWVRQAPGKGLEWVAVIWYDGSNKHYADSVKGRFTITRDNSKNTLNLQMNSLRAEDTAVYYCARAPQWELVHEAFDIWGQGTMVTVSS | SYVLTQPPSVSVAPGQTARITCGGNNLGSKSVHWYQQKPGQAPVLVVYDDSDRPSWIPERFSGSNSGNTATLTISRGEAGDEADYYCQVWDSSSDHVVFGGGTKLTVL |
| T2-Ll | | QVQLVQSGAEVKKPGASVKLSCKASGYTFTAYYINWVRQAPGQGLEWIGRIYPGSGYTSYAQKFQGRATLTADESTSTAYMELSSLRSEDTAVYFCARPPVYYDSAWFAYWGQGTLVTVSS | DIQMTQSPSSLSASVGDRVTITCRASQSISSYLNWYQQKPGKAPKLLIYAASSLQSGVPSRFSGSGSGTDFTLTISSLQPEDFATYYCQQSYSTPPTFGQGTKVEIK |
| T2-Jj | | QLQLQESGPGLVKPSETLSLTCTVSGGSISSRSYYWGWIRQPPGKGLEWIGSIYYSGFTYYQPSLKSRVTISVDTSKNQFSLKLSSVTAADTAVYYCATGGPYGDYAHWFEPWGQGTLVTVSS | EIVLTQSPGTLSLSPGERATLSCRASQSVSSSYLAWYQQKPGQAPRLLIYGASSRATGIPDRFSGSGSGTDFTLTISRLEPEDFAVYYCQQYGSSPITFGQGTRLEIK |
| T2-Hh | | QVQLQESGPGLVKPSETLSLTCTVSGSSLTSYGVHWVRQPPGKGLEGLGVIWPGGSTNYNSALMSRVTISKDNSKSQVSLKMSSLTAADTAVYYCARVTGTWYFDVWGQGTTVTVSS | DIQMTQSPSSLSASLGDRVTISCSASQGISNYLNWYQQKPDGTVKLLIYYTSTLHSGVPSRFSGSGSGTDYTLTISSLQPEDIATYYCQQYSKLPWTFGGGTKLEIK |

Supplementary Table S10 — Final sequences of optimized designs.

Sequence registry for all finalized designs, providing the exact mutation strings, VH amino-acid sequences, and VL amino-acid sequences used for production and characterization.
Columns: Design ID, Mutation, VH Sequence, VL Sequence.

| Target | VH ID (PDB:Chain) | Initial VL Library | Chai-1 Confidence (>0.7) | Structure RMSD (<2Å) | Rosetta Energy Unit | AbAngle Geometry | MM/GBSA Energy | MD Stability | Final Candidates |
|---|---|---|---|---|---|---|---|---|---|
| DDR1 | 4ag4HLA | 2378 | 106 | 152 | 94 | 52 | 63 | 38 | 11 |
| TSLP | 5j13CBA | 2378 | 280 | 274 | 175 | 63 | 96 | 80 | 15 |
| Z13-IL22-2 | 3q1sHLI | 2378 | 184 | 184 | 39 | 23 | 16 | 17 | 5 |
| ERBB2 | 5o4gBAC | 2378 | 184 | 172 | 35 | 22 | 19 | 10 | 2 |
| HAVR2 | 7kqlHLT | 2378 | 219 | 73 | 40 | 20 | 19 | 1 | 1 |
| TNR9 | 8gyeEFA | 2378 | 88 | 182 | 76 | 42 | 46 | 2 | 2 |

Supplementary Table S11 — In-silico triage criteria and candidate gating.
Computational triage metrics used to prioritize designs: Chai-1 confidence (>0.7), structure RMSD (<2 Å) vs input, Rosetta energy units, AbAngle geometry, MM/GBSA energies, MD stability assessment, and the number of final gated candidates per VH/target. Thresholds reflect pass/fail gates used before synthesis.
Abbreviations: RMSD, root-mean-square deviation; MM/GBSA, molecular mechanics/generalized Born surface area; MD, molecular dynamics.

| Design ID | Format | Titer (mg/L) | Recovery (%) | SEC H.M.V. (%) | SEC L.M.V. (%)2 | SEC Main Peak Remain Time (min) | MS Target (%) | MS Mispair(%) | MS Half-Ab (%) |
|---|---|---|---|---|---|---|---|---|---|
| cLC-075 | KiH-cLC | 2782.8 | 87.0 | 2.81 | 10.62 | 8.059 | 34.0 | 2.9 | 63.1 |
| cLC-079 | KiH-cLC | 2529.0 | 89.2 | 2.68 | 0.93 | 7.903 | 95.2 | 1.4 | 3.4 |
| cLC-003 | KiH-cLC | 2603.8 | 95.6 | 4.88 | 0.0 | 8.144 | 93.7 | 4.6 | 1.7 |
| Baseline-075 | KiH-Baseline | 2482.89 | 87.7 | 1.98 | 7.21 | 8.142 | 26.3 | 27.2 | 46.5 |
| Baseline-079 | KiH-Baseline | 2529.0 | 91.9 | 5.91 | 4.82 | 7.971 | 67.8 | 32.2 | 0 |
| Baseline-003 | KiH-Baseline | 2708.95 | 90.7 | 14.23 | 29.02 | 8.434 | 33.9 | 66.1 | 0 |
| | | | | | | | | Ratio of HC1*2+LC*2 or HC2*2+LC*2or HC1+HC2+LC1*2or HC1+HC2+LC2*2 | HC1LC or HC2LC or HC1HC2 |

Supplementary Table S12 — Developability and assembly analytics for selected designs.
Manufacturing-relevant analytics including expression titer (mg/L), recovery (%), SEC high/low molecular-weight variants, main-peak retention time (min), and intact-mass outcomes (target %, mispair %, half-antibody %). These QC metrics substantiate manufacturability of optimized constructs.
Abbreviations: SEC, size-exclusion chromatography; MS, mass spectrometry; H.M.V./L.M.V., high/low molecular-weight variants.

| Target | Uniprot Entry Name | Source | Cat # | Reference Sequence (or UniuProt Entry and range) |
|---|---|---|---|---|
| TSLP | TSLP_HUMAN | ACRO Biosystems | TSP-H52Hb | Q969D9-1 (Tyr29 - Gln 159) |
| VEGFA | VEGFA_HUMAN | ACRO Biosystems | VE5-H5248 | P15692-4(Ala 27 - Arg 191) |
| DDR1 | DDR1_HUMAN | MCE | HY-P75305 | Q08345-1 (Asp21 - Ala417) |
| ERBB2 | ERBB2_HUMAN | ACRO Biosystems | HE2-H5225 | P04626-1(Thr 23 - Thr 652) |
| CD38 | CD38_HUMAN | ACRO Biosystems | CD8-H5224 | P28907-1(Val 43 - Ile 300) |
| Z13-IL22-2 | / | Inhouse | / | MAALQKSVSSFLMGTLATSCLLLLALLVQGGAAHHHHHHHHGGGGSAPISSHCRLDKSNFQQPYITNRTFMLAKEAWNWDDITDVRLIGEKLFHGVSMSERCYLMKQVLNFTLEEVLFPQSDRFQPYMQEVVPFLARLSNRLSTCHIEGDDLHIQRNVQKLKDTVKKLGESGEIKAIGELDLLFMSLRNACI |
| CD19 | CD19_HUMAN | ACRO Biosystems | CD9-H52H2 | P15391-1(Pro 20 - Lys 291) |
| CD47 | CD47_HUMAN | ACRO Biosystems | CD7-H5227 | Q08722-3(Gln 19 - Pro 139) |
| TNR9 | TNR9_HUMAN | ACRO Biosystems | 41B-H53H3 | Q07011-1(Leu 24 - Gln 186) |
| HAVR2 | HAVR2_HUMAN | ACRO Biosystems | TM3-H5229 | Q8TDQ0-1(Ser 22 - Arg 200) |

Supplementary Table S13 — Antigen materials and references.
Reagents list for antigens used in binding and characterization assays, including target names, UniProt entry names, vendors, catalog numbers, and reference sequences (or UniProt entries with residue ranges). This supports full reagent reproducibility.
Abbreviations: UniProt, Universal Protein Resource.
